# ChemCell: Chemical Tethering of Large Biomolecules to Cell Surfaces through Diels-Alder Ligation

**DOI:** 10.64898/2026.04.13.718024

**Authors:** Rastislav Dzijak, Simona Bellová, Anna Kovalová, Veronika Šlachtová, Michal Rahm, Anna Beránková, Radek Pohl, Milan Vrabel

## Abstract

Chemical engineering of cellular surfaces offers a powerful tool to endow cells with new properties and functions, yet methods for adding complex biomolecules to cell membranes remain underdeveloped. While technologies for targeted degradation of cell surface components are advancing, few approaches exist that allow efficient attachment of large biomolecules. To address this, we introduce ChemCell technology—a platform for non-genetic surface engineering of cells via metabolic installation of trans-cyclooctene (TCO) groups. These TCO groups undergo rapid and selective reactions with tetrazine-modified biomolecules, enabling the efficient tethering of diverse functional groups. This method allows for the precise attachment of proteins, peptides, enzymes, oligonucleotides, therapeutic antibodies, and large protein complexes to cell surfaces, expanding the toolbox for cellular modification and offering new possibilities for cell-based therapies and diagnostics.

## INTRODUCTION

The cell surface is an information-rich layer composed of biomolecules involved in various biological processes. Modifying the structure or composition of cell membrane components enables the modulation of cellular properties and functions. In general, these modifications can involve either the removal of specific surface components or the addition of new molecules. Both genetic and non-genetic approaches are available for these modifications.^1^ Genetic methods, such as CRISPR-Cas9 system, can achieve both the removal of a specific target and the knock-in of new proteins. Other approaches, like transfection and transduction, introduce genetic information encoding biomolecules, which are then produced by cells. One prominent example is the genetic engineering of immune cells, such as T cells, to express chimeric antigen receptors (CARs), which direct the cells to a specific antigen and trigger a cytotoxic response upon binding.^2^

However, genetic engineering presents technical challenges and safety concerns, including inconsistent viral transduction efficiency, heterogeneous expression levels, and the potential to affect endogenous genes.^3^ As a result, non-genetic methods offer an attractive, straightforward alternative.

One of the most promising strategies to modify cell surfaces is metabolic glycoengineering (MGE) This method exploits the natural biosynthetic pathways of cells to incorporate modified sugar analogs containing reactive chemical groups into glycoconjugates.^4, 5^ Over the past three decades, MGE has become a widely used method for manipulating the glycome in living organisms, offering great potential for advancing healthcare.^6^ Enzyme substrate promiscuity plays a critical role in this process, as non-natural precursors in MGE must be sufficiently small to be recognized as substrates by the enzymes.^7^ Therefore, sugar analogs bearing small functional groups, such as azides or alkynes, are typically preferred over bulkier chemical reporters.^8^ The chemical versatility of the azido group, which participates in reactions such as Staudinger ligation,^9^ metal-catalyzed^10^ and metal-free dipolar cycloadditions,^11^ has been widely employed in MGE.

To further expand the chemical repertoire and improve labeling efficiency, sugar analogs containing other bioorthogonal functional groups have been developed.^5^ Strained unsaturated moieties such as cyclopropene,^12, 13^ norbornene,^14^ bicyclononyne,^15^ dibenzylcyclooctyne (DBCO)^16^ are examples of such chemical reporters.

The strained *trans*-cyclooctenes (TCOs) are arguably the most reactive dienophiles being employed in modern chemical biology.^17^ The inverse electron demand Diels-Alder reaction (IEDDA) between TCOs and 1,2,4,5-tetrazines (Tz) exhibits reaction rates orders of magnitude higher than other bioorthogonal reactions.^18^ This extraordinary reactivity enables the use of low reagent concentrations, minimizing potential toxicity and reducing production costs.^15^

Despite their favorable properties, the strained *trans*-cyclooctenes (TCOs) are, with few notable exceptions, ^19^,^20^.^21^ rarely employed in metabolic incorporation studies. To expand the toolkit of TCO sugars available for metabolic incorporation we created a TCO derivative of N-acetylneuraminic acid (Neu5Ac).

We show here that the derivative Neu5Ac-TCO derivative is stable under biological conditions for several day, is utilized by cellular enzymatic machinery to efficiently install strained TCO groups onto cell surfaces. We show that the optimized tetrazines peptide and nucleic acid conjugates could be efficiently integrated into cell surfaces under the conditions of cell culture. Most importantly, our method offers substantially improved modification efficiency, particularly with large proteins such as enzymes and antibodies. We were able to target modified NK cells toward the cancer using antidies. We also show that the click reaction mediated by added antibody can trigger cytotoxic lysis of the target cancer cells at submicromolar concentrations. We named this technology ChemCell. Collectively the data presented here demonstrate that ChemCell enables fast and precise cell surface editing across a wide array of cell types. We believe that ChemCell will offer exciting new possibilities for non-genetic cellular or tissue engineering.

## RESULTS AND DISCUSSION

### Design of metabolic precursors and incorporation studies

Our selection of suitable sugar analogs bearing the TCO moiety was guided by several considerations. The most used derivatives for metabolic glycoengineering (MGE) are acetylated analogs of *N*-acetylmannosamine (ManNAc). Typically, small *N*-acyl substituents, such as ketones, alkynes, or azides, are well tolerated by cellular biosynthetic machinery. In contrast, larger substituents often result in reduced incorporation efficiency or may not be tolerated at all.^22^ Given that TCO is a relatively bulky substituent, we used sialic acid and derivatized it at the position 9 with the TCO. This strategy bypasses several key enzymatic steps involved in the biosynthesis of sialic acid from ManNAc. Additionally, substituents of a similar size to TCO have been successfully incorporated into this position previously.^23, 24^ It is known that TCOs differ in reactivity and stability under biological conditions, both of which are crucial factors for successful MGE.^25^ We selected the axial isomers of cyclooct-4-en (4TCO) and cyclooct-2-en (2TCO), which we anticipated would provide a balanced ratio between reactivity and stability. Based on these considerations, we synthesized two sugar derivatives **Sia-4TCO**, and **Sia-2TCO**. Sialic acids modified at position 9 were synthesized from the corresponding azide (**Sia-N_3_**)^24^ via reduction and peptide coupling (**Fig.1A**).

**Figure 1.**
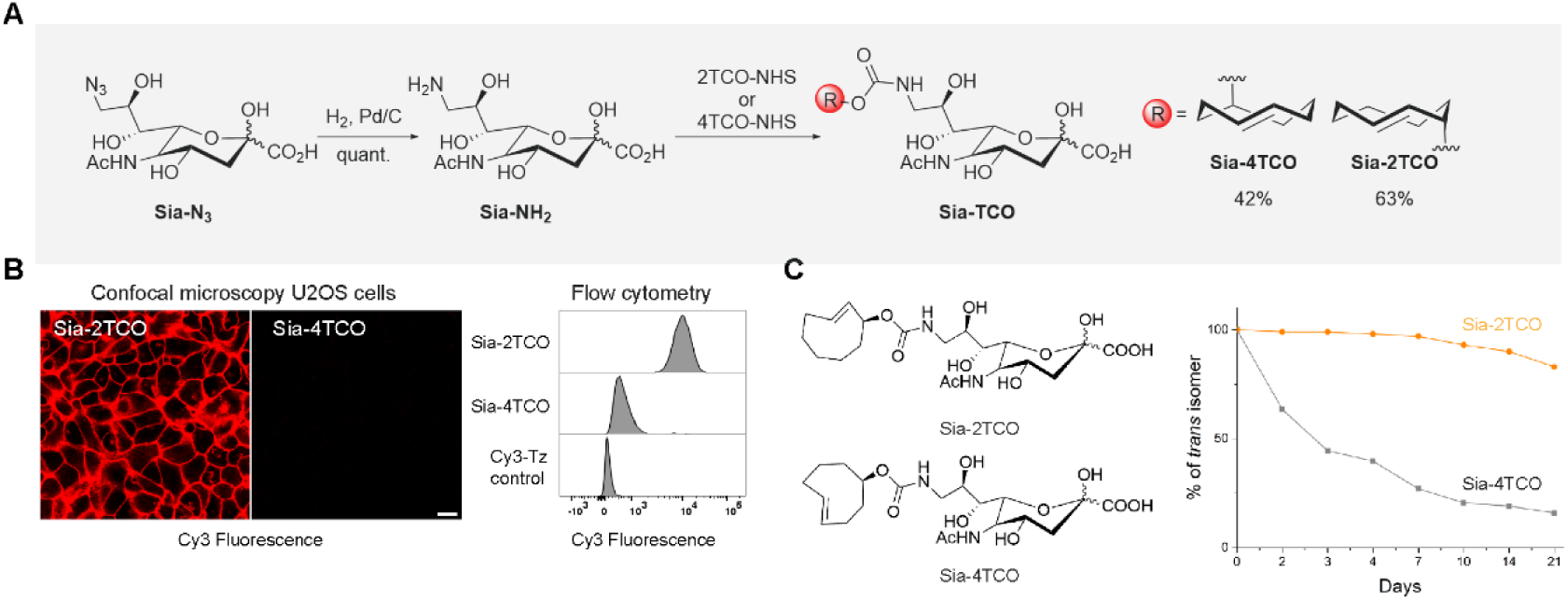
Synthesis and metabolic incorporation of TCO sugars. **A**) Synthesis o SiaTCO derivatives. Sialic acid azide (**SiaN_3_**) was reduced to amine and reacted with NHS esters of TCO. **B**) Sia-2TCO is efficiently incorporated into cellular glycoconjugates. U2OS cells were cells incubated with 1mM of each version of the SiaTCO sugar for 48h, washed and reacted with 2.5 µM of **Tz-Cy3**. Cells were immediately photographed using confocal microscope, detached from cultivation dish and analyzed using flow cytometer. **C**) Stability of each sugar was measured using NMR. Sugars were dissolved in PBS/D_2_O and measured each day over 21 days. Amount of *trans* isomer in **Sia-4TCO** sample decreased to 50% in 3 days.

To evaluate whether these precursors can be metabolically processed by living cells, we incubated the osteosarcoma cell line U2OS with 1mM concentrations of the free acids **Sia-4TCO** and **Sia-2TCO**. After 48h of incubation we visualized the incorporated sugars on cellular surface of live cells using negatively charged Sulfo-Cy3-H-tetrazine (Cy3-Tz). After staining, we imaged the cells using confocal microscopy followed by measurement of cellular fluorescence using flow cytometry. We could observe a clear signal on cells treated with **Sia-2TCO** and very faint fluorescence on cells treated with **Sia-4TCO** (**Fig.1B**) indicating that **Sia-4TCO** is inferior to **Sia-2TCO** as a metabolic precursor. Despite their identical size, there was striking difference in incorporation efficiency between **Sia-4TCO** and **Sia-2TCO**. This difference likely arises from the varying stability of the two TCO groups.^25^ Indeed, NMR measurements revealed that approximately 40% of **Sia-4TCO** isomerized to the unreactive *cis* isomer after two days in D₂O solution. In contrast, over 95% of **Sia-2TCO** remained in the *trans* conformation under the same conditions (**Fig.1C, ESI**). The isomerization of the latter was much slower, with 80% of the *trans* isomer remaining in solution even after three weeks. Under the conditions of cellular environment this process could be even faster than in NMR tube. Based on these findings, we identified **Sia-2TCO** as the only suitable metabolic precursor for the incorporation of TCO groups into glycoconjugates.

### Optimization of conditions of metabolic Sia-2TCO incorporation into cellular glycans

We used initially high concentration of sialic acid that was previously used in literature. To better understand the incorporation dynamics, we systematically tested the condition for incorporation in various cell lines. To follow the cellular fate of the sugar we used two types of tetrazine conjugates. After the incubation period with sugar-TCO cells were briefly washed to remove unincorporated sugar and then incubated with cell-impermeable sulfo-Cy3-modified tetrazine dye (**Tz-Cy3**) to react it with surface TCO-sialoglycans. After which the cells were washed and incubated with second cell-permeable, fluorogenic coumarin tetrazine dye (**Coum-Tz**). The second dye penetrates cells and becomes highly fluorescent upon reaction with the TCO^26^. Using this approach, we could follow both the metabolic incorporation of sugar into membranes and intracellular accumulation of sugar (**Fig.2A**). The incorporation of **Sia-2TCO** was found to depend on both dose (**Fig.2B**) and time (**Fig2C**). Confocal microscopy analysis of HeLa cells fed with varying concentrations of **Sia-2TCO** for two days and subsequently labeled with **Tz-Cy3** confirmed the successful incorporation of the analog even at low millimolar concentrations (**Fig.2B**). With U2OS and NK92-MI cells the signal intensity of plasma membrane signal peaked at 2.5mM. Further increase in sugar concentration paradoxically led to a decreased plasma membrane signal suggesting saturation of cellular enzyme or toxicity (**ESI**). We thus concluded that higher concentrations were not required to achieve strong signals and used 1mM of **Sia-2TCO** in subsequent experiments. To assess how incubation time affects the rate of incorporation, we added 1 mM of **Sia-2TCO** to media of U2OS, NK92-MI and THP-1 cells and monitored the signal of incorporated sugar over three consecutive days. A detectable signal was observed on the cell surface after just 24 hours of incubation. The intensity increased further after 48 hours but the rate of increase was less pronounced after 72 hours (**Fig.2C**). Based on this data, we selected 1 mM of **Sia-2TCO** and a 48-hour incubation period as the optimal conditions for subsequent experiments.

**Figure 2.**
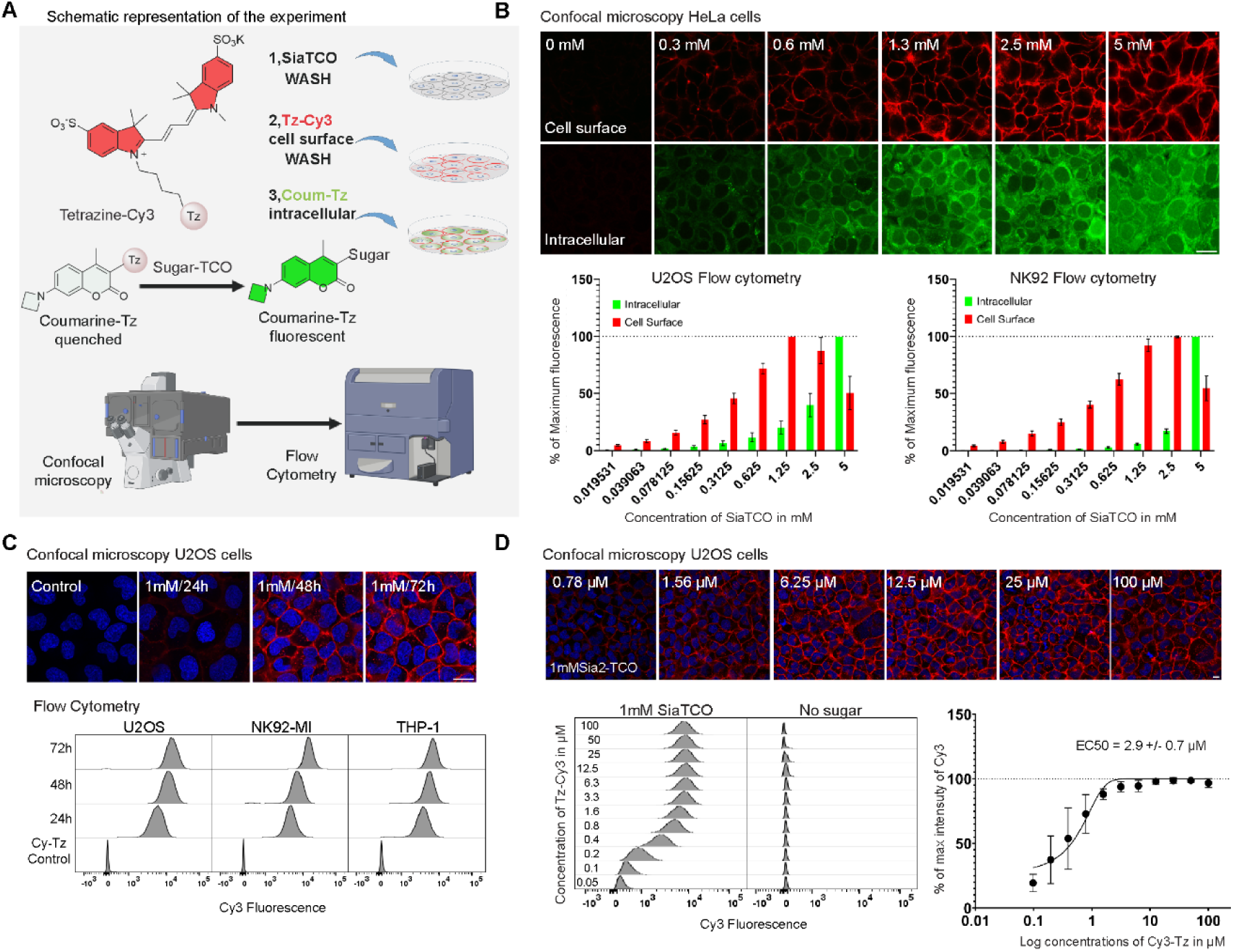
Defining the optimal labeling conditions. **A**) Schematic overview of the double labeling experiment. HeLa, U2OS and NK92-MI cells we incubated with different concentrations of **Sia-2TCO** for 48 hours. After this time, they washed and cell surface exposed sugar was visualized with cell-impermeable **Tz-Cy3** (2.5 µM) followed by intracellular detection of sugars with the cell-permeable **Coum-Tz** (1 µM). Cells were imaged using confocal microscope, detached enzymatically from the dish and further analyzed using flow cytometer. **B**) Confocal pictures of Hela, cells. Red signal shows cell surface sugars, green signal depicsts intracellular sugar. The U2OS and NK92-MI cells were directly analyzed using the flow cytometer. **C**) Optimal time for incorporation. U2OS, NK92-MI and THP-1 cells were incubated with 1mM **Sia-2TCO** for 24-72h, washed and reacted with 2.5 µM **Tz-Cy3**. U2OS cells were photographed each day and detached from dish and fixed using formaldehyde. After the incubation period the fluorescence of Cy3 was measured using flow cytometer. NK92-MI and THP-1 cells were processed similarly with the omission of the microscopy step. Each day picture. Median fluorescence intensities were plotted as a function of sugar concentrations. incubated with different concentrations of sugar ranging from 0 to 5mM. **D**) Finding optimal tetrazine concentration. U2OS were incubated with 1mM **Sia-2TCO** for 48 hours, washed and treated with **Tz-Cy3** (1-100 µM) for 30 minutes. After which time they were using confocal microscope, detached from dish and further analyzed using flow cytometry. The data from flow cytometer are shown as histograms comparing the shift of fluorescence Median fluorescence intensities were then plotted against the concentrations and EC50 for **Tz-Cy3** concentration was estimated with the GraphPad prism software.

### Optimization of proper tetrazine concentration

To determine the optimal concentration of tetrazine, U2OS cells were incubated with **Sia-2TCO** for 48 hours and subsequently reacted with increasing concentrations of Tz-Cy3 (0-100 µM). Cells without **Sia-2TCO** served as controls for background reactivity. Following a 30-minute reaction, cells were photographed using fluorescent microscope and detached from the cultivation dish for fluorescence intensity analysis by flow cytometry. **Fig.2D** shows that the signal intensity increases with increasing concentration of Cy3-Tz but reaches clear plateau. To estimate the effective dose, we plotted the median fluorescence intensities as a function of concentration, and we obtained EC50 of 2.9 µM. In all further experiments we used a slightly lower concentration (2.5 µM).

### Signal specificity

To verify that the cellular staining of TCO is not a result of chemical modification but a result of metabolic incorporation we perturbed several steps of the sialic acid synthesis pathway.

The route N-azido derivatives of mannosamine hijack the N -acetyl mannosamine (ManNAc) salvage pathway that transforms them into sialic acid and ultimately incorporates them into cellular sialoglycans. Cells can also process exogenous sialic acid using the same pathway. When cells are exposed to both sugars, they prefer the fast-accumulating substrate. To test that the **Sia-2TCO** enters this biochemical pathway we co-incubated the U2OS cells with **Sia-2TCO** and peracetylated N-Acetyl mannosamine (Ac_4_ManNAc) that has higher rate of cellular accumulation than the free sialic acid. As shown in **Fig.3A**, addition of 100µM Ac_4_ManNAc blocked the incorporation of **Sia-2TCO** into cellular surface proteins suggesting that the 2 sugars compete for the same enzymes. Importantly co-inbubation with peracetylated 100µM peracetylated N-Acetyl galactosamine (Ac_4_GalNAc) that is utilized by other enzymes of the N-Acetyl galactosamine salvage pathway did not change the incorporation efficiency of **Sia-2TCO** (**Fig.3A**). Cytidine monophosphate N-acetylneuraminic acid synthetase (CMAS) is the key enzyme in the metabolic pathway of sialic acid, responsible for generating CMP-sialic acid in cells. This activated sugar is then ultimately used by sialic acid transferases. To verify that this enzyme is involved in processing **Sia-2TCO**, we prepared a CMAS gene knockout (KO) cell line (**ESI**). These KO cells were unable to incorporate TCO groups into cell surface glycans when grown in the presence of **Sia-2TCO** (**Fig.3B**). The activated sugar CMP-Sialic acid enters the Golgi apparatus and there is used by sialic acid transferases. Finally, we used the known inhibitor of these enzymes 3xFax-Neu5Ac to block the incorporation **of Sia-2TCO** sugar into cellular surface. As shown in (**Fig.3C**) co-incubation of cells with 1mM **Sia-2TCO** and 200 µM **3xFax-Neu5Ac led** to decreased incorporation of TCO into cellular surface.

**Figure 3.**
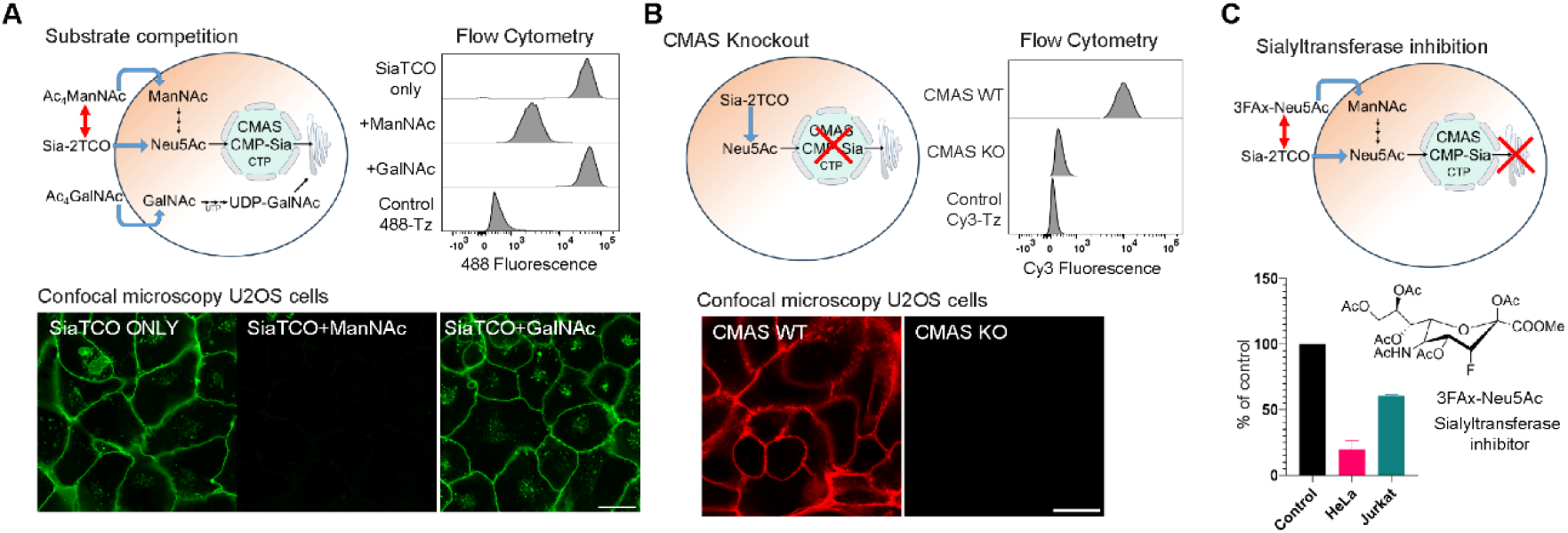
Proof of metabolic **Sia-2TCO** incorporation **A)** Cells use exogenous **Sia-2TCO** similarly to Neu5Ac. The enzyme N-acyl neuraminate cytidylyltransferase (CMAS) activates sialic acids with Cytidyl triphosphate (CTP). This activated sugar is used by sialic acid transferases in Golgi apparatus. Extracellularly added sugars compete for the enzymes in the sialic acid salvage pathway. U2OS were incubated for 48h with **Sia-2TCO** (1mM) and with the combination of **Sia-2TCO**(1mM)/ **Ac_4_ManNAc (**100 µM). **Ac_4_ManNAc** is more hydrophobic due to peracetylation, thus accumulates faster in cells. Higher cellular concentrations then outcompete **Sia-2TCO** and lower the amount of utilized **Sia-2TCO** as documented with low cellular fluorescence TCO visualized by reaction with **Tz-AF488** (2.5 µM). **B)** Upon CRISPs mediated deletion of CMAS the signal of incorporated **Sia-2TCO** sugar (1mM, 48h) is diminished as visualized by click with **Tz-Cy3** (2.5 µM) using confocal microscopy and flow cytometry. **C)** Cells were co-incubated with **Sia-2TCO** (1mM, 48h, control) or combination of Sia-2TCO and peracetylated sialyl transferase inhibitor **3Fax-Neu5Ac** (200 µM). In the presence of the inhibitor the cells showed decreased incorpotation rate of the TCO into cell surface sialoglycans suggesting that cellular sialic acid transferases use the activated **Sia-2TCO**. Scale bars in all microscopic images represent 10µm.

Collectively, these results provide strong evidence that **Sia-2TCO** enters the cellular metabolic pathway of sialic acid and leads to the installation of TCO groups in the glycoconjugates of living cells.

### Comparison of incorporation efficiency of Sia-2TCO with Ac_4_ManN-4TCO and Ac_4_ManN-2TCO

Recent work has shown that TCO-modified mannosamine can be metabolically processed by EMT6^27^ and A549 cells.^28^ Therefore we wanted to compare incorporation efficiency N-substituted mannosamines with our sialic acid derivative. The peracetylated mannosamine analogs were prepared from the known **Ac_4_ManN-Naph** in two steps using the corresponding TCO active esters **(Fig.4A).** We incubated these sugars along with 1mM **Sia-2TCO** with U2OS cells. As initial conditions we chose typical concentrations and incubation time used in MGE experiments that are 48 hours and 50 µM. After the incubation time, the surface TCO was reacted with Tz-Cy3 followed by visualization of intracellular sugars with 1uM Coum-Tz (**Fig.4B)**. This analysis revealed a much stronger intracellular signal from the coumarin probe in cells treated with peracetylated **Ac_4_ManN-4TCO** and **Ac_4_ManN-2TCO** compared to negatively charged **Sia-2TCO**, suggesting that a larger amount of mannosamines penetrated the cells. This finding aligns with the increased lipophilicity of these compounds compared to the sialic acid derivative **Sia-2TCO**. Conversely, minimal signal was detected on the cell surface following treatment with **Ac_4_ManN-4TCO** and **Ac_4_ManN-2TCO**, indicating that, despite their intracellular presence, these compounds were not efficiently processed by cellular biosynthetic machinery and incorporated into cell surface glycans (**Fig.4B**). We repeated the experiment using C33, and Hela cells **(ESI**)

**Figure 4.**
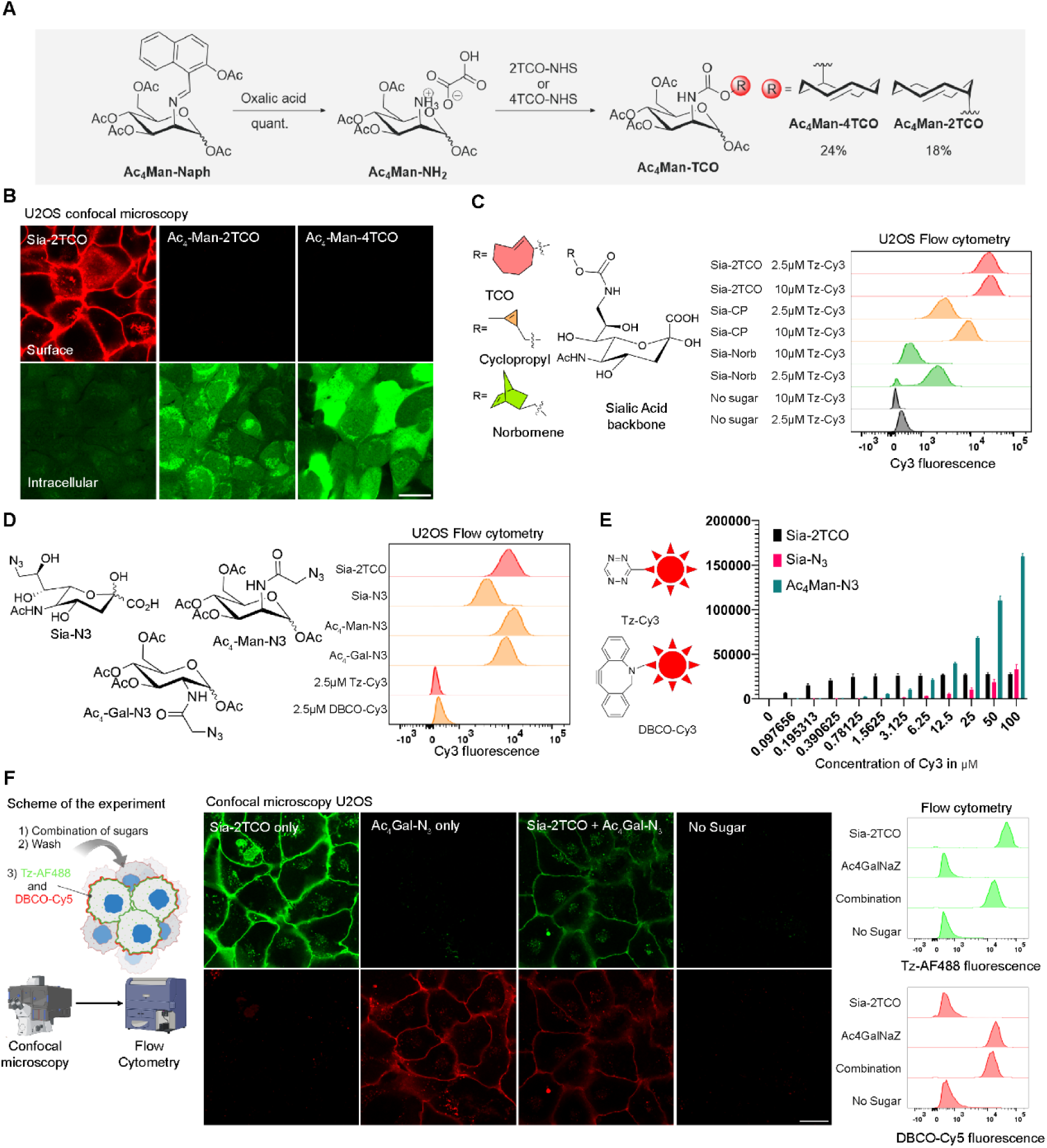
**A)** Synthesis of ManN-TCO sugars. **B)** Confocal microscopy analysis of cells grown in the presence of different amounts **Sia-2TCO** (1mM)**, Ac_4_ManN-2TCO** (100µM) and **Ac_4_ManN-4TCO** (100uM). After 48h of incubation cells were reacted with **Tz-Cy3** (2.5µM), and **Coum-Tz** (1µM) and imaged using confocal microscope. Red signal represents cell surface incorporated glycans, green is intracellular sugar. **C)** Comparison of Sia-2TCO with derivatives containing the same sialic acid scaffold but differing in their reactive groups that operate via the same IEDDA reaction mechanism. U2OS cells were incubated with 1mM concentrations of the indicated sugar washed and reacted with either 2.5 or 10µM **Tz-Cy3** for 30 minutes. After this time cells were detached from the dish and analyzed using flow cytometer. **D)** Benchmarking **Sia-2TCO** against a panel of azide containing sugars utilizing the SPAAC reaction mechanism. Cells were incubated with 1mM of **Sia-2TCO** and **Sia-N_3_** and with 100µM of **Ac_4_Gal-N_3_**, **Ac_4_Man-N_3_** for two days and stained with 2.5 µM **Tz-Cy3** (for Sia-2TCO) or with 2.5 µM **DBCO-Cy3** (Sia-N_3_, Ac_4_Gal-N_3_, Ac4ManN-N_3_) for 30 min. **E)** Graph showing median fluorescence intensity of cells metabolically labeled for 48h with constant concentration of sialic acids (**Sia-2TCO**, **Sia-N_3_**;1mM) and **Ac_4_ManN-N_3_** (50µM). Upon 48h of incubation and wash cells were reacted with range of concentrations of **Tz-Cy3** (1-100 µM) (Sia-2TCO) or of **DBCO-Cy3** (1-100 µM) (Sia-N_3_, Ac_4_Man-N_3_) for 30 min. **F)** Demonstration of orthogonality of IEDDA and SPAAC reactions. U2OS cells were treated with 1mM **Sia-2TCO**, 100 µM **Ac_4_Gal-N_3_** or a combination of these sugars. **G)** Incorporated sugars were visualized using a mixture of 2.5 µM **Tz**-**F488** and 2.5 µM **DBCO-Cy5** and pictured using confocal microscope. **H)** After detachment from the dish they were further analyzed by flow cytometry.

### Comparison Tz/TCO pairing to cyclopropene and norbornene IEDDA labeling

We use in visualization of the TCO groups the inverse electron demand Diels-Alder reaction (IEDDA). Metabolic precursors containing strained alkenes, such as cyclopropene (Cp)^13^ and norbornene (Norb)^14^ have previously been successfully employed together with tetrazine IEDDA in MGE. The second-order rate constants for the reactions of cyclopropenes and norbornenes with tetrazines range from 10⁻¹ to 10 M⁻¹ s⁻¹.^29^ To benchmark **Sia-2TCO** against these dienophile-bearing sugars, we synthesized two sialic acid derivatives: **Sia-Cp** and **Sia-Norb** (ESI). U2OS cells were incubated with 1 mM of each sugar and subsequently labeled with 2.5 or 10 µM **Tz-Cy3** for 30 minutes, followed by flow cytometry analysis (**Fig.4C**). This analysis revealed that cells containing TCO groups exhibited the highest fluorescence intensity. Interestingly, even at 10 µM, the Cy3 signal in cells with Cp and Norb did not reach the intensity of the TCO sugars. These experiments further confirm the superior efficiency of the TCO/Tz pairing for labeling of cell surfaces.

### SPAAC vs Tetrazine Ligation in MGE

The strain-promoted azide-alkyne cycloaddition (SPAAC) is arguably the most employed method for cell surface labeling in MGE. To compare the efficiency of this reaction with tetrazine ligation, we conducted a series of experiments using various azide-containing metabolic precursors. In the first set of experiments, we utilized **Sia-2TCO**, **Sia-N_3_**, and the peracetylated sugars **Ac_4_Man-N_3_** and **Ac_4_Gal-N_3_** as precursors. The sialic acid derivatives were applied at a concentration of 1mM, while the peracetylated sugars were used at 100 µM. Following a 48-hour incubation, we labeled the azide-containing glycoconjugates for 30 minutes with 2.5 µM **DBCO-Cy3**. For cells treated with **Sia-2TCO**, 2.5 µM **Tz-Cy3** was employed (**Fig.4D**). These experiments were performed across three different cell lines **(ESI)**. In all cases, cells were incubated with **Sia-2TCO** exhibited comparable fluorescent labeling with those treated with azido sugars. The highest, signal intensity was observed in cells labeled with **Ac_4_Man-N_3_**, the gold standard in MGE.^30^ In previous experiment with **Sia-Cp** and **Sia-Norb** with we observed increased signal intensity with higher amount of **Tz-Cy3.** In subsequent experiments, we evaluated the efficiency of the click reaction at varying concentrations of the labeling reagent **(Fig.4E**). First we incubated NK92-MI cells with 1mM **Sia-2TCO** and **Sia-N_3_** for 48h. Next, we reacted the TCO or azido groups on the cells with concentrations of **Tz-Cy3** and **DBCO-Cy3** ranging from 0-100 µM and measured the amount of fluorescent signal using flow cytometer. The Cy3 signal intensity on **Sia-2TCO** cells plateaued after 30 minutes using low micromolar concentrations of **Tz-Cy3** in accordance with the previous result (**Fig.4E,2D**). However, the signal from **DBCO-Cy3** on the **Sia-N_3_** cells did not peak even at the highest concentration used **Fig.4E**. To highlight this effect more we also used the standard MGE sugar **Ac_4_Man-N_3_.** In this case even with 100 µM **DBCO-Cy3** the fluorescent signal did not plateau. This indicates that although more azide groups are present on the cell surface of **Ac_4_Man-N_3_** treated cells, SPAAC labeling is significantly less efficient, requiring a much larger quantity of DBCO labeling reagent to attain high staining intensity within a reasonable timeframe. This finding is consistent with the considerably slower kinetics of the DBCO-azide reaction, which is typically 2-3 orders of magnitude slower (*k*_2_ of about 1 M^-1^. s^-1^) than tetrazine ligation.^31^

### Combination of SPAAC and Tetrazine Ligation

As next we aimed to exploit the mutual orthogonality of the TCO/Tz and SPAAC reactions to perform tandem dual-labeling experiments on cells modified with two different chemical reporters. A recent report highlighted an unwanted side reaction between symmetric TCO and azido groups on an antibody, indicating that these two groups may not remain orthogonal under certain conditions.^32^ Additionally, the reaction of TCO with electron-poor azides has been utilized in dipolar cycloaddition-triggered release reactions for prodrug activation.^33^

Given these documented instances, we investigated whether the new **Sia-2TCO** would remain compatible and orthogonal to the azide-alkyne cycloaddition. To explore this, cells were grown in the presence of two metabolic precursors: **Sia-2TCO** and **Ac_4_Gal-N_3_**. We selected this GalNAc derivative because it does not interfere with **Sia-2TCO** incorporation. After 48 hours the incorporated TCO groups were labeled using a green, fluorescent **Tz-AF488** probe, while the azide groups were labeled with a **DBCO-Cy5** conjugate (Fig.4F).

As shown in **Figures 4G** and **4H**, cells cultured exclusively with Sia-2TCO were labeled only with the green **Tz-AF488**, while cells incubated with **Ac_4_Gal-N_3_** alone exhibited signal from **DBCO-Cy5**. No cross-labeling was observed with either **Tz-AF488** or the **DBCO-Cy5**. In contrast, cells cultured with a combination of **Sia-2TCO** and **Ac_4_Gal-N_3_**, showed successful labeling in both channels (Fig.4G, 4H). These experiments demonstrate that the two bioorthogonal reporter groups, TCO and azide, can be efficiently incorporated into cellular glycoconjugates on the same cell, making them available for use in mutually orthogonal metal-free click reactions. This finding opens the door to advanced labeling studies utilizing two popular bioorthogonal ligations within a single biological system.

### Cell surface labeling with biomolecules

Labeling cellular surfaces after MGE typically involves small molecule imaging agents, such as fluorophores or biotin probes, which are used to visualize or pull down the respective glycoconjugates for downstream analysis. In contrast, cell surface engineering through MGE using larger biomolecules remains exceptional and challenging. One reason for this is the low efficiency of the current labeling reactions, which often require a large excess of labeling reagents—an approach that can be costly and impractical.^34^ Intrigued by the exceptional efficiency of tetrazine ligation, even at low reagent concentrations, we explored the potential for decorating cell surfaces with biomolecules of different molecular weights (**Fig.5A**).

**Figure 5.**
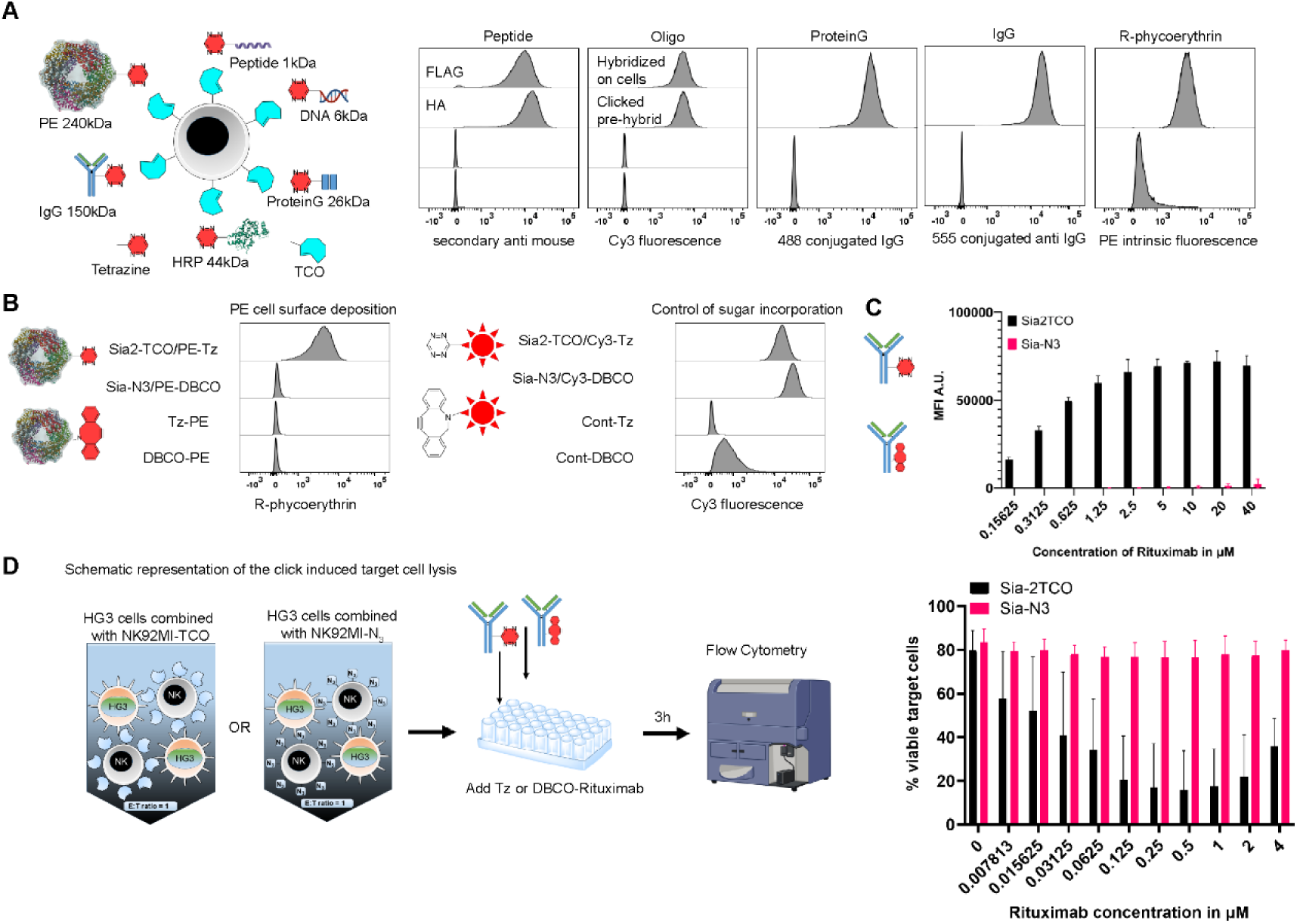
Demonstration of the broad versatility of ChemCell technology. **A)** NK92-MI cells were treated with 1mM of Sia-2TCO for 48h, washed and subsequently reacted with a panel of Tz-decorated biomolecules of varying molecular weights. **PEPTIDE)** NK92-MI-TCO cells were incubated with 2.5 µM of Tz-FLAG or Tz-HA peptide for 30 min, washed and bound molecules were detected using anti FLAG or HA antibodies followed by staining with secondary fluorescent conjugates. Fluorescence was measured using flow cytometry. **OLIGONUCLEOTIDE)** NK92-MI were reacted with 2.5 µM of the hybridized DNA duplex containing two oligos with the tetrazine polyethylene glycol (**Tz-PEG**) moiety and Cy3 dye. In the latter experiment first modified cells with **Tz-PEG** containing single stranded oligonucleotide, followed by incubation with the complementary Cy3 containing ssDNA. As seen in the histogram of fluorescence, the cells could be successfully modified using both approaches. **PROTEIN)** NK92-MI cells were modified with TCO using the standard protocol. Then they reacted with recombinant protein G that was pre-decorated with PEG-Tz linker. The attached protein G was then visualized using mouse IgG-488 conjugate. **IgG)** NK92-MI cells were modified with TCO and then incubated with 4 µM Cetuximab modified with PEG-Tz linker. **PROTEIN COMPLEX)** R-phycoerythrin (PE), was first modified with **Tz-PEG** linkers using active ester chemistry (**Tz-PE**). Then 5 µM of **Tz-PE** was incubated for one hour with U2OS cells that were previously treated with 1mM of TCO for 48h. The attached protein has intrinsic fluorescence that was subsequently measured with flow cytometer upon detachment of cells from the dish. **B)** Benchmarking ChemCell to SPAAC using **Tz-PE** and **DBCO-PE**. U2OS cells were treated with 2mM SiaTCO, 2mM Sia-N_3_ for 48hours, washed and reacted with 5 µM of either the **Tz-PE** or **DBCO-PE.** The reaction time was prolonged to 1 hour after which time, cells were detached from the dish, and the total cell fluorescence was measured by flow cytometry. **C)** Benchmarking ChemCell to SPAAC using **Tz-Rituximab** and **DBCO-Rituximab**. NK92-MI cells were modified with 1mM of both **Sia-2TCO** and **Sia-N_3_** for 48 hours, washed and reacted with different concentrations **Tz-Rituximab** or **DBCO-Rituximab**. The reaction time was prolonged to 1hour. Bound antibody was quantified with secondary anti human antibody and subsequently by flow cytometry. **D)** NK mediated cell lysis. NK92-MI cells were modified with 1mM **Sia-2TCO** and **Sia-N_3_** for 48 hours. Cells were washed and combined in 96 well plate with the target CD20 positive HG3 cells at 1:1 ratio. HG3 cells were labeled with fluorescein to distinguish them from NK92-MI cells. The target cell lysis was initiated by adding Rituximab (Tz or DBCO) in concentrations ranging from 0.008 µM up to 4 µM. After 3 hours of co-incubation cells stained with annexinV and the ratio of annexin positive target HG3 cells were measured using flow cytometry. We plotted the amounts of annexin negative cells as a function of added Rituximab. Marked decrease in viability was observed only when cells were labeled with **Sia-2TCO** and **Tz-Rituximab** was added to the mixture.

### Labeling with peptides

We began our investigation with peptides, which are biomolecules widely utilized as therapeutics, recognition sequences, targeting moieties, and useful tags. To facilitate convenient detection of the peptides following cellular labeling, we employed the well-established FLAG-Tag and HA-Tag sequences. Peptides labeled with a tetrazine (Tz) moiety at the N-terminus (**Tz** -**FLAG** and **Tz -HA**) were incubated with cells treated with **Sia-2TCO** labeled cells under standard conditions (2.5 µM for 30 minutes). After washing, the presence of the peptides was confirmed using primary mouse anti-FLAG or anti HA antibodies, followed by staining with a fluorescently labeled secondary antibody (**Fig.5A, peptide**).

### Labeling with oligonucleotides

Nucleic acids are attractive functional materials for cell surface engineering due to their self-recognition capabilities, outstanding programmability, and diverse structural forms. These characteristics make cell surface-anchored nucleic acids suitable for various technological applications, including promoting cell-cell recognition, regulating cellular pathways, protecting therapeutic cells, and sensing intracellular or extracellular environments.^35^ All these emerging applications would benefit from a technology that allows for the efficient covalent attachment of DNA molecules to cell surfaces. To assess whether our cell modification technology could meet this demand, we prepared a 17-mer DNA strand labeled at the 3’ end with tetrazine via C7 linker (**17-mer-3’-Tz**, sequence: 5’-TTGAATAAGCTGGTAAT-3’-C7-Tz). A complementary strand labeled with a Cy3 dye at the 5’ end was used for detection in the experiments.

To simulate different scenarios, we designed two experiments. In the first, cells grown with **Sia-2TCO** were incubated for 30 minutes with 2.5 µM of hybridized double-stranded DNA, consisting of the Tz-modified oligonucleotide hybridized to the Cy3-labeled primer. In the second experiment, the **17-mer-3’-Tz** was clicked first, followed by hybridization with the Cy3-primer directly on the cell surface. Flow cytometry analysis of these modified cells confirmed successful labeling in both cases (**Fig.5A, Oligo**). Importantly, control experiments using the Cy3-labeled primer alone and with cells that were not cultured with **Sia-2TCO** showed no labeling.

### Labeling with antibodies and protein complexes

Modification of cell surfaces with proteins can be interesting from many aspects. Equipping cell surfaces with enzymes has broad applications, such as proximity labeling for profiling cellular surfactomes, studying cell-cell interactions. ^36^ Additionally, modifying cell surface with enzymes and antibodies have been shown to enhance the efficiency of therapeutic cells.^37^

Particularly antibodies can provide specific recognition elements for diverse antigens, thereby enhancing the targeting capabilities of immune cells. Normally these cells use their cell surface receptors for binding the Fc part of IgG. In our initial testing we used Natural killer (NK) cell line that does not express the CD16 (Fc receptor) unable to bind the IgG. Therefore, we first modified cell surface with tetrazine-decorated recombinant Protein G. This protein enables the attachment of any IgG via their Fc parts leaving the antigen recognizing part facing the cellular environment. After modification with Protein G, we detected it by incubation of the cells with fluorescent 488-IgG conjugate (**Fig.5A, proteinG**). We also investigated the possibility to modify the cell surface of NK92-MI cells with whole monoclonal antibody modified by a Tz-PEG linker. To do this, we incubated NK-92MI cells with the Sia-2TCO for 48 hours then we incubated them for 30 minutes with the modified Tz-Rituximab. The bound antibody was detected with secondary anti-human antibody coupled with Alexa 555 dye (**Fig.5A, IgG**)

We further investigated whether the feasibility cell modification with enzymes using TCO/Tz strategy is and assessed whether the enzyme retains its functionality following attachment. We modified horseradish peroxidase (HRP) with a tetrazine moiety (**HRP-Tz**) using active ester chemistry and attached the enzyme to live U2OS cells at a concentration of 2.5 µM for 30 minutes. Incubating the modified cells with a fluorescent tyramide HRP substrate (**Tyramide AF647**) and H2O2 revealed strong signal on cells modified with peroxidase while the control cells remained devoid of signal **(ESI)**

Finally, we attempted to modify the cells with a protein complex. To do that, we used commercially available R-phycoerythrin (**PE**), a red fluorescent, 240 kDa multimeric protein complex from red algae.^38^ This protein consists of monomers organized into large oligomeric complex reaching 240 kDa. We modified first modified PE with Tz-PEG using NHS chemistry and subsequently reacted it with TCO modified U2OS cells. After 1h reaction of 5µM **Tz-PE** we detached the cells from the dish and measured the fluorescence of attached PE using flow cytometer. The measurement of fluorescence intensity revealed a pronounced shift toward the higher intensities consistent with successful modification (Fig.5A, R-Phycoerythrin).

### Tz/TCO **I**EDDA outperforms SPAAC for protein modification

Next, we directly compared the TCO/Tz IEDDA and Azide/DBCO SPAAC in cellular surface labeling with PE protein complex. U2OS cells were preincubated with 1mM **Sia-2TCO**, **Sia-N_3_** for 48h. Then they were incubated for one hour with 2.5 µM **Tz-PE** and **DBCO-PE** conjugates. Upon 1h of reaction the cells were pictured with confocal microscopy (**ESI)** and further analyzed using flow cytometry. As a control for sugar utilization, we incubated TCO and azide containing cells with 2.5 µM **Tz-Cy3** and **DBCO-Cy3** for the same amount of time. As seen in the histogram (**Fig.5B**) there was large difference in PE deposition between the TCO and azide modified cells. The signal on Azide modified cels was so weak that it was more comparable with control cells that did not receive any sugar. On the other hand, the control cells showed slightly more Cy3 signal on cells treated with **Sia-N_3_** in comparison to **Sia-2TCO.** (Fig.5B). We repeated the experiment with the classical Azide sugars and obtained similar results (**ESI**).

In the previous experiment we used a large protein and a low concentration of Tz/DBCO modified protein. Therefor we repeated the experiment with smaller protein and broader range of concentrations. We also used cells that grow in suspension to avoid the enzymatic treatment that was necessary to detach U2OS cells from the cultivation dish. As next we prepared tetrazine-and DBCO modified humanized anti-CD20 antibody, Rituximab.

NK92-MI cells were metabolically labeled with 1mM **Sia-2TCO** or **Sia-N_3_** for 48 hours after which time they were incubated for 1 hour with varying concentrations of DBCO- or Tz-modified rituximab ranging from 0 to 40µM. The presence of the antibody on the cell surface was confirmed by a fluorescently labeled anti-human secondary antibody and quantified by flow cytometry. Even at the highest concentration tested (40 µM), efficient labeling with DBCO-modified rituximab was not achieved (**Fig.5C**). In contrast, the Tz-modified rituximab reacted efficiently, even at low micromolar concentrations. The estimated EC50 for Tz-Rituximab was 3 µM (**ESI**).

We think that the explanation for this difference could be the high reaction rate of the IEDDA reaction (k₂ ∼ 10^3^ M^-1^ s^-1^). This drives rapid and efficient labeling even at low micromolar concentrations, overcoming the reduced reactivity typically associated with bulky biomolecules. For example, at a 5 µM reagent concentration, the TCO/Tz reaction has a half-life (t_1/2_) of just 3 minutes. In contrast, the SPAAC reaction, with a typical k₂ ∼ 1 M⁻¹ s⁻¹, has a t_1/2_ of around 56 hours under the same conditions. Therefore, the superior kinetics of IEDDA results in much more efficient and cost-effective labeling.

### Antibody dependent cell lysis in the absence of CD16

To further demonstrate the utility of Tz/TCO reaction we assessed its performance under physiologically relevant conditions at low reactant concentrations. The NK92-MI cell line that we used in previous experiments are natural killer cells that are cytotoxic to a wide range of malignant cells. The NK cells lyse their targets usually by mechanism called Antibody-Dependent Cellular Cytotoxicity (ADCC). They initiate the cytotoxic reaction by binding to antibody-opsonized targets via their Fc-receptors (CD16). NK92-MI cells are negative for expression of CD16 and as such cannot use this canonical mechanism to initiate cell lysis. We have decided to restore this capability using our Tz/TCO technology. We modified the NK92-MI with 1mM of **Sia-2TCO** for 48hours. Then we combined them in 1:1 ratio with HG3 characterized by high expression of the CD20 antigen recognized by the Rituximab antibody. Then we initiated the cellular lysis reaction by addition of different amounts of **Tz-Rituximab (Fig.5D)**. As a control we used the same setup with NK92-MI cell line modified with **Sia-N_3_** and the cell lysis was initiated with **DBCO-rituximab**. NK cell activity was evaluated by measuring the fraction of lysed or apoptotic HG3 targets after 3 hours of co-culture. We plotted the amounts of unaffected HG3 cells (intact, annexin negative) as a function of the rituximab concentrations. Graph (**Fig.5D**) clearly shows that in case of TCO/Tz ligation the amounts of viable target cells decrease with increasing concentrations of added antibody. On the other hand, the control experiment with **Sia-N_3_** and **DBCO-rituximab**-initiated target cell lysis shows relatively constant numbers of annexin negative HG3 cells, regardless of the concentration of added antibody. Importantly, the steep decrease of viable target cells occurs at relatively low concentrations lower that of peak levels of therapeutic antibodies observed in blood of patients treated with rituximab (CIT. <u>10.3389/fphar.2022.788824</u>)

## CONCLUSION

This work introduces Sia-2TCO a new sugar for the MGE that uses the Tz mediate IEDDA for efficient modification of cellular surfaces. We name the technology – **ChemCell**. It is a versatile technology for tethering large biomolecules to cell surfaces that outperforms the currently known SPAAC using technologies. The key innovation in ChemCell is the new sialic acid derivative, **Sia-2TCO**, which combines the stability required for metabolic glycoengineering with high reactivity in subsequent click labeling. **Sia-2TCO** integrates into the sialic acid biosynthetic pathway, leading to the installation of *trans*-cyclooctene groups onto cellular glycoconjugates.

Compared to other metabolic precursors containing azide, cyclopropene, or norbornene reporters, **Sia-2TCO** allows much more efficient labeling, making the process both cost-effective and environmentally friendly. The high efficiency of the tetrazine ligation reaction allows the decoration of cell surfaces with a wide range of functional biomolecules, including peptides, oligonucleotides, proteins, enzymes, therapeutic antibodies, and large protein complexes. We demonstrated here only a small number of applications that this technology could be used for.

Given its versatility and efficiency, we anticipate that ChemCell will become a valuable method in basic research as well as in industry as an alternative to genetic methods, opening new opportunities in cell engineering for diagnostics, biotechnology, and cell therapies.

## Supporting information

Experimental supplementary infomation (ESI)

## Acknowledgements

This work was supported by the Academy of Sciences of the Czech Republic (RVO: 61388963); by the project New Technologies for Translational Research in Pharmaceutical Sciences /NETPHARM, project ID CZ.02.01.01/00/22_008/0004607, co-funded by the European Union; by the project TN02000109 Personalised Medicine: From Translational Research into Biomedical Applications co-financed with the state support of the Technology Agency of the Czech Republic as part of the National Centers of Competence Program, and by the European Research Council PoC grant under the EU’s Horizon 2020 research and innovation programme (grant agreement no. 101081736). We thank our colleagues Petro Khoroshyy at Microscopy facility and Jana Günterová at Flow Cytometry facility at IOCB Prague for excellent help.

## MEHTODS

### Cell lines

U2OS (ATCC No.HTB-96), HeLa (ATCC No. CCL-2) C33 (ATCC No.HTB-31) were cultivated in DMEM high glucose supplemented with 10% fetal bovine serum (Thermo, A5256701), NK92-MI (ATCC No. CRL-2408) were cultivated in RPMI 1640 supplemented with 10% fetal bovine serum and 10% heat inactivated horse serum (Thermo, 26050088). THP-1 (ATCC no TIB-202) were cultivated in RPMI 1640 10% Fetal bovine serum. HG-3 cells (DSZM No ACC 765) were grown in RPMI 1640 supplemented with 10% heat inactivated FBS. RAJI (DSZM No.ACC 319) were cultivated in RPMI1640 with 10% h.i. FBS. Jurkat, Clone E6-1 (ATCC No. TIB-152) were cultivated int RPMI1640 + 10% FBS. All cell lines were kept in thermal incubators at 37 degC, 5% CO2 atmosphere.

### Microscopy

For microscopic observations cells were seeded on a 96 well plate dish with a glass bottom (Cellvis, P96-1.5P). Upon treatment they were washed and DMEM medium was changed to Leibovitz’s L-15 (Thermo, No. 21083027) with 10% FBS and pictured using Zeiss LSM 980 confocal microscope. Pictures were exported in using the Zen software and further processed using Fiji software.

### Flow cytometry

Adherent cells were detached from the cultivation dish using Accutase (Thermo No. 00-4555-56). Transferred in V shaped 96 well plate in PBS/5% BSA and measured using flow cytometer BD LSRFortessa Cell Analyzer. Histograms were generated in FlowJo software.

### Dose dependence Sia-2TCO

Hela and U2OS cells were seeded at 15 000/well, in a glass bottom 96 well plate, next day **Sia-2TCO** was added into media at indicated concentrations and cells were incubated for 48h. Media were removed, cells were washed 3x with complete DMEM media (10 % FBS). Click labeling was done on a plate, adding 100 µL of 2.5µM Tz-Cy3 (BroadPharm, BP-23321) for 30 min at 37 degC, washed 3x with complete DMEM medium and further incubated for 10 min with 100 µL of L-15 with 1µM Coum-Tz and photographed on a confocal microscope. NK92-MI cells were seeded at a density of 100 000 per well of 96WP incubated and processed in the same way as HeLa and U2OS except for the microscopy step. In all cases cells were not fixed but analyzed live using the flow cytometer. Median fluorescence intensities were extracted from the raw cytometric data using FlowJo and graphs were generated in GraphPad Prism software.

### Time dependence

15 000/well U2OS or were seeded in a 96 well plate. One day after plating, cells were incubated with 1mM **Sia-2TCO** for 24, 48 and 72 hours. Each timepoint cells were washed with complete DMEM (10%FBS) and incubated with 2.5 µM Cy3Tz (BroadPharm, BP-23321) in complete medium for 30 minutes. (with addition of Hoechst 33342– 5ug/ml to visualize nuclei). After click reaction cells were washed 3x in complete DMEM and pictured using confocal microscope. The same settings of laser intensity and detector sensitivity were used trough the experiment. Each time cells were detached from dish after microscopy and fixed using 4% formaldehyde. Before measurement cells were washed with PBS/5% BSA and fluorescence was measured using flow cytometer. NK-92MI and THP-1 were not pictured using microscope they were washed 3x in their complete cultivation media, incubated with 2.5 µM Cy3-Tz washed fixed with 4% formaldehyde and analyzed on flow cytometer the same way as U2OS.

### Concentration of tetrazine on U2OS

15 000/well U2OS or were seeded in a 96 well plate. One day after plating, cells were incubated with 1mM **Sia-2TCO** for 48 hours. After this time, cells were washed with complete DMEM (10%FBS) and incubated with serial dilutions of Cy3-Tz ranging from 0-100µM in complete medium for 30 minutes. (with addition of Hoechst 33342– 5ug/ml to visualize nuclei). Control cells did not contain **Sia-2TCO.** After the incubation period cells were washed 3x in L15 medium and imaged using confocal microscope. After imaging cells were detached from cultivation plate using Accutase and transferred into V shaper 96 dish for measurement using flow cytometer. Median fluorescence intensities were extracted from the raw cytometric data using FlowJo and graphs were generated in GraphPad Prism software. EC50 was estimated in Prism using nonlinear fit of log (concentration)/normalized response curve.

### ManAc vs SiaTCO competition experiment

15 000/well U2OS or were seeded in a 96 well plate. One day after plating, cells were incubated with indicated sugars either alone or in combination. After 48 hours cells were washed to remove the unincorporated sugar and reacted with 100ul of complete DMEM containing 2.5 µM AF488-Tz and 2.5 µM DBCO-Cy5. The click reaction took place for 30 minutes in thermal incubator. After this time cells were washed 3x to remove the unreacted dyes and imaged using confocal microscope. After the imaging cells were detached from the dish and analyzed using flow cytometer.

### Sialyl transferase inhihitor (2 biological replicates)

200 000 Jurkat cells or 50 000 HeLa cells per well were seeded onto 6-well plate (TPP, 92406). The complete medium (RPMI1640 + 10 % FBS – Jurkat, DMEM + 10 % FBS – HeLa) was supplemented with 200 µM sialyltransferase inhibitor (Sigma, 566224-10MG). After 3 days of incubation, SiaTCO in final concentration 1 mM was added into the medium and cells were incubated for another 2 days. The control cells were treated the same way except adding the sialyltransferase inhibitor. After incubation, the cells were 3 times washed with fresh complete medium and the 2.5 µM Tz-Cy3 (BroadPharm, BP-23321) was added for 30 minutes. Then the cells were washed, HeLa cells were deattached using the accutase and analysed by flow cytometry.

### Comparison of Ac4Man-2TCO and Ac4Man-4TCO with Sia-2TCO

U2OS, HeLa and C33 cell line were seeded at density 12 500 cells per well onto 96-well plate (Cellvis, P96-1.5P). After 24 hours, the medium was exchanged with complete DMEM (10 % FBS) containing sugar derivatives – 1 mM **Sia-2TCO** (AK469), 50 µM **Ac4Man-2TCO** and 50 µM **Ac4Man-4TCO**. Control cells were not incubated with sugar derivatives. After 48 hours of incubation, the cells were 3 times washed with fresh complete medium. Then 2.5 µM Tz-Cy3 (BroadPharm, BP-23321) was added in fresh complete medium and incubated for 30 minutes. After incubation the cells were 2 times washed and complete medium containing 1 µM Coum-Tz was added to the cells. The cells were imaged on confocal microscope.

### SiaTCO vs SiaCp vs SiaNorb (2 biological replicates)

12 500 U2OS cells per well were seeded onto 96-well plate (Celvis). Sugar derivates Sia-2TCO, Sia-CP and Sia-Norb were added in final concentration 1 mM concentration to the cells for 48 hours. The control cells were not incubated with sugar derivatives. After incubation the cells were 3 times washed with complete DMEM (10 % FBS), then 2.5 µM or 10 µM Tz-Cy3 (BroadPharm, BP-23321) was added and after 30 min, the cells were washed with PBS + 1mg/ml BSA and analysed by flow cytometry.

### Comparison of Tz/TCO IEDDA and DBCO/N3 SPAAC

Hela and U2OS cells were seeded at 15 000/well, in a glass bottom 96 well plate, next day 1mM **Sia-2TCO** and **Sia-N3** and 100µM of Ac4Gal-N3, Ac4Man-N3was added. Cell were incubagted with the compounds for another 2 daysafter which time they were washed 3x in complete cultivation media and stained with 2.5 µM Tz-Cy3 (for **Sia-2TCO**) or with 2.5 µM DBCO-Cy3 (Sia-N3, Ac4Gal-N3, Ac4ManN-N3) for 30 min. Cells were detached from dish and analyzed using flow cytometry. Histograms representing median fluorescence intensities were constructed in FlowJo software.

### Comparison of different concentration of Tz-Cy3 and DBCO-Cy3 on NK92-MI cells (3 biological replicates)

100 000 NK92MI per well on a 96 WP were cultured in presence of 1mM SiaTCO or 1mM SiaN3 or 50 µM Ac4Man-N3 for 48 hours. Cells were washed 3x in complete cultivation medium and reacted with different concentrations of TzCy3 or DBCO-Cy3 for 1 hour incubation. After the reaction cells were washed 2x and analyzed using flow cytometry. Median fluorescence intensities were extracted from the raw cytometric data using FlowJo and graphs were generated in GraphPad Prism software.

### Double labeling Sia-2TCO+Ac4Gal-N3

U2OS cells were seeded at a density 15 000/well, in a glass bottom 96 well plate, next day **Sia-2TCO and Ac4Gal-N3** were added into media were added at concentrations 1mM of SiaTCO and 0.1mM **Ac4Gal-N3** and cells were incubated for 48h. Media were removed, cells were washed 3x with complete DMEM media (10 % FBS). Click labeling was done on a plate, adding 100 µL 2.5µM Tz-AF488 and 2.5µM DBCO-Cy3 (with 2000x diluted Hoechst 33342– 5ug/ml) for 30 min at 37 degC, washed 3x with complete DMEM medium and photographed on a confocal microscope. After imaging cells were detached from cultivation plate using Accutase and transferred into V shaper 96 dish for measurement using flow cytometer. Median fluorescence intensities were extracted from the raw cytometric data using FlowJo and graphs were generated in GraphPad Prism software.

### Labelling with FLAG and HA (2 technical replicates)

100 000 NK92-MI cells were seeded per well, SiaTCO in final concentration of 1 mM was added to the cells and incubated for 48 hours. After the incubation, cells were 3 times washed with fresh complete RPMI1640 (10 % FBS + 10% HI Horse serum) and FLAG-Tag-Tz or HA-Tag-Tz was added in concentration 2.5 µM for 30 minutes. Then the cells were 3 times washed with complete RPMI1640 (10 % FBS + 10% HI Horse serum). The FLAG-Tag-Tz and HA-Tag-Tz modified cells were incubated with 1 ug/ml of Monoclonal ANTI-FLAG® M2 antibody (F1804, Sigma) or HA Tag Monoclonal Antibody (26183, Invitrogen) for 30 min. The cells were washed with complete RPMI1640 (10 % FBS + 10% HI Horse serum) and the primary antibodies were detected with Goat anti-Mouse IgG (H+L) Cross-Adsorbed Secondary Antibody, Alexa Fluor™ 555 in dilution 1:400 (Invitrogen, A-21422). The cells were analysed by flow cytometry.

### Labelling with Oligonucleotides

Two complementary synthetic oligonucleotides were purchased form Sigma-Aldrich. One contained NH2 group that was modified using NHS-PEG tetrazine. Olingonucleodide was dissolved in HEPES buffer pH=8.3, 2.5 molar excess of the NHS-PEG tetrazine was added to the oligonucleotide and reaction was proceeded for 1hour at room tempterature. After this time another aliquot of 2.5 molar excels of NHS-PEG was added and the reaction was left on shaker at room temperature overgight. Oligonucleotide was precipitated using Ethanol/Acetate method. The pellet was washed 2x using 70% Ethanol and dried. Then it was dissolved in water. Oligonucletide complementary to the previous one was purchasech with Cy3 fluorophore. The two complementaryu oligonucleotides were hybridized in 100mM NaCl by briefly 1min heating ant then gradually cooling to RT.

1 milion of NK92-MI cells were incubated with the 1mM SiaTCO for 48 hours, washed and 100 000 cells was incubated with the TZ-PEG modified oligonucleotide for 30 min in serum free L15 medium. Cells were washed 2x with the L15 medium (5% BSA) and incubated with 2.5 uM the complementary oligonucleotide in serum free L15 medium with 5% BSA. After this time cells were washed and the fluorescence of attached Cy3 label was measured using flow cytometer.

100 000 NK92-MI cells labeled with TCO was reacted with 2.5 uM of the hybridized dimer containing the Tz-PEG oligo / Cy3 oligo in serum free L15 for 30 minutes. Cells were washed 2x with complete L15 medium with serum and subsequently analyzed in flow cytometer.

### ProteinG labeling

Recombinant protein G was disslved in PBS, then the buffer was exchanged to 100mM NaCl/ 50mM HEPES pH=8.3. Protein was reacted with 2.5 molar excess of NHS-PEG-Tz for one hour, then another portion of 2.5 molar excess of NHS-PEG-Tz was added and the reaction was proceeded for another hour, after chich time unreacted compounds were removed by Zeba desalting column. The Tz-protein G was ready for reaction.

1 milion of NK92-MI cells were incubated with the 1mM SiaTCO for 48 hours, washed and 500 000 cells was incubated with the TZ-PEG modified ProteinG for 30 min serum free L15 medium with 5% BSA. Cells were washed to remove the unreacted protein and incubated with fluorescent Alexa488 conjugate IgG antibody for 15 minutes. After this time cells were washed again and analyzed using flow cytometry.

### Preparation of Phycoerythrin-PEG-Tz conjugates

50 μL of R-phycoerythrin (precipitate 4 mg/mL solution in 60% saturated ammonium sulfate, 50 mM potassium phosphate, pH 7.0, ThermoFischer, P801) was pelleted at 25 000g/10 min. The supernatant was removed and the pellet was resuspended in 100 μL of PBS and buffer-exchanged into 150mM NaCl, 50mM HEPES pH8.3 using Zeba desalting columns (ThermoFischer). Active ester of tetrazine-PEG5-NHS (Conjuprobe, CP-6025) was added to the protein in 5x molar excess (1 µL) from 10 mM stock in DMSO. After one hour of incubation, another aliquot of tetrazine-PEG5-NHS ester (5x molar excess) was added and the reaction was incubated for another 1 hour. After this time unreacted tetrazine was removed using Zeba desalting column giving Phycoerythrin-PEG5-Tz conjugate. The same procedure using DBCO-PEG5-NHS (Broadpharm, BP-24055) ester was used for preparation of Phycoerythrin-PEG5-DBCO.

### Labeling with PE (1 experiment)

12 500 U2OS cells per well were seeded onto 96-well plate (Celvis). SiaTCO or SiaN3 in final concentration 1 mM was added to the cells for 48 hours. The control cells were not incubated with sugar derivative. After incubation the cells were 3 times washed with complete DMEM (10 % FBS), 5 µM and 8 µM Tz-phycoerythrine or DBCO-phycoerythrine was added to the cells for 1 hour. Then the cells were washed and Hoechst (10 mg/ml diluted 1:10 000) in L-15 (10 % FBS) was added and the cells were imaged on confocal microscope and after detachment form dish measured on flow cytometer.

### Comparison of azide and TCO labelling, Rixathon, Modification of antibody

Injection solution (12 mg/ml) – 0.1ml taken, desalted into 50 mM HEPES pH 8.3, 150 mM NaCl on a Zeba spin column. Tetrazine-peg5-NHS ester (Conju Probe, CP-6025) or DBCO-PEG5-NHS (Broadpharm, BP-24055) diluted to 10mM in DMSO, 1 µl taken and combined with the antibody incubated for 1 hour after which time another 1 µl were added to the antibody and incubated for another hour. Then the antibody was desalted on a Zeba spin column into PBS and diluted with glycerol 1:1.

### Gel

Modified antibodies Rixathon-DBCO and Rixathon-Tz (230 ug) were mixed with N3-Cy3 or TCO-Cy3 in final concentration 20 µM and incubated for 1 hour. Then the samples were prepared for electrophoresis either without reduction or with reduction using 100 mM DTT and loaded on 4–15% Mini-PROTEAN® TGX™ Precast Protein Gels, 12-well, 20 µl (BioRad, 4561085). After protein separation the gel was scanned using Amersham™ Typhoon™ Biomolecular Imager. Precision Plus Protein™ Kaleidoscope™ Prestained Protein Standards (BioRad, 1610375)

### DBCO- vs Tz-Rixathon (2 biological replicates)

100 000 NK92-MI cells per well were seeded onto 96-well plate (TPP, 92696). SiaTCO or SiaN3 in final concentration 1 mM was added to the cells for 48 hours. The control cells were not incubated with sugar derivatives. After incubation the cells were 3 times washed with fresh complete medium and modified Rixathon with Tz- or DBCO- was added to the cells in concentration range 0.156 µM to 40 µM and incubated for 1 hour. Then the cells were 3 times washed with fresh medium RPMI1640 (10 % FBS + 10 % HI Horse serum) and Goat anti-Human IgG Fc Cross-Adsorbed Secondary Antibody, DyLight™ 550 (1:100, ThermoFischer, SA5-10135) was added to the cells for 30 minutes. Then the cells were washed and analysed by flow cytometry.

### Modification of antibody in general

Before conjugation of antibody, buffer was exchanged for 50 mM HEPES pH 8.3, 150 mM NaCl and antibody was combined with 5x excess of tetrazine-peg5-active ester (Conju Probe CP-6025) and incubated one hour. After this time another aliquot of tetrazine-PEG5-NHS added and further incubated for another hour. After the incubation buffer was desalted into PBS and antibody was diluted with glycerol (50% final) for storage at – 20 °C.

### NK mediated cellular cytotoxicity experiment

NK92-MI cells were seeded at a density 10 million / 10 cm dish (final volume of the dish is 10 ml of medium sugars (**Sia-2TCO and Sia-N3**) were added to reach 1mM final concentration. Cells were incubate for 48 hours to allow the incorporation into sialoglycans.

After this time cells labeled with **Sia-2TCO and Sia-N3** were collected into 15 ml tubes and spin 300xg/3min, cells were washed 3 times with 10ml of complete medium and spun at 300xg/3min. Ther they were resuspended in 3 ml o medium and counted.

The target HG-3 cells, grown on a 10 cm plate were collected similarly in 15ml tube washed twice in PBS and resuspend in 2 ml of PBS (approx. 10 millions of cells). Thery were labeled with fluorescein (5(6)-Carboxyfluorescein diacetate N-succinimidyl ester, sigma #**21888**, **1µM** final concentration in PBS, for 30 min and washed with complete cultivation medium for NK92-MI cells (RPMI1640 with 10% of FBS and 10% of horse serum) and counted. Cells were distributed into wells of a 96 WP dish at 100 000 per well.

Both - the target HG-3 and sugar labeled NK92-MI were combined in 96 well plate with V shaped bottom at a count 100 000/100 000 i.e 1:1 ratio. The plate was centrifuged 300xg/3min. And media was removed In another plate serial dilutions ranging from 0-4uM of either Tz or DBCO labeled Rituximab antibody was prepared. Each well contained 150 ul of the antibody at a given concertation in RPMI1640 (10% FBS, 10 % Horse serum). 100 ul of each dilution of antibody was combined with the Effector/Target cell mixture and the antibody mediated cell lysis was initiated.

After 3h, the plate was spun at 300g/3min, supernatant was removed and cells were washed twice using 150 ul of Annexin binding buffer (10mM Hepes pH=7.4, 140mM NaCl, 2.5 mM CaCl2). After las wash cells were resuspended in 50µl of Apoptotic/Dead cells labeling solution. And incubated for 15 min at room temperature.

Apoptotic/Dead cells labeling solution: 1/20 dilution of AnnexinV-Pacific blue (#A35122 from Thermo) +1/5000 eFluor780 (Thermo #65-0865-18)

After this time cells were spun 300g/3min, washe in 150µl of Annexin binding buffer and fixed with 4% formaldehyde solution in PBS.

Cells were then measured using flow cytometer. First the cells were gated for target cells (488 - 530/30) then in this gate we selected either the dead cells (640 – 780/60) or the apoptotic cells (405 – 450/40)

### Checking the labelling of NK92cells with antibody

100 000 of labeled cells were taken into a V-shaped 96 WP, spun 300xg/3 min. Resuspended in PBS/BSA with 1/100 dilution of the Goat anti-Human IgG 555-conjugate and incubate for 30 min at room temperature. Washed with 150µl of PBS/BSA once and fix with 4% formaldehyde. Then they were measured using flow cytometer.

## Figures and figure legends

**Scheme 1.**
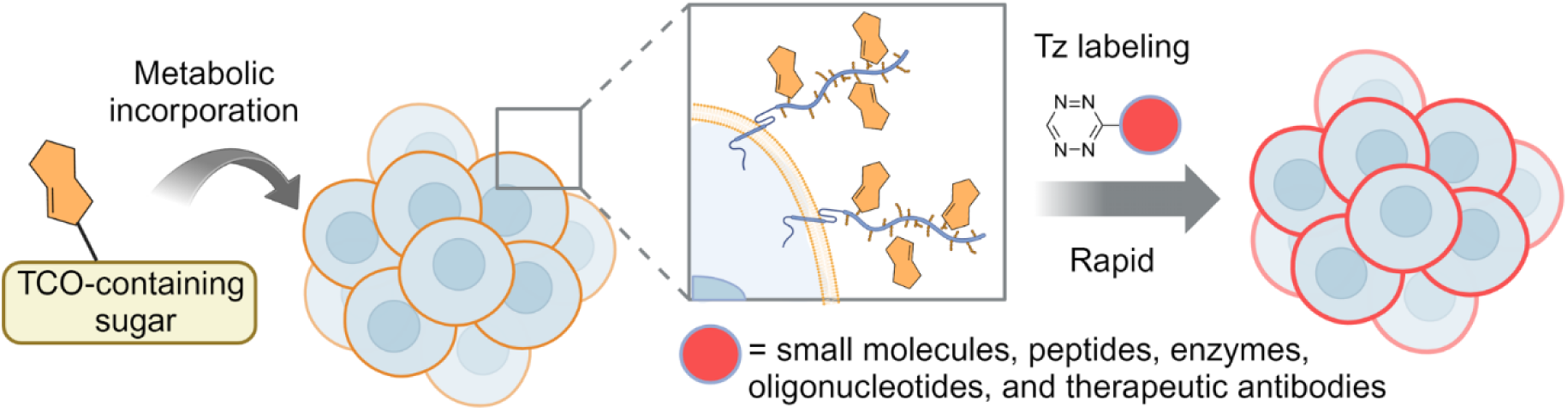
General concept of tethering biomolecules to cell surfaces through MGE with TCO sugars and tetrazine ligation

## Supplementary figures

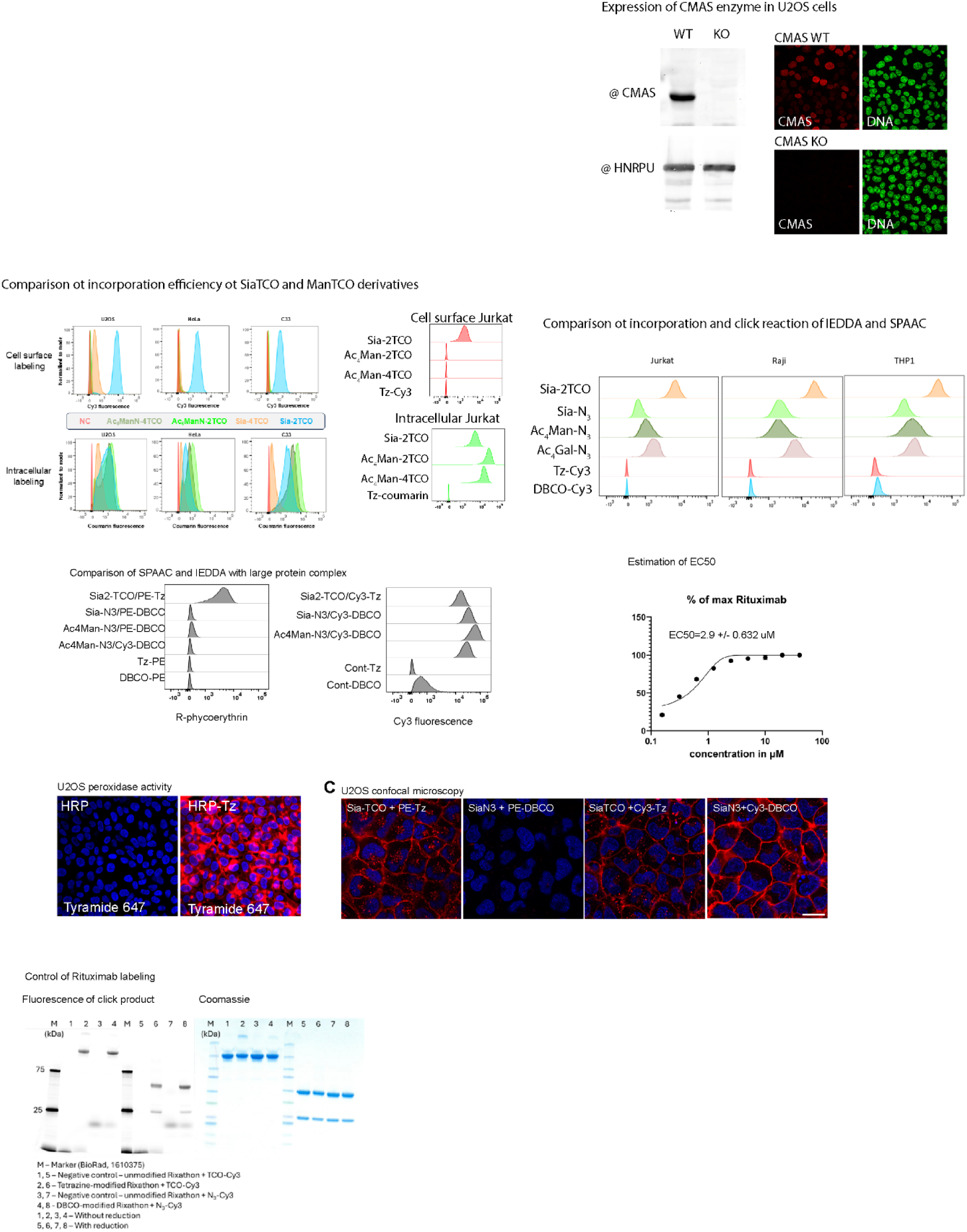

