## Supplementary material for "ChemCell: Chemical Tethering of Large Biomolecules to Cell Surfaces through Diels-Alder Ligation": Experimental supplementary infomation (ESI): ESI.pdf

**Supplementary information**

**Stability of Sia-2TCO and Sia-4TCO**

Sia-2TCO and Sia-4TCO (ca. 7 mg) were separately dissolved in $D_2O$ (0.4 mL) and $^1H$ NMR spectra of the solutions were recorded over time (on a 400 MHz machine). Meanwhile, the samples were stored in the dark at room temperature (NMR cuvette wrapped in aluminum foil, with parafilm around the cap). The NMR cuvettes were shaken before each measurement to homogenize the solution. Residual peak of water was used as an internal standard.

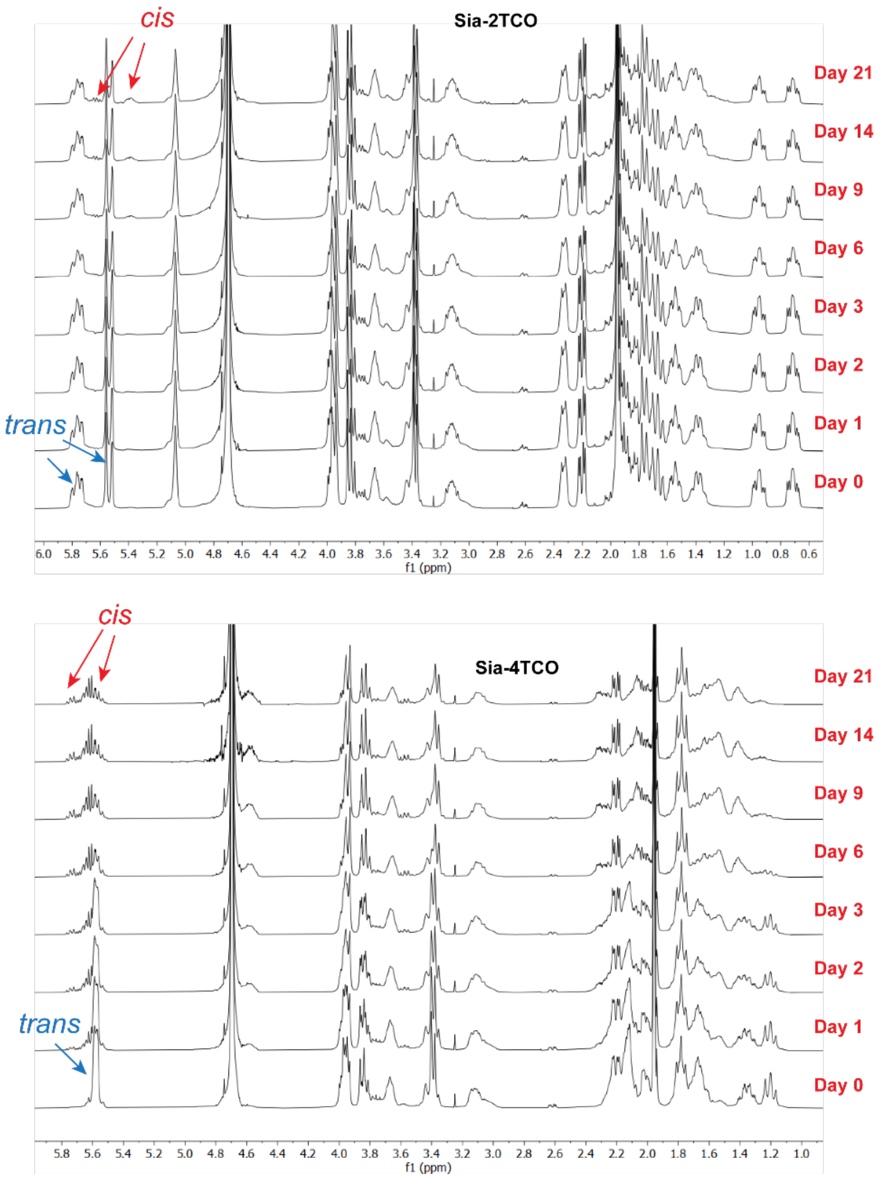


*Figure S1.* $^1H$ NMR spectra of Sia-2TCO and Sia-4TCO measured in $D_2O$ at different time points.

Toxicity on PBMC and HUVEC cells was performed by CellTiter-Glo Cell Viability assay following the instructions of the provider (Promega). These measurements showed that Sia-2TCO has an $IC_{50}$ above 5 mM. The cell viability was evaluated also on U2OS cells as follows: Cells seeded at 15 000/well density were incubated with Sia-2TCO, or sialic acid (control) at concentrations ranging from 5 mM to 0 mM for 48 h. For cell viability assay using crystal violet, the cells were fixed with ice-cold methanol for 10 min at -20°C. Fixed cells were then washed 3× with $H_2O$ and incubated with 0.1% (w/V) solution of crystal violet for 15 min. Cells were washed 3× with $H_2O$ and the bound dye was dissolved using methanol. Absorbance was measured at 595 nm using Thermo Multiscan FC spectrophotometer. The resazurin cell viability assay was performed by following the instructions of the provider (Thermo Scientific).

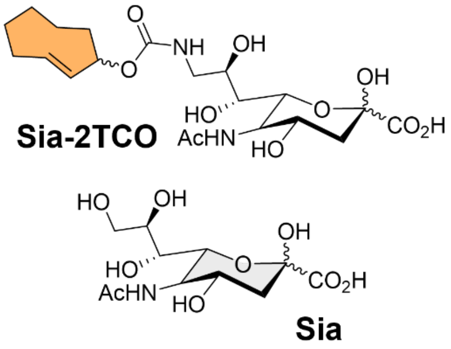


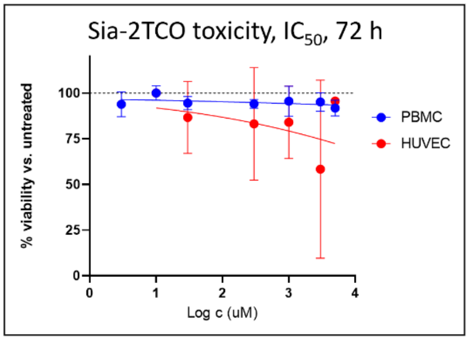


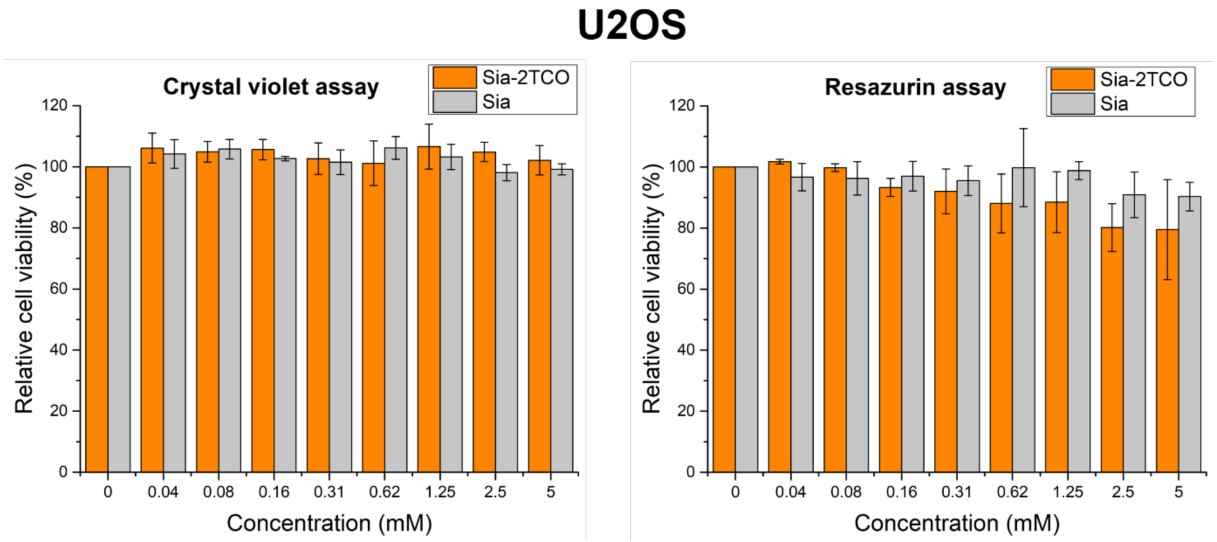

*Figure S2.* Cell viability of PBMC, HUVEC and U2OS cells incubated with the indicated concentrations of sugars. Two different assays were performed, and the results show cell viability relative to control, untreated cells or to cells treated with natural sialic acid (mean ± SD, n = 3, technical replicates).

**CMAS KO**

Guide RNA targeting the sequence CTGCAGCGCAACTCTCGCGG was cloned into lentiCRISPR v2-Blast plasmid (Addgene No #83480). U2OS Cells were transfected with the plasmid, 24 hours later they were selected with the blasticidine (Thermo 10µg/ml No. A1113903) after one-week cells were fed with the SiaTCO and for 48 hours, click labelled with the Cy3Tz and sorted into 96 well plates. Each well contained one cell. After the outgrowth cells were passaged, replicas of each plate were then analyzed for expression of CMAS protein using the anti CMAS polyclonal antibody (Sigma-Aldrich No. HPA039905). CMAS Negative clones were expanded and used in experiments.

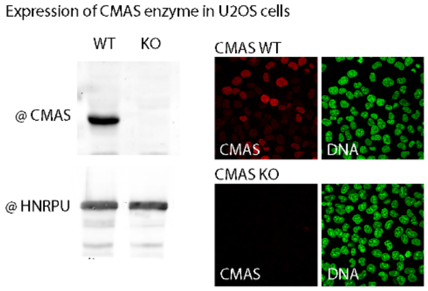


*Figure S2.* Expression of CMAS in U2OS cells visualized by anti CMAS antibody on western blot form U2OS cellular extract. Cells were plated on microscopy glass, fixed with 4% formaldehyde and permeabilized using 0.1% TX-100. CMAS expression was visualized using the same polyclonal antibody as on the western blot. and in cells using confocal microscope (red signal) green signal is DNA (Hoechst 33258).

**Incorporation Study**

U2OS, HeLa and C33 cell line were seeded at density 12 500 cells per well onto 96-well plate (Cellvis, P96-1.5P). After 24 hours, the medium was exchanged with complete DMEM (10 % FBS) containing sugar derivatives – 1 mM Sia-2TCO, 1 mM Sia-4TCO, 50 µM $Ac_4$Man-2TCO, and 50 µM $Ac_4$Man-4TCO. Control cells were not incubated with sugar derivatives. After 48 hours of incubation, the cells were 3 times washed with fresh complete medium. Then 2.5 µM Tz-Cy3 (BroadPharm, BP-23321) was added in fresh complete medium and incubated for 30 minutes. After incubation the cells were 2 times washed and complete medium containing 1 µM Coum-Tz and DRAQ5 (dilution: 1:1000, ThermoFischer, 62251) was added to the cells. The cells were imaged on confocal microscope and analyzed by flow cytometry.

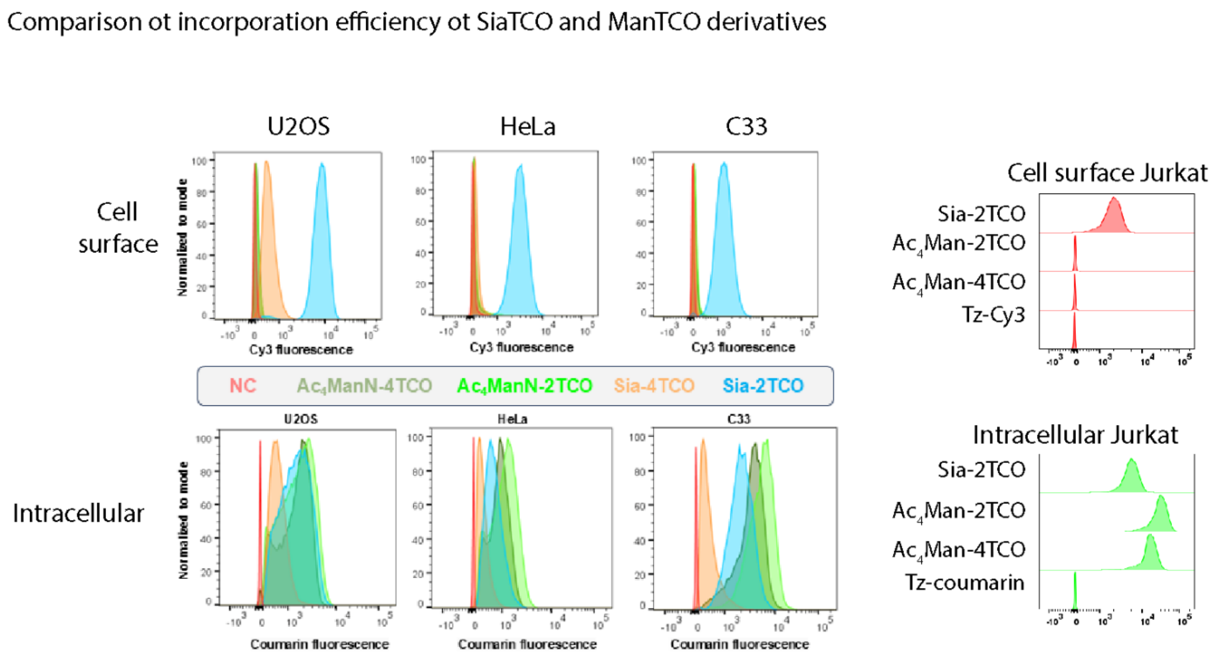


*Figure S3* Flow cytometry analysis of U2OS, HeLa C33 and Jurkat Cells incubated with 1 mM Sia-2TCO, 1 mM Sia-4TCO, 50 µM Ac4Man-2TCO, and 50 µM Ac4Man-4TCO for 48h. After this time cells were washed and reacted with 2.5 µM Tz-Cy3 after brief wash cells were reacted with 1 µM Tz-Coum to highlight the intracellular sugar. Cells were analyzed using flow cytometry.

**SPAAC vs IEDDA on suspension cells**

One million of suspension cells (Raji, Jurkat, THP1) was seeded on a 6 well plate and incubated with 1mM Sia-2TCO, 1mM Sia-N3 , 100µM Ac4Gal-N3, 100µM Ac4Man-N3 for 48h, transferred into falcon tube, spun down, washed 2x in 1ml of L15 medium, after second wash cells resuspended in 1ml of L15 medium and either 2.5uM Tz-Cy3 od 2.5uM DBCO cy3 added and incubated for 30 min on a rotator. After this time cells were washed 2x in 1 ml PBS and 100 µl of this suspension was measured by flow cytometry.

Comparison ot incorporation and click reaction of IEDDA and SPAAC

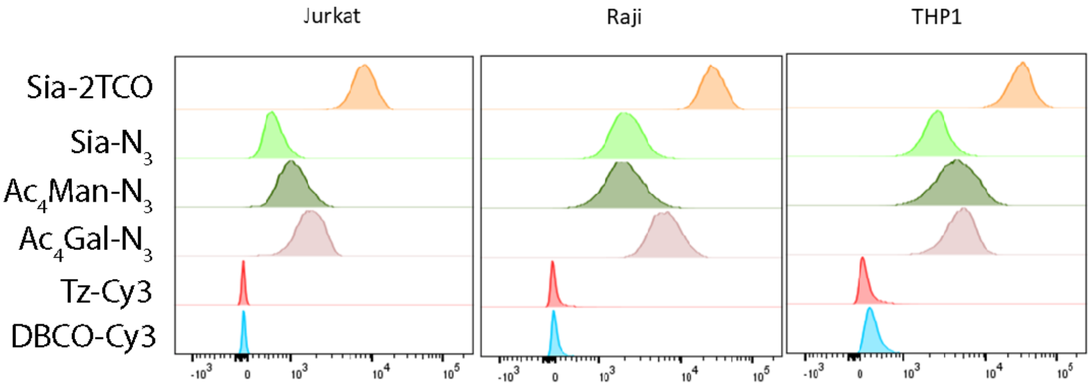


*Figure S4* Comparison of incorporation and of various sugars in suspension cell lines.

**HRP peroxidase**

12 500 U2OS cells per well were seeded onto 96-well plate (Celvis). SiaTCO in final concentration 1 mM was added to the cells for 48 hours. The control cells were not incubated with sugar derivative. After incubation the cells were 3 times washed with PBS, then the cells were fixed with 4 % formaldehyde (P-Lab) for 15 min at RT. After 15 min the cells were washed with PBS and 0.1% Triton-X in PBS was added to the cells for 15 min. Then the cells were 3 times washed with PBS-Tween + BSA. Stock solution of modified HRP with Tz-NHS or unmodified HRP in concentration (12 mg/ml) was diluted 1:500 in PBS + 0.1 %Tween + BSA and added to the cells. After 30 min, the cells were 3 times washed with PBS and 0.015 % H2O2 (31642-500ML-D, Sigma-Aldrich) + 1:100 AF647 Tyramide reagent (from 100x stock in DMSO, ThermoFischer, B40958) were added to the cells and incubated for 15 min in PBS. Then the solution was

exchanged for PBS + 0.1 %Tween with Hoechst (10 mg/ml) diluted 1:10 000. The cells were imaged by confocal microscope.

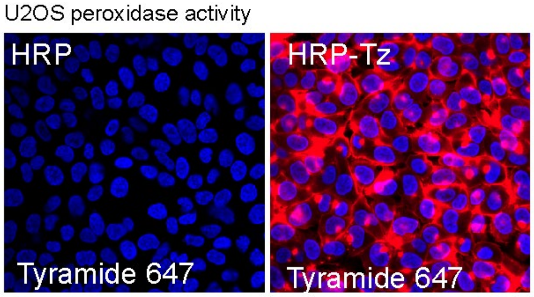


*Figure S5* U2OS cells were modified with Sia-2TCO and reacted with HRP-Tz. Control cells are without sugar. HRP was then visualized using AF647Tyramide reaction.

**Labelling with PE (2 experiment)**

U2OS cells were co-incubated with Sia-2TCO, Sia-N3 (both 1mM) and with 100µM of Ac4Gal-N3, Ac4Man-N3 for 48h. After this time cells were washed to remove unincorporated sugars and subsequently reacted with 2.5uM Tz-PE or DBCO-PE conjugates for one hour. As a control of sugar incorporation. All reactants were in complete DMEM medium  (10% FBS ) and with addition of Hoechst 33342– 5ug/ml to visualize nuclei. After the 1h of reaction time cells were imaged using confocal microscope and detached from dish using accutase and further analyzed using flow cytometry.

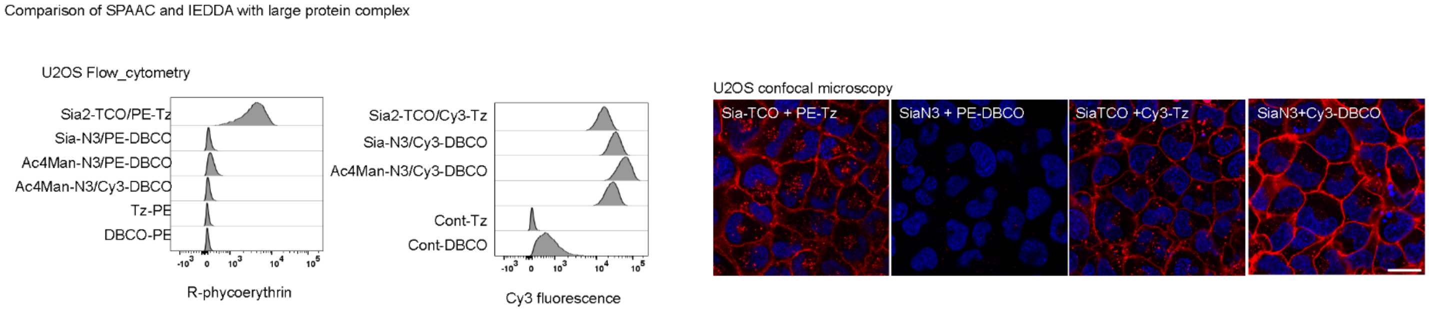


*Figure S6* Comparison of click efficiency using different azide sugars and Sia2TCO. Cells were modified with the indicated sugar and reacted with either Tz-PE or DBCO-PE, imged on confocal microscope and analysed using flow cytometry.

Estimation of EC50

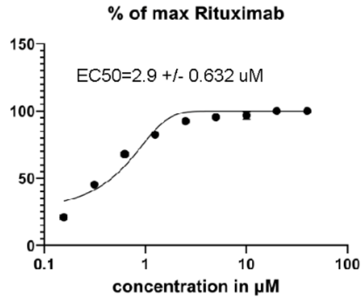


*Figure S7* estimation of the EC50 for Rituximab-Tz. Cells were analyzed using flow cytometer. Median fluorescence intensities were extracted form raw data using FlowJo. Data were plotted in Prism as % of maximum intensity.

**Comparison of azide and TCO labelling, Rixathon, Modification of antibody**

Injection solution (12 mg/ml in infusion solution, Pharmacy, VFN Prague, produced by Sandoz GmbH, Kund) – 0.1ml taken, desalted into 50 mM HEPES pH 8.3, 150 mM NaCl on a Zeba spin column. Tetrazine-peg5-NHS ester (Conju Probe, CP-6025) or DBCO-PEG5-NHS (Broadpharm, BP-24055) diluted to 10mM in DMSO, 1 µl taken and combined with the antibody incubated for 1 hour after which time another 1 µl were added to the antibody and incubated for another hour. Then the antibody was desalted on a Zeba spin column into PBS and diluted with glycerol 1:1.

Gel

Modified antibodies Rixathon-DBCO and Rixathon-Tz (230 ug) were mixed with N3-Cy3 or TCO-Cy3 in final concentration 20 µM and incubated for 1 hour. Then the samples were prepared for electrophoresis either without reduction or with reduction using 100 mM DTT and loaded on 4–15% Mini-PROTEAN® TGX™ Precast Protein Gels, 12-well, 20 µl (BioRad, 4561085). After protein separation the gel was scanned using Amersham™ Typhoon™ Biomolecular Imager. Precision Plus Protein™ Kaleidoscope™ Prestained Protein Standards (BioRad, 1610375)

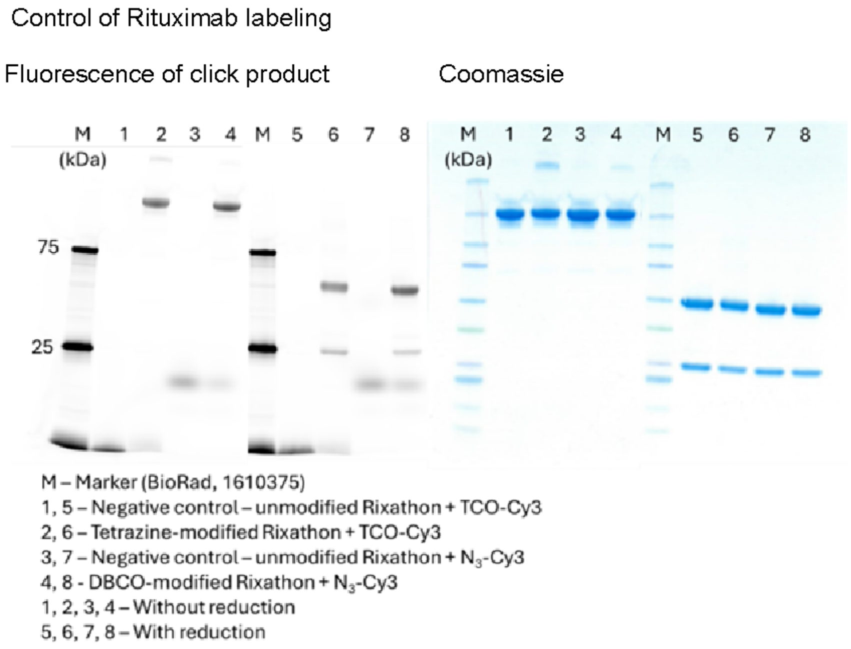


*Figure S8* click labelling and gel analysis of the Tz-PEG and DBCO-PEG Rixathon antibody.

### Materials

##### General methods

The chemicals were obtained from *Sigma Aldrich*, *Acros Organics*, *ABCR, Flouorochem, Iris Biochem, Carbosynth* or *VWR* unless otherwise stated, and were used without further purification. Solutions were concentrated in a rotary evaporator from *Heidolph* equipped with a PC3001 VARIOpro pump from *Vacuubrand.* Column chromatography was carried out on silica gel 60 Å (particle size: 40-60 µm) from *Acros Organics*. p.a.-quality solvents from *Lach-Ner* and *Penta* were used for elution. A Combi*Flash*® Rf+ from *Teledyne ISCO* was used for flash column chromatography. $^{1}$H- and $^{13}$C-NMR spectra were measured in a Bruker Avance III™ HD 400 MHz NMR system equipped with a Prodigy cryo-probe or in a Bruker Avance III™ HD 500 MHz Cryo. Analytical HPLC-MS and fluorescence measurements were performed on an LCMS-2020 system from *Shimadzu* equipped with a RF-20A XS Prominence fluorescence detector, and Luna® C18(2) column (3 µm, 100A, 100 × 4.6 mm) or CORTECS column (C18 from Waters). Preparative HPLC was performed in HPLC-MS Infinity 1260 system equipped with 6120 Quadrupole LC/MS detector from *Agilent Technologies* and either preparative column Luna® 5 µm C18 (2), 100 Å, 250 x 21.2 mm (*Phenomenex*) or preparative column Actus Triart C18, 250 x 20 mm, S-5µm, 12 nm *(YMC)*.

*Oligonucleotides*: DNA oligonucleotide modified at the 3' end with Amine-C7 (sequence 5'-TTGAATAAGCTGGTAAT-3'-[AmC7] and modified at 5' end with Cy-3 dye (sequence [Cy3]-5'-ATACCAGCTTATTCAATT-3'were obtained from Sigma Aldrich.

*Antibodies*: Rituximab – Rixathon (500 mg in infusion solution, Pharmacy, VFN Prague, produced by Sandoz GmbH, Kundl).

### Synthetic procedures

#### Preparation of (E)-5-Acetamido-9-{[(R,E)-(cyclookt-2-en-1-yl)oxycarbonyl]amino}-3,5,9-trideoxy-D-glycero-D-galacto-non-2-ulopyranosic acid (Sia-2TCO)

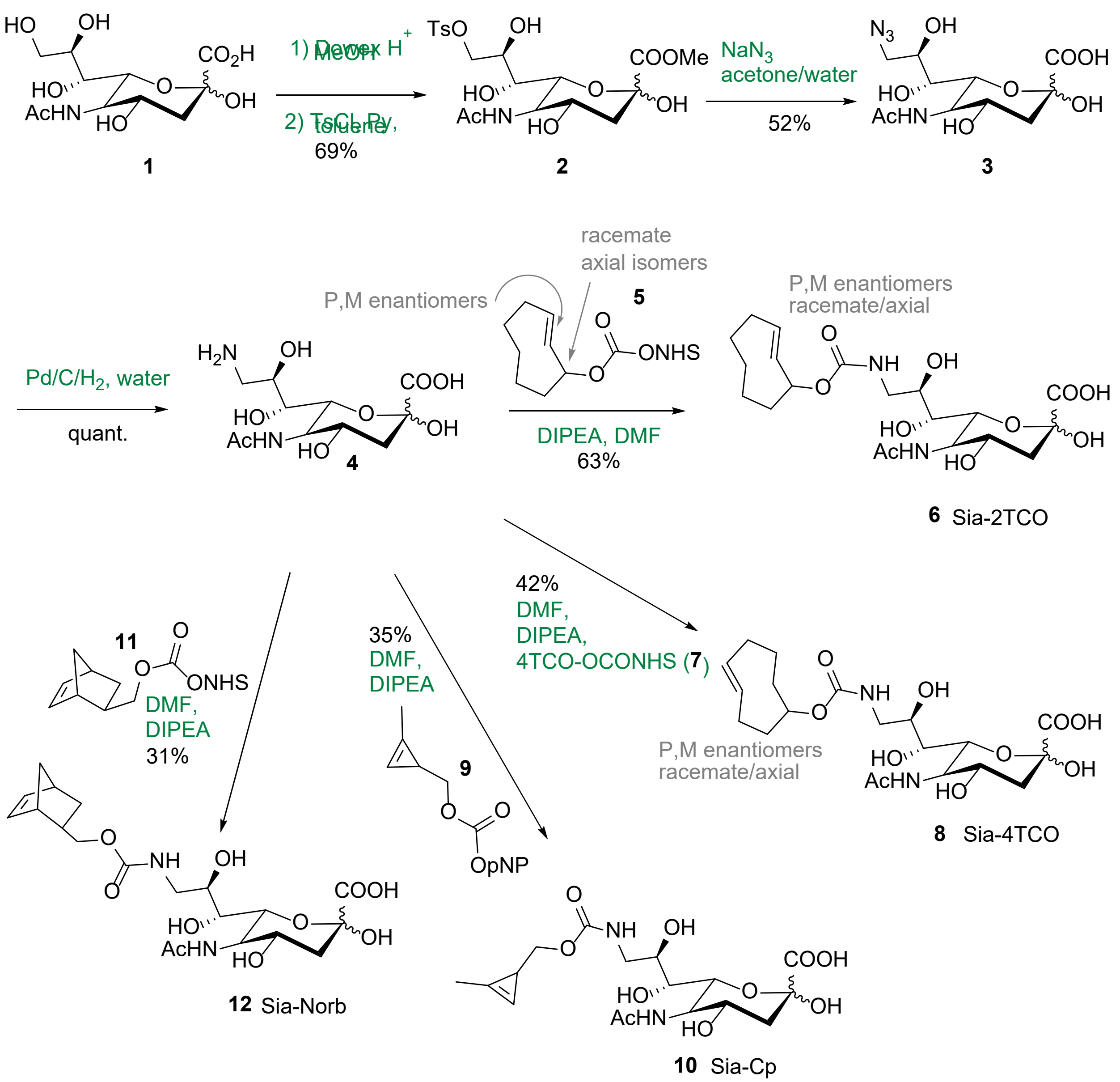

#### Preparation of (E)-5-Acetamido-9-{[(R,E)-(cyclookt-2-en-1-yl)oxycarbonyl]amino}-3,5,9-trideoxy-D-glycero-D-galacto-non-2-ulopyranosic acid (Sia-2TCO)

1. Freshly prepared Dowex50WX8 in $H^+$ cycle (4 g) was added to sialic acid (**1**; 15 g; 48.5 mmol) dissolved in dry MeOH (300 mL), and the mixture was stirred under an argon atmosphere for 48 hours. The resulting clear solution of the product was concentrated under vacuum to yield the methyl ester of sialic acid as an off-white solid. Crystallization (MeOH - EtOAc; 2 : 3) yielded the methyl ester of sialic acid, 14.7 g (94 %). Analytical data agreed with lit.[1]. $^{1}$H NMR (400 MHz, MeOD) δ = 1.90 (dd, *J*=12.9, 11.3, 1H, H-3a), 2.03 (s, 3H, NHAc), 2.23 (dd, *J*=12.9, 4.9, 1H, H-3b), 3.49 (dd, *J*=9.1, 1.5, 1H, H-7), 3.63 (dd, *J*=11.2, 5.7, 1H, H-9a), 3.71 (ddd, *J*=8.8, 5.7, 2.8, 1H, H-8), 3.79 (s, 3H, $OCH_3$), 3.77 – 3.88 (m, 2H, 10, H-5), 3.97 – 4.10 (m, 2H, H-6, H-4). $^{13}$C NMR (101 MHz, MeOD) 22.64 (NHCO*C*$H_3$), 40.70 (C-3), 53.14 (C-5), 54.33 ($OCH_3$), 64.83 (C-9), 67.85 (C-4), 70.19 (C-6), 71.64 (C-8), 72.09 (C-7), 96.68 (C-2), 171.77 (*C*OO$CH_3$), 175.12 (NH*C*O$CH_3$).

The methyl ester of sialic acid **2** (4 g; 12.4 mmol) was dissolved in dry pyridine (80 mL), and the mixture was cooled down on ice before the solution of paratoluenesulfonyl chloride (3.1 g; 16.1 mmol) in dry toluene (40 ml) was added. After stirring for 18 hours at room temperature, the solvent was removed under vacuum, co-distilled with toluene and the resulting crude product was purified by silica gel flash column chromatography (using a gradient of MeOH in DCM 0→40%) to obtain the product **2** as a white solid (4.1g; 69 %), which was used without further purification in the next step. Analytical data agreed with lit.[2] $^{1}$H NMR (401 MHz, MeOD): 1.88 (dd, *J* = 13.0, 11.4 Hz, 1H, H-8a), 2.02 (s, 3H, C**H**$_3$CO), 1.97 – 2.07 (m, 1H), 2.21 (dd, *J* = 12.9, 4.9 Hz, 1H, H-8b), 2.47 (s, 3H, $CH_3$), 3.34 – 3.48 (m, 1H, H-7), 3.74 (d, *J* = 10.3 Hz, 1H, H-5), 3.78 (s, 3H, $CH_3O$), 3.87 (ddd, *J* = 9.2, 6.0, 2.3 Hz, 1H, H-8), 3.95 (dd, *J* = 10.5, 1.5 Hz, 1H, H-6), 3.98 – 4.11 (m, 2H, H-4, H-9a), 4.24 – 4.34 (m, 1H, H-9b), 7.41 – 7.49 (m, 2H, H-3', H-5'), 7.76 – 7.86 (m, 2H, H-2', H-6'). $^{13}$C NMR (101 MHz, MeOD): 21.57 ($CH_3$), 22.63 (**C**$H_3$CO), 40.72 (C-3), 53.17 ($CH_3O$), 54.28 (C-5), 67.68 (C-4), 69.27 (C-8), 69.96 (C-7), 71.83 (C-6), 73.76 (C-9), 96.62 (C-

2), 129.10 (C-2', C-5'), 131.03 (C-3', C-5'), 134.25 (C-1'), 146.41 (C-4'), 171.65 ($CH_3O$**C**O), 175.19 ($CH_3$**C**O).

2. Sodium azide (0.83 g; 12.8 mmol) was added to compound **2** (1.5 g; 3.2 mmol) dissolved in acetone/water mixture (3/1, 24 mL), and the resulting mixture was stirred at 73°C for 5-18 hours. After cooling down to room temperature, the solvents were removed under vacuum and the residue was co-evaporated with EtOH and further dried. The crude reaction product was purified by C18 flash column chromatography (using water as the eluent). The product (**3**) was isolated as a beige syrupy residue and was repurified by preparative HPLC (C18 Triart column, using water). Product **3** was isolated (0.57 g; 53 %) as a white solid after lyophilization. Analytical data in accordance with lit.[2] $^1$H NMR (400 MHz, MeOD): 1.90 (dd, *J* = 12.7, 11.3 Hz, 1H, H-3a), 2.03 (s, 3H, C**H**$_3$CO), 2.09 – 2.18 (m, 1H, H-3b), 3.33 – 3.44 (m, 2H, H-7, H-9a), 3.45 – 3.58 (m, 1H, H-9b), 3.85 (ddd, *J* = 9.3, 6.8, 2.7 Hz, 1H, H-8), 3.88 – 3.96 (m, 1H, H-5), 3.97 – 4.09 (m, 2H, H-5, H-6). $^{13}$C NMR (101 MHz, MeOD): 22.81(**C**H$_3$CO), 41.73 (C-3), 54.05 (C-5), 55.73 (C-9), 68.78 (C-4), 71.10 (C-8), 71.16 (C-7), 71.76 (C-6), 97.57 (C-2), 174.43 ($CH_3$**C**O), 177.16 (COOH).

3. Pd/C (10%, 150 mg) was added to compound **3** (0.87 g; 2.6 mmol) dissolved in water (35 mL), and the flask was purged with $H_2$ gas. The mixture was stirred under an $H_2$ atmosphere at room temperature for 4.5 hours. The crude reaction mixture was passed through a pad of celite to remove Pd/C, and the resulting aqueous solution was lyophilized to yield product **4** as a beige powder (0.3 mg; 98 %), which was used without further purification in the next step. Data in accordance with lit[2]. $^1$H NMR (400 MHz, $D_2O$) 1.78 – 1.88 (m, 1H, H-3a), 1.97 – 2.09 (m, 1H), 2.05 (s, 3H, C**H**$_3$CO), 2.17 – 2.26 (m, 1H, H-3b), 2.63 – 2.81 (m, 1H, H-9a), 3.06 (dd, *J*=13.4, 3.1, 1H, H-9b), 3.45 (dd, *J*=13.4, 8.6, 1H, H-7), 3.74 (tt, *J*=8.8, 8.8, 4.7, 4.7, 1H, H-8), 3.78 – 3.96 (m, 1H, H-5), 3.96 – 4.09 (m, 2H, H-6, H-4). $^{13}$C NMR (101 MHz, $D_2O$) 22.05 (**C**H$_3$CO), 39.35 (C-3), 43.54 (C-9), 52.21 (C-5), 67.24 (C-4), 69.89 (C-8), 70.15 (C-6, C-7), 96.38 (C-2), 174.69 ($CH_3$**C**O), 176.65 (COOH).

4. 2TCO-NHS active ester (TCO*NHS-carbonate/axial isomer, **5**, from SiChem, SC-8070) (0.5g; 1.9 mmol) at 0°C was added to compound **4** (0.64 g; 2.1 mmol) dissolved in dry dimethylformamide (18 mL), followed by DIPEA (1.7 mL; 4.6 mmol). The reaction mixture was

stirred at room temperature for 1.5 hours. The reaction was quenched by the addition of water (4 mL), and the reaction mixture was stirred at room temperature for 15 minutes. The MeOH (10 ml) was added, the solvents were evaporated under reduced pressure, and the residue was purified by C18 reverse phase HPLC chromatography (using a gradient of $CH_3CN$ in water 5→95%, both containing 0.05% HCOOH). The product Sia-2TCO (**6**) was obtained as a light white solid (0.63 g; 66 %) after lyophilization (mixture of anomers). $^1$H NMR (600.1 MHz, $CD_3OD$): 0.83 – 0.91 (m, 1H, H-6b-cyclooctene); 1.13 – 1.20 (m, 1H, H-7b-cyclooctene); 1.46 – 1.53 (m, 1H, H-5b-cyclooctene); 1.60 – 1.68 (m, 1H, H-7a-cyclooctene); 1.69 – 1.76 (m, 1H, H-8b-cyclooctene); 1.82 (dd, 1H, $J_{gem}$ = 12.8, $J_{3b,4}$ = 11.6, H-3b); 1.83 – 1.90 (m, 1H, H-6a-cyclooctene); 1.95 – 2.07 (s, 6H, $CH_3CO$, H-4b,5a,8a-cyclooctene); 2.21 (dd, 1H, $J_{gem}$ = 12.0, $J_{3a,4}$ = 4.9, H-3a); 2.42 – 2.48 (m, 1H, H-4a-cyclooctene); 3.15 – 3.32 (m, 1H, H-9b); 3.39 (bd, 1H, $J_{7,8}$ = 9.5, H-7); 3.546, 3.552 (2 × dd, 2 × 1H, $J_{gem}$ = 14.2, $J_{9a,8}$ = 5.3, H-9a); 3.71 – 3.76 (m, 1H, H-8); 3.84 (t, 1H, $J_{5,4}$ = $J_{5,6}$ = 10.4, H-5); 4.012, 4.014 (2 × dd, 2 × 1H, $J_{6,5}$ = 10.4, $J_{6,7}$ = 1.3, H-6); 4.03 (ddd, 1H, $J_{4,3}$ = 11.6, 4.9, $J_{4,5}$ = 10.4, H-4); 5.23 – 5.27 (m, 1H, H-1-cyclooctene); 5.55 (dd, 1H, $J_{2,3}$ = 16.4, $J_{2,1}$ = 2.2, H-2-cyclooctene); 5.82 – 5.89 (m, 1H, H-3-cyclooctene). $^{13}$C NMR (150.9 MHz, $CD_3OD$): 22.70 ($CH_3CO$); 25.20, 25.21 ($CH_2$-7-cyclooctene); 30.09 ($CH_2$-6-cyclooctene); 36.79 ($CH_2$-4-cyclooctene); 37.03 ($CH_2$-5-cyclooctene); 40.99, 41.00 ($CH_2$-3); 41.63, 41.66 ($CH_2$-8-cyclooctene); 45.40, 45.46 ($CH_2$-9); 54.24 (CH-5); 67.89 (CH-4); 70.84, 70.89 (CH-8); 71.49, 71.54 (CH-7); 72.08, 72.10 (CH-6); 75.30, 75.31 (CH-1-cyclooctene); 96.61 (C-2); 132.61, 132.64 (CH-3-cyclooctene); 132.81, 132.85 (CH-2-cyclooctene); 159.08 (OCON); 173.46 ($CH_3CO$); 174.93 (C-1). HRMS $[M-H]^-$ m/z calcd. for $[C_{20}H_{31}O_{10}N_2]^-$ 459.19842, found 459.19775.

#### Preparation of (E)-5-Acetamido-9-{[(R,E)-(cyclookt-4-en-1-yl)oxycarbonyl]amino}-3,5,9-trideoxy-D-glycero-D-galacto-non-2-ulopyranosic acid (Sia-4TCO)

TCO4-NHS active ester (TCO4-NHS carbonate/axial isomer, **7**, from SiChem SC-8072; 32 mg; 0.12 mmol) at 0°C was added to compound **4** (37 mg; 0.12 mmol) dissolved in dry dimethylformamide (2 mL), followed by DIPEA (78 µL; 0.45 mmol). The reaction mixture was stirred at room temperature for 1 hour. The reaction was quenched by the addition of water (3.5 mL), filtered, and the solution subjected to preparative HPLC (using a gradient of $CH_3CN$ in water, both containing 0.05% HCOOH). The product TCO4eq-Sia (**8**) was obtained as a colorless solid after lyophilization (mixture of anomers; 23 mg; 42 %). $^{1}$H NMR (401 MHz, MeOD): 1.60 (dq, *J* = 15.7, 6.4, 6.4, 5.5 Hz, 1H, H-8a'), 1.66 – 1.77 (m, 2H, H-8b', H-3'), 1.83 (dd, *J* = 12.8, 11.3 Hz, 1H, H-3a), 1.87 – 2.00 (m, 2H, H-3', H-6'), 2.01 (s, 3H, $CH_3$), 2.21 (dd, *J* = 12.8, 4.9 Hz, 1H, H-3b), 2.26 – 2.39 (m, 3H, H-6b', H-7b'), 3.10 – 3.20 (m, 1H, H-9a), 3.33 – 3.40 (m, 1H, H-7), 3.52 (dt, *J* = 14.0, 3.1, 3.1 Hz, 1H, H-9b), 3.71 (ddd, *J* = 9.0, 7.2, 3.2 Hz, 1H, H-8), 3.83 (t, *J* = 10.2, 10.2 Hz, 1H, H-5), 3.96 – 4.08 (m, 2H, H-4, H-6), 4.32 (s, 1H, H-1'), 5.48 (ddd, *J* = 16.0, 10.7, 3.5 Hz, 1H, H-4,), 5.53 – 5.73 (m, 1H, H-5'). $^{13}$C NMR (101 MHz, MeOD): 22.69 ($CH_3$), 32.11 (C-3'), 33.48 (C-7'), 35.17 (C-6'), 39.60 (d, *J* = 5.8 Hz, C-8'), 41.00 (C-3), 42.19 (C-2'), 45.40 (C-9), 54.22 (C-5), 67.91 (C-4), 70.82 (C-8), 71.52 (C-7), 72.08 (C-6), 81.90 (H-1'), 96.61 (C-2), 133.79 (C-4'), 136.10 (C-5'), 159.35 (CO-9), 173.52 (COOH), 174.90 (CO-5). HRMS [M-H]$^{-}$ m/z calcd. for $[C_{20}H_{31}O_{10}N_2]^-$ 459.19842, found 459.19883.

#### (E)-5-Acetamido-9-{[(2-methylcycloprop-2-en-1-yl)methoxycarbonyl]amino}-3,5,9-trideoxy-D-glycero-D-galacto-non-2-ulopyranosic acid (Sia-Cp, 10)

#### (E)-5-Acetamido-9-[(2-methylcycloprop-2-en-1-yl)methoxycarbonylamino]-3,5,9-trideoxy-D-glycero-D-galacto-non-2-ulopyranosic acid (Sia-Cp, 10)

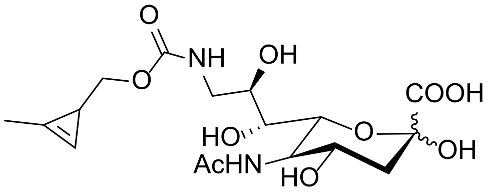


Methylcyclopropene-OpNP active ester **9** (15 mg; 60.2 µmol) at 0°C was added to compound **4** (28 mg; 90.3 µmol) dissolved in dry dimethylformamide (1.8 mL), followed by DIPEA (31 µL; 180.6 µmol). The reaction mixture was stirred at room temperature for 5.5 hours. The reaction mixture was filtered and loaded directly on C18 reverse phase HPLC chromatography column (using a gradient of $CH_3CN$ in water 0→55%). The product Sia-Cp (**10**) was obtained as a light white solid (8.9 mg; 35 %) after lyophilization (mixture of anomers). $^{1}$H NMR(MeOD, 500 MHz): 1.62 (td, 1H, *J*=5.1, 5.1, 1.5 Hz, H-1Cp), 1.88 (t, 1H, *J*=11.8, 11.8 Hz, H-3a), 2.00 (s, 3H, NH**Ac**),

1.99 – 2.08 (m, 2Hz), 2.13 (d, 5H, *J*=1.2 Hz, H-3b, $CH_3Cp$), 3.14 (dd, 1H, *J*=11.1, 5.4 Hz, H-9a), 3.35 (dt, 1H, *J*=10.0, 4.1, 4.1 Hz, H-8), 3.51 (q, 1H, *J*=13.9, 13.9, 11.7 Hz, H-9b), 3.70 (d, 1H, *J*=12.2 Hz, H-8), 3.82 (dt, 1H, *J*=10.8, 5.2, 5.2 Hz, CH2-Cpa), 3.87 – 4.03 (m, 4H, CH2-Cpb, H-4, H-5, H-6), 4.60 (s, 2H), 4.83 (d, 1H, *J*=1.8 Hz), 4.91 (s, 1H) artefakt?, 6.65 (s, 1H, H-3Cp). $^{13}C$ NMR (126 MHz, MeOD) 11.59 ($CH_3Cp$), 18.30 (C-1Cp), 22.85 ($CH_3NH$), 41.74 (C-3), 45.47 (C-9), 49.07, 54.05 (C-5), 68.83 (C-6), 71.15 (C-8), 71.53 (C-7), 71.90 (C-4), 73.45 (C-3), 97.77 (C-2), 102.92 (C-3Cp), 122.26 (C-2Cp), 159.92 (COCp), 174.34 (N**C**$OCH_3$), 177.46 (COOH). HRMS $[M-H]^-$ m/z calcd. for $[C_{17}H_{25}O_{10}N_2]^-$ 417.15117, found 417.15147.

#### (E)-5-Acetamido-9-{[((1S,4S)-bicyclo[2.2.1]hept-5-en-2-yl)methoxycarbonyl]amino}-3,5,9-trideoxy-D-glycero-D-galacto-non-2-ulopyranosic acid (Sia-Norb, 12)

#### (E)-5-Acetamido-9-[(bicyclo[2.2.1]hept-5-en-2-yl)methoxycarbonylamino]-3,5,9-trideoxy-D-glycero--D-galacto-non-2-ulopyranosic acid (Sia-Norb, 12)

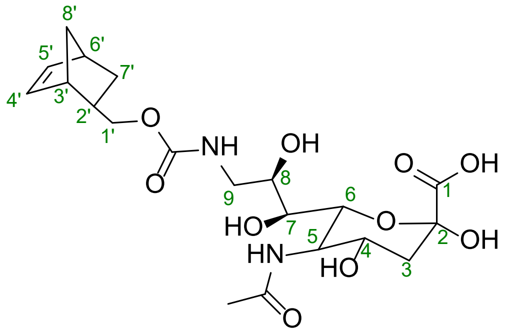


Endo/exo (5:6) norbornene-NHS active ester, prepared as described in lit.[3] (**11**, 35 mg; 131 µmol) was added at 0°C to compound **4** (45 mg; 146 µmol) dissolved in dry dimethylformamide (2.6 mL). DIPEA (114 µL; 657 µmol) was then added to the mixture. The reaction mixture was stirred at room temperature for 1 hour. The reaction mixture was filtered and loaded directly on C18 reverse phase HPLC chromatography column (using a gradient of $CH_3CN$ in water 20→60%). The product Sia-Norb (**12**) was obtained as a light white solid (21 mg; 31 %) after lyophilization (mixture of anomers, mixture of endo/exo norbornene 3:2). $^1H$ NMR (400 MHz, MeOD) 0.55 (ddd, *J*=11.7, 4.5, 2.7, 1H, 7'a endo), 1.14 – 1.40 (m, 3H, 7'a exo, 7'b exo, 8'a endo, 8'exo), 1.44 (dd, *J*=8.1, 2.2, 1H, 8'b endo), 1.70 (s, 1H, 2' exo), 1.78 – 1.90 (m, 2H, H-3a, 7'b endo), 2.01 (s, 4H, $CH_3$), 2.06 (s, 0H), 2.21 (dd, *J*=12.8, 4.9, 1H, H-3b), 2.40 (s, 1H, 2' endo), 2.71 (s, 1H, 3' exo), 2.80 (d, *J*=6.4, 2H, 6'), 2.88 (s, 1H, 3' endo), 3.13 – 3.21 (m, 1H, H-9a), 3.29 – 3.42 (m, 2H, H-7), 3.49 – 3.67 (m, 1H, H-9b, 1'a endo), 3.73 (td, *J*=7.2, 6.9, 3.6, 1H, H-8), 3.78 – 3.89 (m, 2H, H-5, 1'b endo), 3.94 (td, *J*=10.2, 9.8, 5.6, 1H,

1’a exo), 3.99 – 4.06 (m, 2H, H-4, H-6), 4.12 (dt, *J*=11.2, 5.7, 5.7, 1H, 1’b exo), 5.95 (d, *J*=6.0, 1H, 4’endo), 6.09 (dd, *J*=2.4, 1.2, 1H, 4’exo, 25’ exo), 6.16 (dd, *J*=5.7, 3.1, 1H, 5’endo). $^{13}$C NMR (101 MHz, MeOD) 22.70 ($CH_3$), 29.77 (7’endo), 30.34 (7’exo), 39.44 (2’endo), 39.73 (2’exo), 41.02 (C-3), 42.80 (6’exo), 43.45 (6’endo), 44.81 (3’exo), 45.06 (3’endo), 45.48 (C-9), 45.78 (8’exo), 49.50, 50.29 (8’endo), 54.23 (C-5), 67.92 (C-4), 69.39 (1’endo), 70.04 (1’exo), 70.80 (C-8), 71.56 (C-7), 72.09 (C-6), 96.66 (C-2), 133.22 (4’ endo), 137.30 (4’exo), 137.94 (5’exo), 138.45 (5’endo), 159.82 (C=O Norb), 173.62 (COOH), 174.93 ($CH_3$*C*O). HRMS $[M-H]^-$ m/z calcd. for $[C_{20}H_{29}O_{10}N_2]^-$ 457.18251, found 457.18277.

#### 1,3,4,6-Tetra-O-acetyl-2-{[(R,E)-(cyclookt-4-en-1-yl)oxycarbonyl]amino}-2-deoxy-α-D-mannopyranose (Ac4Man-4TCO, 15)

Overit, jestli mame smes anomeru, nebo jenom alpha (schema)!- dle Radka pouze Alpha

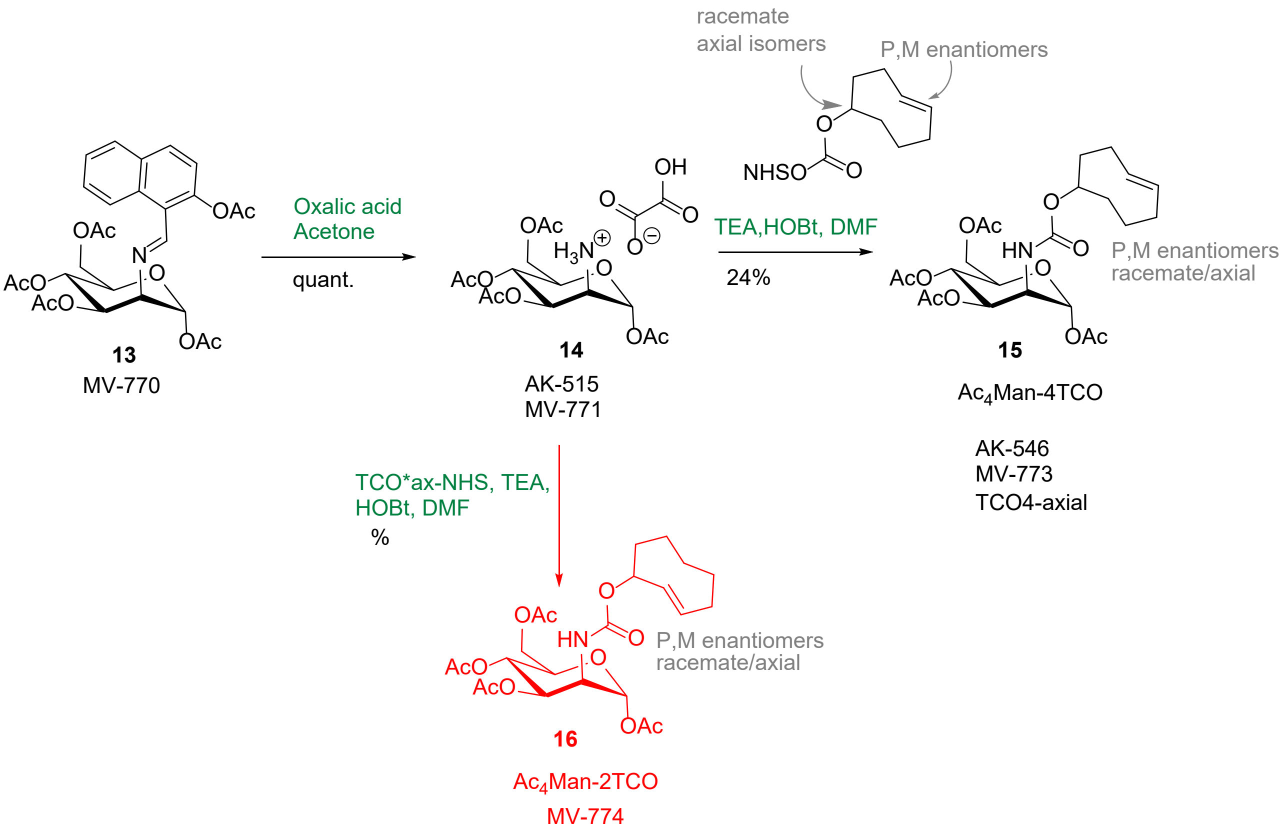


1. Ac4Man-4TCO (**15**) preparation: Solution of **13**[4] (150 mg; 0.3 mmol) in acetone (4.5 ml) was cooled to 0°C. Oxalic acid dihydrate (157 mg; 1.2 mmol) was added as solid, followed by acetone

(3 ml). The mixture was stirred for 35 min at 0°C and 1h at room temperature. The mixture was cooled again, filtered off, and the white solids were washed with cold acetone (2x) and diethyl ether (3x). After drying, solid **14** was subjected to the next step (73 mg; 61 %). $^1$H NMR (400 MHz, MeOD): 2.06 (s, 3H, $CH_3CO$), 2.07 (s, 3H, $CH_3CO$), 2.12 (s, 3H, $CH_3CO$), 2.20 (s, 3H, $CH_3CO$), 3.92 (dd, *J* = 4.7, 1.8 Hz, 1H, H-2), 4.10 (dd, *J* = 12.3, 2.6 Hz, 1H, H-6a), 4.23 (ddd, *J* = 8.8, 5.6, 2.6 Hz, 1H, H-4), 4.34 (dd, *J* = 12.3, 5.7 Hz, 1H, H-6b), 5.37 (t, *J* = 9.9, 9.9 Hz, 1H, H-4), 5.49 (dd, *J* = 9.9, 4.7 Hz, 1H, H-3), 6.27 (d, *J* = 1.7 Hz, 1H, H-1).$^{13}$C NMR (101 MHz, MeOD): 20.47, 20.50, 20.52, 20.58 (**C**$H_3$CO-1,3,4,6), 52.31(C-2), 63.29 (C-6), 66.26 (C-4), 68.91 (C-3), 71.72 (C-5), 90.89 (C-6), 126.51, 147.99, 165.03, 169.51 ($CH_3$**C**O-1), 170.94 ($CH_3$**C**O-3), 171.27 ($CH_3$**C**O-4), 172.31 ($CH_3$**C**O-6). HRMS $[M+H]^+$ m/z calcd. for $[C_{14}H_{21}O_9NNa]^+$ 370.11085, found 370.11059.

2. The crude product **14** from the previous step (70 mg; 0.16 mmol) and 4TCO-NHS (TCO4-NHS carbonate/axial isomer, from Sichem, SC-8072, 42 mg; 0.16 mmol) in dry DMF (2 ml) were cooled to 0°C. TEA (122 µl; 0.88 mmol) was added, than HOBt (solid, 22 mg; 0.16 mmol ) was added, the mixture was stirred 10 min at 0°C and 2 h at room temperature. The mixture was diluted with acetonitrile/water (1 : 1) to total volume of 3.5 ml, filtered and loaded onto a C18 HPLC column, where it was purified (using a gradient of $CH_3CN$ in water, both containing 0.05% HCOOH). The $Ac_4$Man-4TCO **(15)** was isolated as a white solid (19 mg; 24 %) after lyophilization. $^1$H NMR (400.1 MHz, $CD_3OD$): 1.24 – 1.34 (m, 2H, H-8′b); 1.53 – 1.81 (m, 6H, H-2′b,7′); 1.81 – 1.91 (m, 2H, H-6′b); 1.97, 1.99 (2 × s, 2 × 3H, $CH_3CO$-3); 2.060, 2.064 (2 × s, 2 × 3H, $CH_3CO$-6); 2.069, 2.071 (2 × s, 2 × 3H, $CH_3CO$-4); 2.07 – 2.13 (m, 2H, H-3′b); 2.17 (s, 6H, $CH_3CO$-1); 2.21 – 2.35 (m, 6H, H-2′a,6′a,8′a); 2.38 – 2.53 (m, 2H, H-3′a); 4.06 – 4.14 (m, 4H, H-5,6b); 4.26 – 4.36 (m, 4H, H-2,6a); 4.79 – 4.86 (m, 2H, H-1′); 5.25, 5.27 (2 × dd, 2 × 1H, $J_{3,4}$ = 10.1, $J_{3,2}$ = 4.5, H-3); 5.35, 5.36 (2 × t, 2 × 1H, $J_{4,3}$ = $J_{4,5}$ = 10.1, H-4); 5.56, 5.57 (2 × ddd, 2 × 1H, $J_{4',5'}$ = 16.0, $J_{4',3'}$ = 11.2, 3.2, H-4′); 5.77, 5.80 (2 × ddd, 2 × 1H, $J_{5',4'}$ = 16.0, $J_{5',6'}$ = 7.0, 3.7, H-5′); 5.97, 5.98 (2 × d, 2 × 1H, $J_{1,2}$ = 1.8, H-1). $^{13}$C NMR (100.6 MHz, $CD_3OD$): 20.62, 20.63, 20.64, 20.68, 20.74 (**C**$H_3$CO-1,3,4,6); 29.01, 29.16 (C-7′); 30.81, 30.82 (C-3′); 33.66, 33.70 (C-8′); 35.25, 35.28 (C-6′); 41.93, 41.94 (C-2′); 52.32, 52.39 (C-2); 63.86, 64.01 (C-6); 67.03, 67.27 (C-4); 70.92, 71.14 (C-3); 71.85, 71.89 (C-5); 72.25, 72.37 (C-1′); 93.73, 93.77 (C-1); 132.51, 132.53 (C-4′); 136.53 (C-5′); 158.42, 158.54 (NCOO); 170.10, 170.12 ($CH_3$**C**O-1); 171.46, 171.54

($CH_3$**C**O-4); 171.64, 171.69 ($CH_3$**C**O-3); 172.43 ($CH_3$**C**O-6). NMR není v souladu s lit-Zhang a spol[5]

HRMS (ESI): m/z calculated for $C_{23}H_{33}O_{11}NNa$ = 522.19458 $[M+Na]^+$, found: 522.19470

#### 1,3,4,6-Tetra-O-acetyl-2-{[(R,E)-(cyclookt-2-en-1-yl)oxycarbonyl]amino}-2-deoxy-α-D-mannopyranose (Ac4Man-2TCO, 16)

Overit, jestli mame smes anomeru, nebo jenom alpha (schema)!

$Ac_4$Man-2TCO **16** was prepared in analogy to $Ac_4$Man-4TCO using the 2TCO-NHS active ester (TCO*NHS-carbonate/axial isomer, **5**, from SiChem, SC-8070) with the following modification: 2TCO-NHS (61 mg, 1 equiv.) was added in portions to a suspension of **13** (100 mg, 0.23 mmol) in dry DCM (5 mL) cooled down in ice-water bath. DIPEA (0.2 mL, 5 equiv.) was added dropwise to this mixture, which was allowed to warm to room temperature. After 72 hours, TLC analysis (DCM/AcOEt = 9/1, $KMnO_4$ staining) indicated the reaction finished. Solvents were removed under reduced pressure and the residue was purified by column chromatography using DCM to DCM/AcOEt = 9/1 gradient. Fractions containing the product were pooled, solvents were evaporated, and the residue was redissolved in $CH_3CN/H_2O$ = 1/1 and lyophilized to give the title compound **16** as white solid (20 mg, 18 %).

$^1$H NMR (401 MHz, MeOD): 0.89 (ddd, $J$ = 19.6, 11.0, 5.8 Hz, 1H, H-6’), 1.17 – 1.32 (m, 2H, H-7a’), 1.53 (tdd, $J$ = 16.6, 16.6, 10.4, 3.9 Hz, 1H, H-5a’), 1.60 – 1.80 (m, 1H, H-7b’), 1.89 (ddt, $J$ = 16.3, 11.3, 3.5, 3.5 Hz, 1H, H-8a’), 1.95 (s, 3H, Ac-6), 1.98 (s, 1H, H-4a’), 2.04 – 2.08 (m, 7H, Ac-3, H-5b’, H-8b’, Ac-4), 2.17 (s, 3H, Ac-6), 2.47 (dd, $J$ = 10.3, 4.6 Hz, 1H, H-4b’), 4.04 – 4.13 (m, 2H, H-5, H-6a), 4.31 (ddt, $J$ = 9.6, 7.5, 3.7, 3.7 Hz, 2H, H-2, H-6b), 5.17 – 5.38 (m, 3H, H-1’, H-3, H-4), 5.56 (ddd, $J$ = 16.4, 9.7, 2.3 Hz, 1H, H-2’), 5.84 – 5.95 (m, 1H, H-3’), 5.96 (t, $J$ = 2.0, 2.0 Hz, 1H, H-1).

$^{13}$C NMR (101 MHz, MeOD): 20.39 – 21.10 (m, **C**$H_3$CO), 25.25 (d, $J$ = 11.0 Hz, C-7’), 30.07 (d, $J$ = 3.7 Hz, C-6’), 36.75 (d, $J$ = 7.3 Hz, C-4’), 37.00 (C-5’), 41.48 (d, $J$ = 18.7 Hz, C-8’), 52.34 (C-

2), 63.95 (C-6), 67.14 (C-3), 70.92 (C-4), 71.83 (C-5), 75.92 (C-1’), 93.81 (C-1), 132.51 (C-3’), 132.90 (d, $J$ = 14.3 Hz, C-4’), 158.30 (**C**ONH), 170.09 (CO-1), 171.50, 171.64 (CO-3), 171.80 (CO-4), 172.46 (CO-6).

HRMS (ESI): m/z calculated for $C_{23}H_{33}O_{11}NNa$ = 522.19458 $[M+Na]^+$, found: 522.19467

#### Synthesis of HA-Tag-Tz peptide

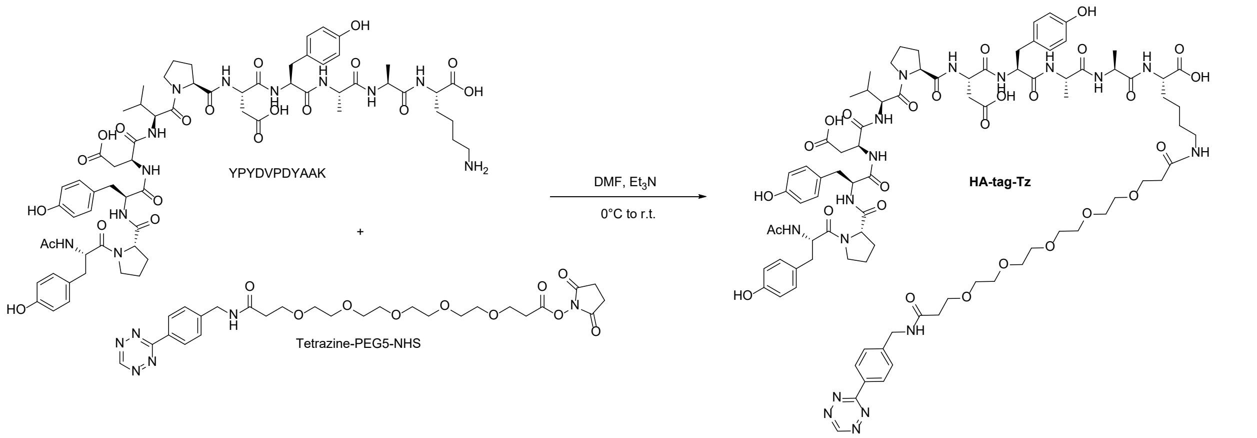


The HA-tag peptide sequence with alanine-lysine at the C terminus and acetylated at the N-terminus (Ac-YPYDVPDYAAK) was synthesized on trityl resin using standard Fmoc procedure. The cleaved and deprotected peptide (10 mg) was dissolved in dry DMF (0.4 mL) and tetrazine-PEG5-NHS (5.2 mg, 1.15 equiv., Conju-probe CP-6025) was added. The solution was cooled down in ice-water bath and $Et_3N$ (5 µL, 4.8 equiv.) was added. The reaction was stirred at room temperature and progress was followed by LC-MS. After ca. 1 hour, the reaction was quenched by diluting it with a 1:1 mixture of $CH_3CN/H_2O$ containing 0.05% formic acid (ca. 1 mL). This solution was filtered and purified by RP-HPLC using a gradient of $CH_3CN$ in $H_2O$ (5-60% over 20 min) containing 0.1% TFA to yield HA-Tag-Tz peptide as a pink solid (10 mg, 73%) after lyophilization.

#### Preparation of FLAG-Tag-Tz

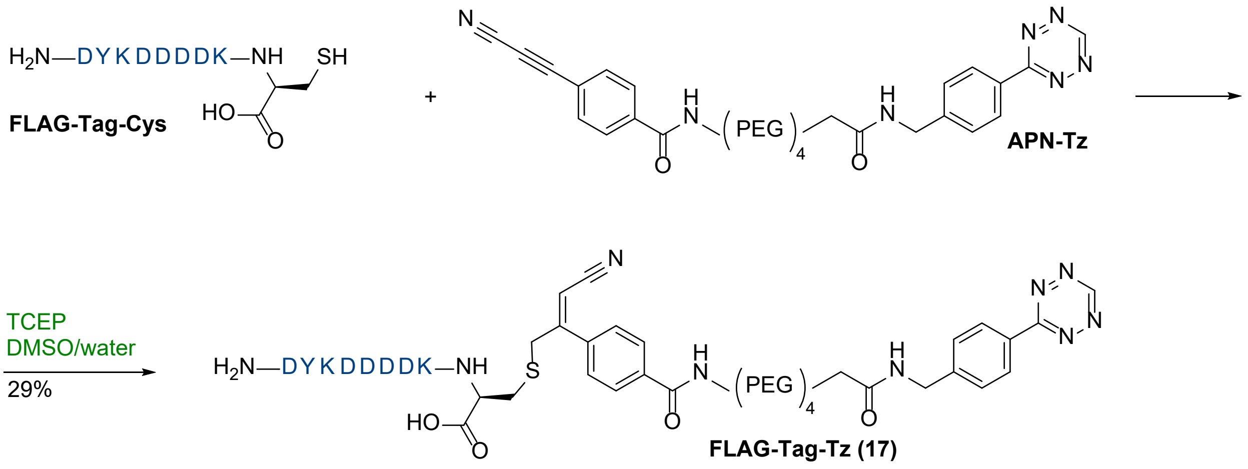


FLAG-Tag-Tz (**17**) preparation was based on lit [8]. Peptide FLAG-Tag-Cys (DYKDDDDKC, prepared by Synpeptide.com) (7.2 mg; 6.5 µmol in water/PBS = 1:1, 646 µL) was mixed with the solution of APN-Tz (3.8 mg; 6.5 mmol in DMSO, 430 µL, from Conju Probe CP-8010). The vial was flushed with argon and shaken at 37°C for 3h. TCEP (0.74 mg; 2.6 µmol in water, 65 µL) was then added, DMSO (325 µL) was added, the mixture was shaken at 37°C for 2h. An additional portion of APN-Tz (0.35 mg; 0.6 µmol in water, 40 µL) and DMSO (200 µL) were added, shaken 37°C for 3h. The mixture was diluted with water to a total volume of 3.5 ml, filtered and loaded onto a C18 HPLC column, where it was purified (using a gradient of $CH_3CN$ in water, both containing 0.05% HCOOH). The product FLAG-Tag-Tz (**17**) was obtained as a pink solid after lyophilization (3.2 mg, 29 %). HRMS $[M-H]^-$ m/z calcd. for $[C_{74}H_{97}O_{27}N_{18}S]^-$ 1701.64842, found 1701.64967.

#### Fluorogenic properties of Tz-AF488 and Coum-Tz

The turn-on fluorescence measurements were performed as follows: 1 mM solution of tetrazines (1.0 µL) in DMSO (100%) (final concentration 1 µM) was diluted in 1 mL PBS (100%) and solution of Sia-2TCO (5 µL from 10 mM fresh stock in 100% PBS) was added. The cuvette was inserted into fluorescence spectrophotometer and the measurement was started. All probes were excited in accordance with their characteristic absorption (405 nm for Coum-Tz and 490 nm for Tz-AF488) and the fluorescence was collected between 415-650 nm for Coum-Tz and from 500-700 nm for Tz-AF488. The spectra were recorded at several time points and all measurements were typically repeated three-four times. The data were processed using OriginPro software. All spectra were subtracted from the baseline (PBS (100%) as the blank). The data were finally processed, and the turn-on values were calculated from the observed fluorescence intensities of the click products at the emission maximum divided by the highest residual fluorescence of the quenched tetrazine heterodienes. These values are presented in Figure S2.

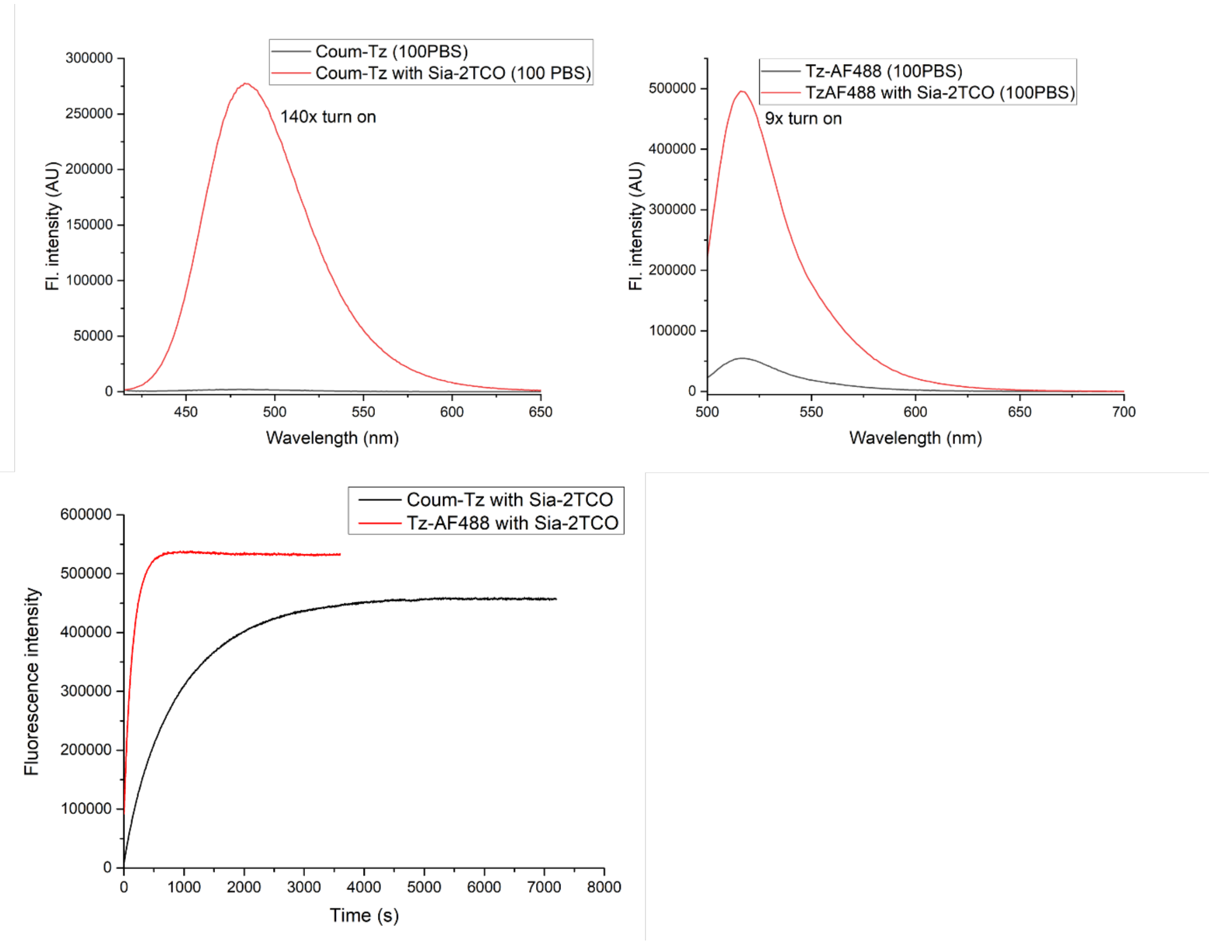


*Figure S2.* Fluorescence spectra of the click products formed in the reaction of fluorogenic tetrazine Coum-Tz or Tz-AF488 with Sia-2TCO (indicated as number x = x-fold) in (100%) PBS.

### NMR spectra

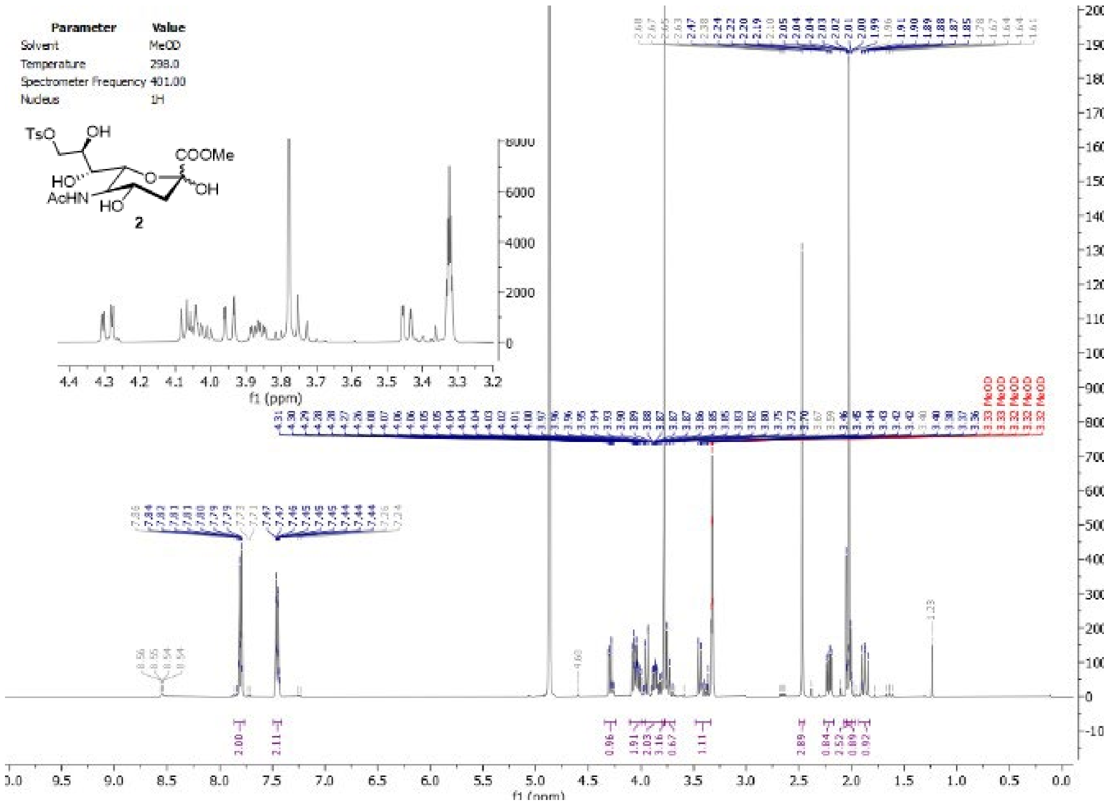
Parameter Value
Solvent MeOD
Temperature 298.0
Spectrometer Frequency 401.00
Nucleus 1H
TsO
OH
COOMe
HO
OH
AcHN
HO
2
f1 (ppm)

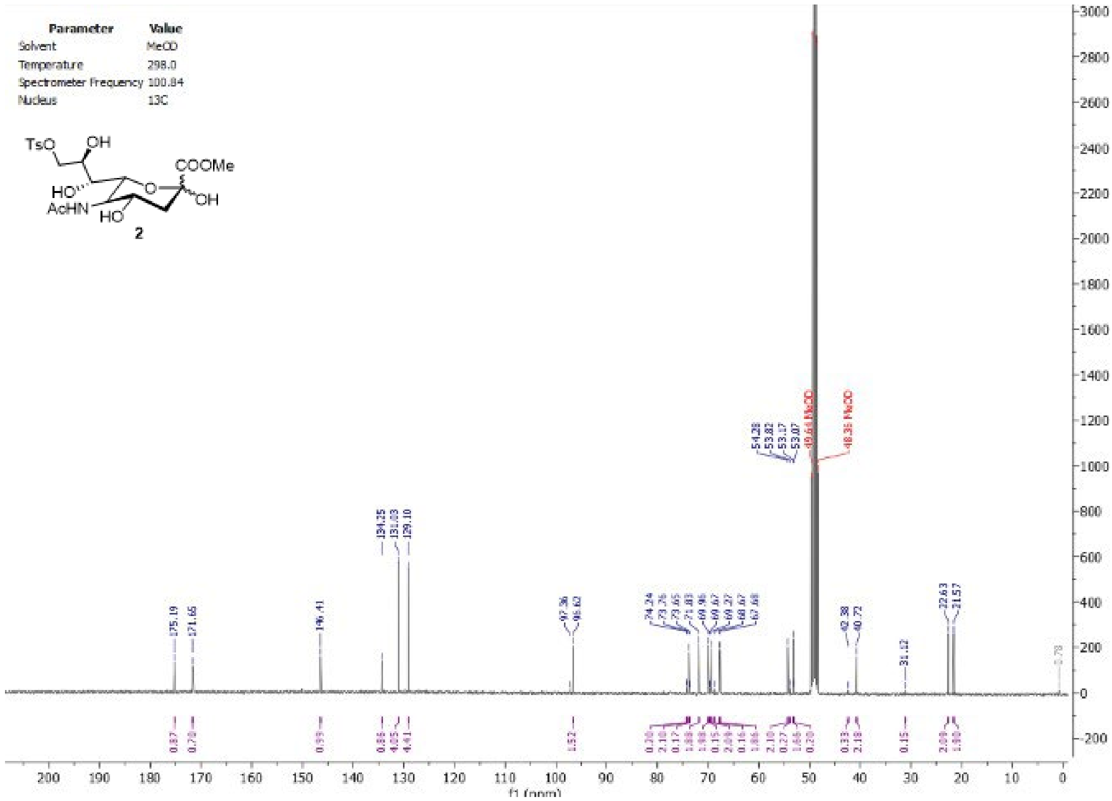
Parameter Value
Solvent MeOD
Temperature 298.0
Spectrometer Frequency 100.84
Nucleus 13C
TsO
OH
COOMe
HO
OH
AcHN
HO
2
f1 (ppm)

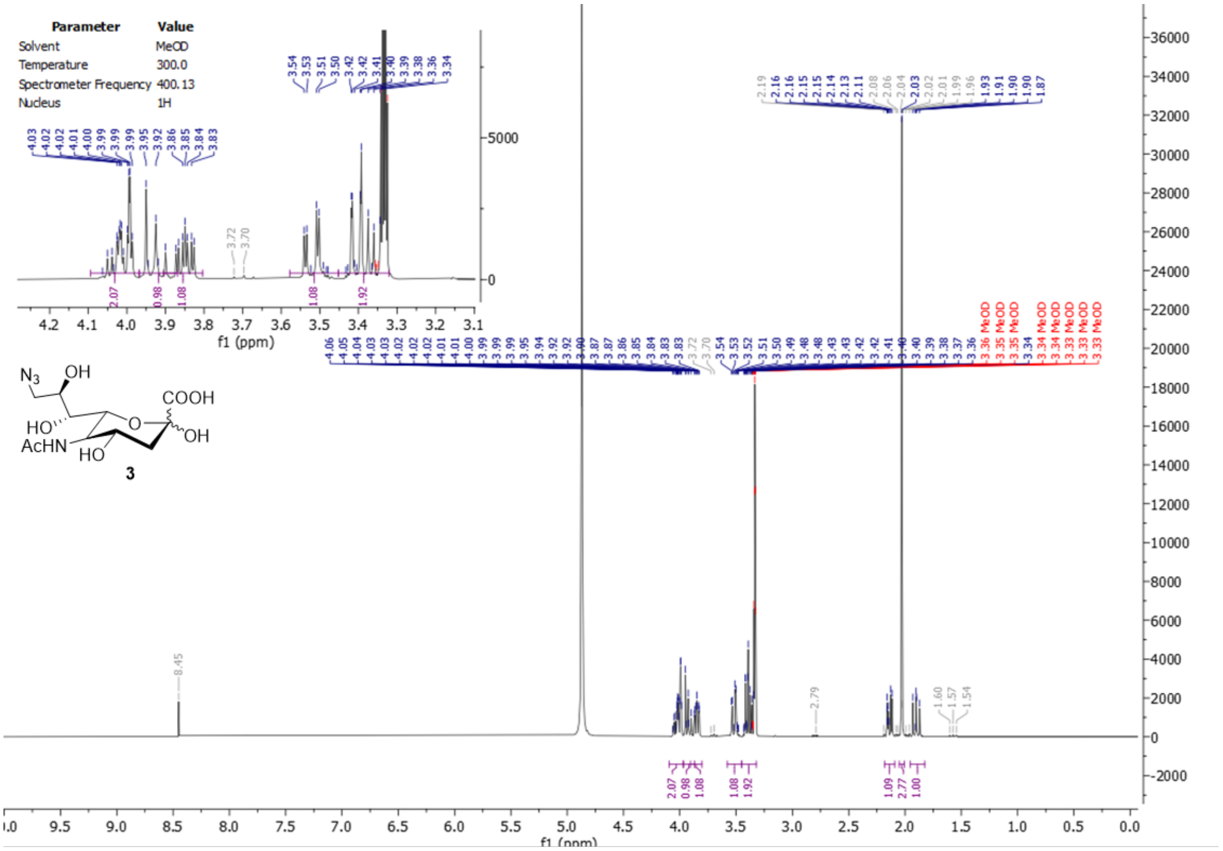

Parameter Value
Solvent MeOD
Temperature 300.0
Spectrometer Frequency 400.13
Nucleus 1H
3
f1 (ppm)


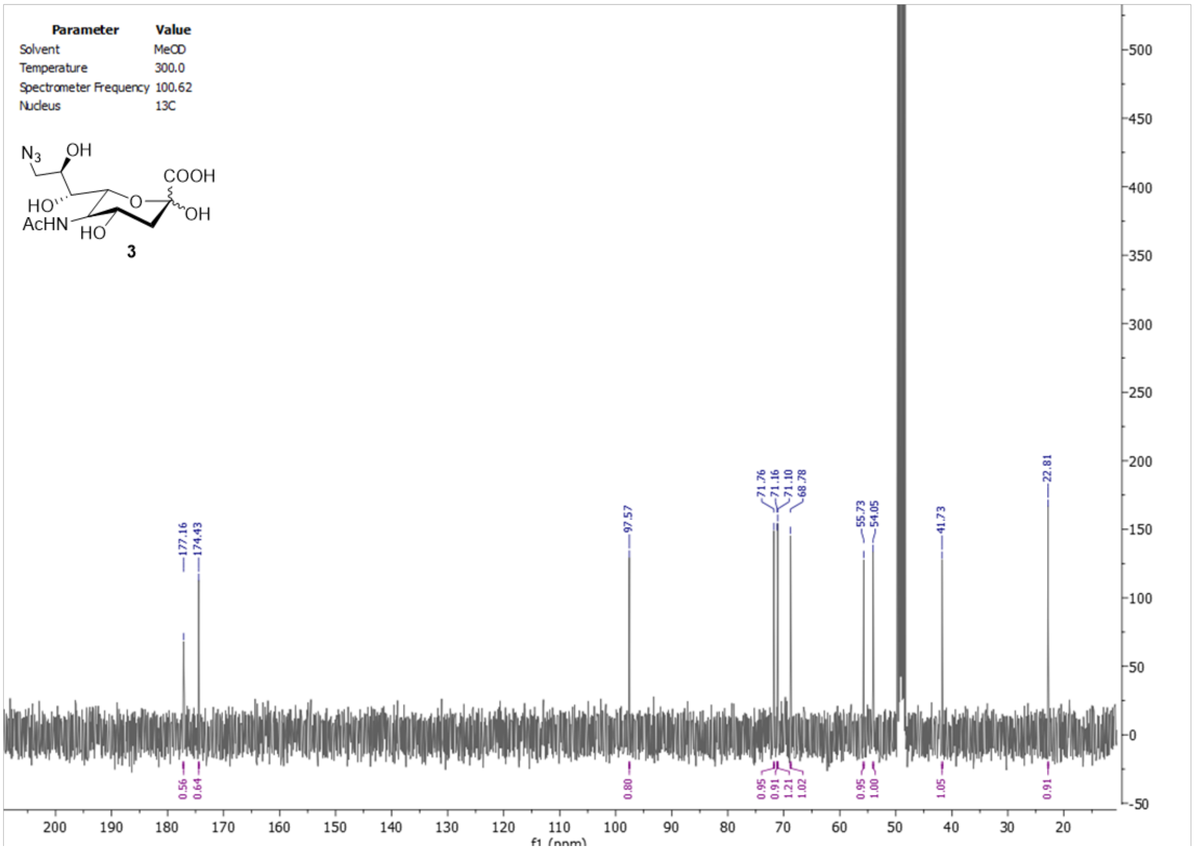

Parameter Value
Solvent MeOD
Temperature 300.0
Spectrometer Frequency 100.62
Nucleus 13C
3
177.16
174.43
97.57
71.76
71.16
71.10
68.78
55.73
54.05
41.73
22.81
f1 (ppm)

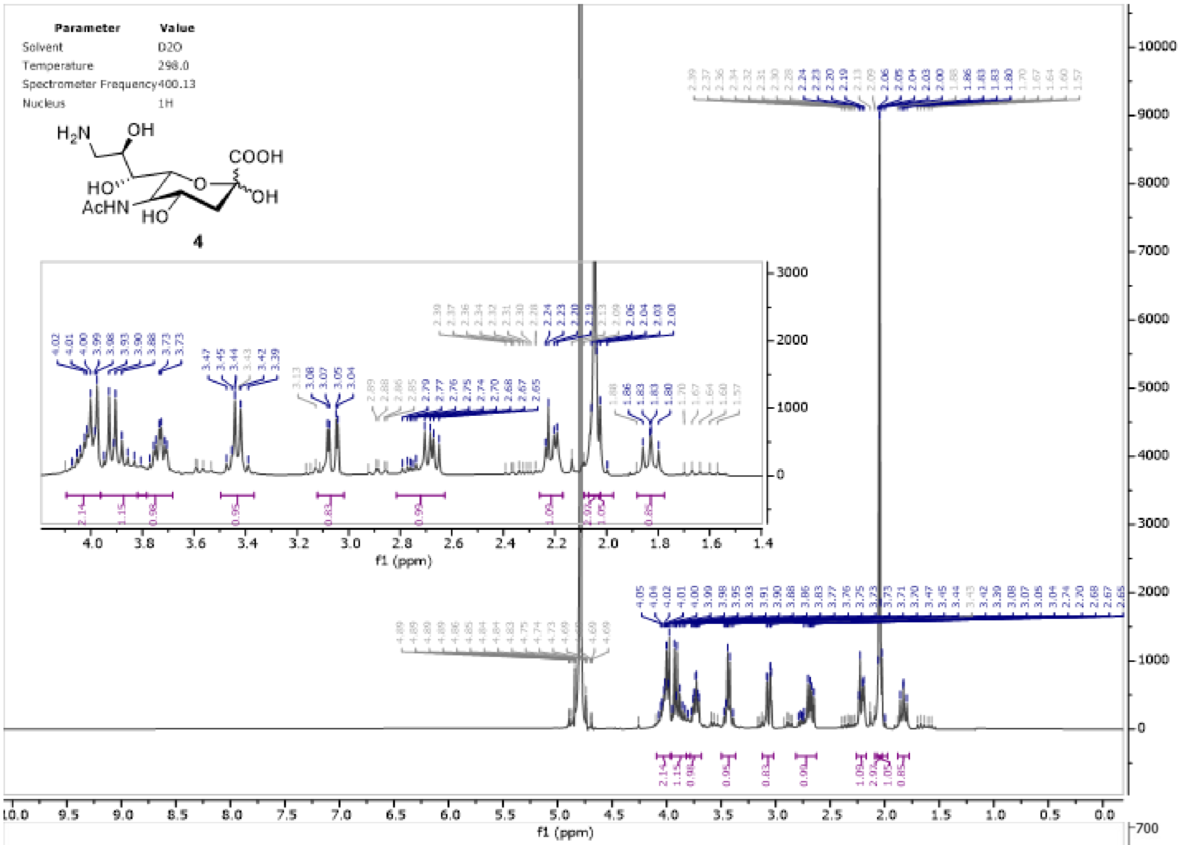
Parameter Value
Solvent D2O
Temperature 298.0
Spectrometer Frequency 400.13
Nucleus 1H
4
f1 (ppm)

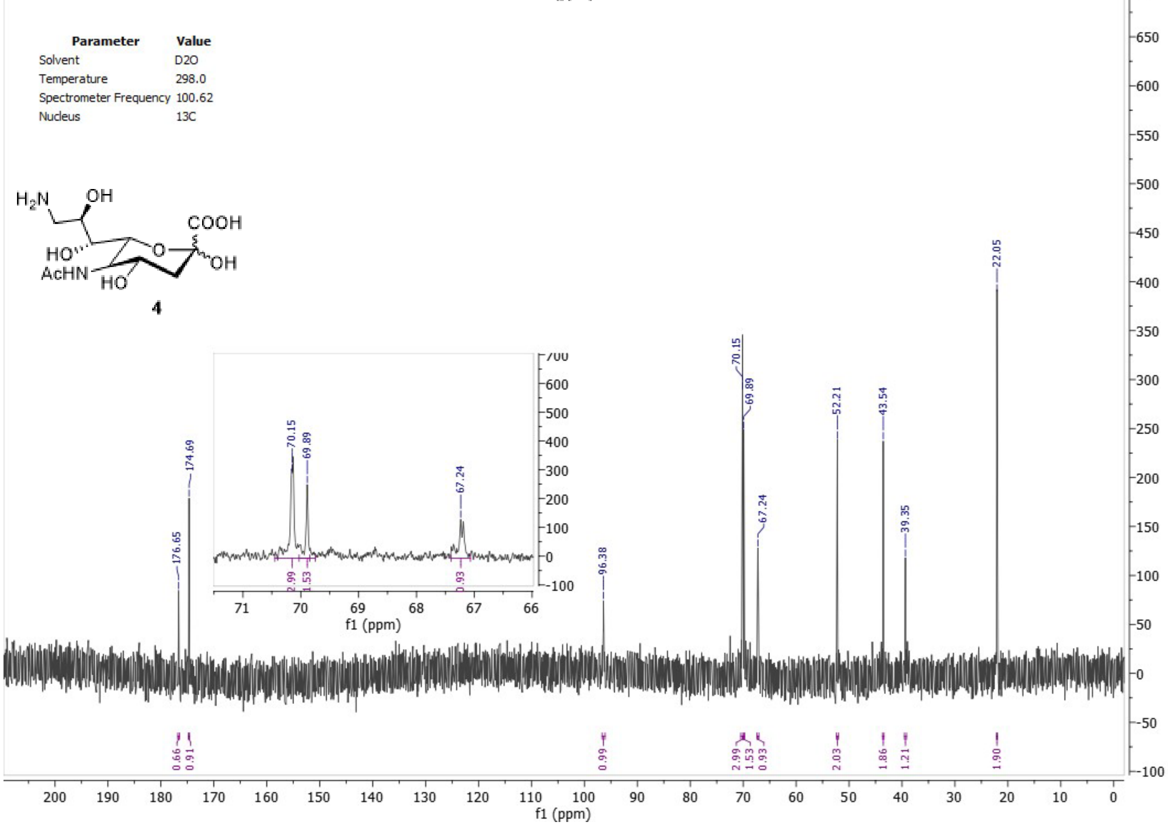
Parameter Value
Solvent D2O
Temperature 298.0
Spectrometer Frequency 100.62
Nucleus 13C
4
176.65
174.69
96.38
70.15
69.89
67.24
52.21
43.54
39.35
22.05
f1 (ppm)

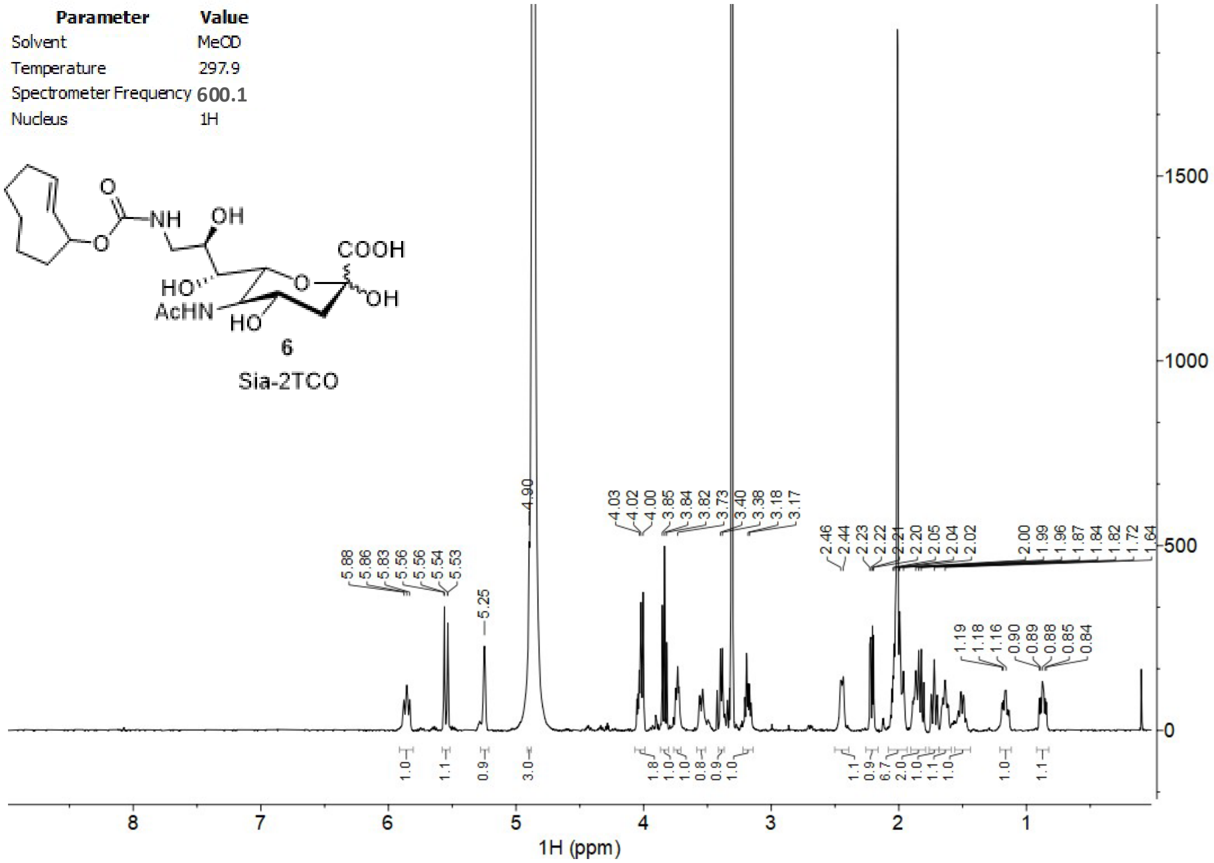
Parameter Value
Solvent MeOD
Temperature 297.9
Spectrometer Frequency 600.1
Nucleus 1H
COOH
NH
OH
O
HO
AcHN
HO
OH
6
Sia-2TCO
5.88 5.86 5.83 5.56 5.56 5.54 5.53
5.25
4.90
4.03 4.02 4.00 3.85 3.84 3.82 3.73 3.40 3.38 3.18 3.17
2.46 2.44 2.23 2.22 2.21 2.20 2.05 2.04 2.02
2.00 1.99 1.96 1.87 1.84 1.82 1.72 1.64
1.19 1.18 1.16 0.90 0.89 0.88 0.85 0.84
1.0 1.1 0.9 3.0 1.8 1.0 1.0 0.8 0.9 1.0 1.1 0.9 6.7 2.0 1.0 1.1 1.0 1.0 1.1
1500
1000
500
0
8 7 6 5 4 3 2 1
1H (ppm)
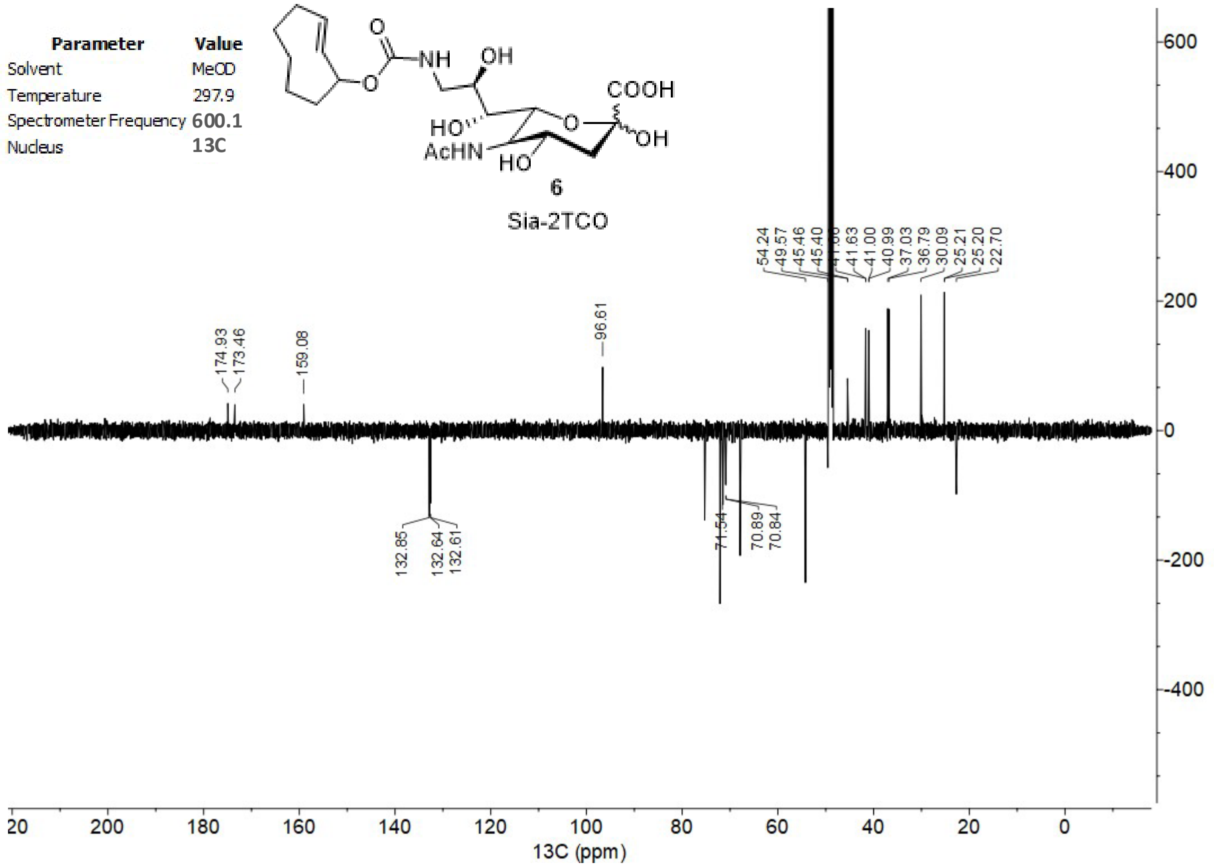
Parameter Value
Solvent MeOD
Temperature 297.9
Spectrometer Frequency 600.1
Nucleus 13C
COOH
NH
OH
O
HO
AcHN
HO
OH
6
Sia-2TCO
174.93 173.46
159.08
96.61
54.24 49.57 45.46 45.40 41.63 41.00 40.99 37.03 36.79 30.09 25.21 25.20 22.70
132.85 132.64 132.61
71.54
70.89 70.84
600
400
200
0
-200
-400
20 200 180 160 140 120 100 80 60 40 20 0
13C (ppm)

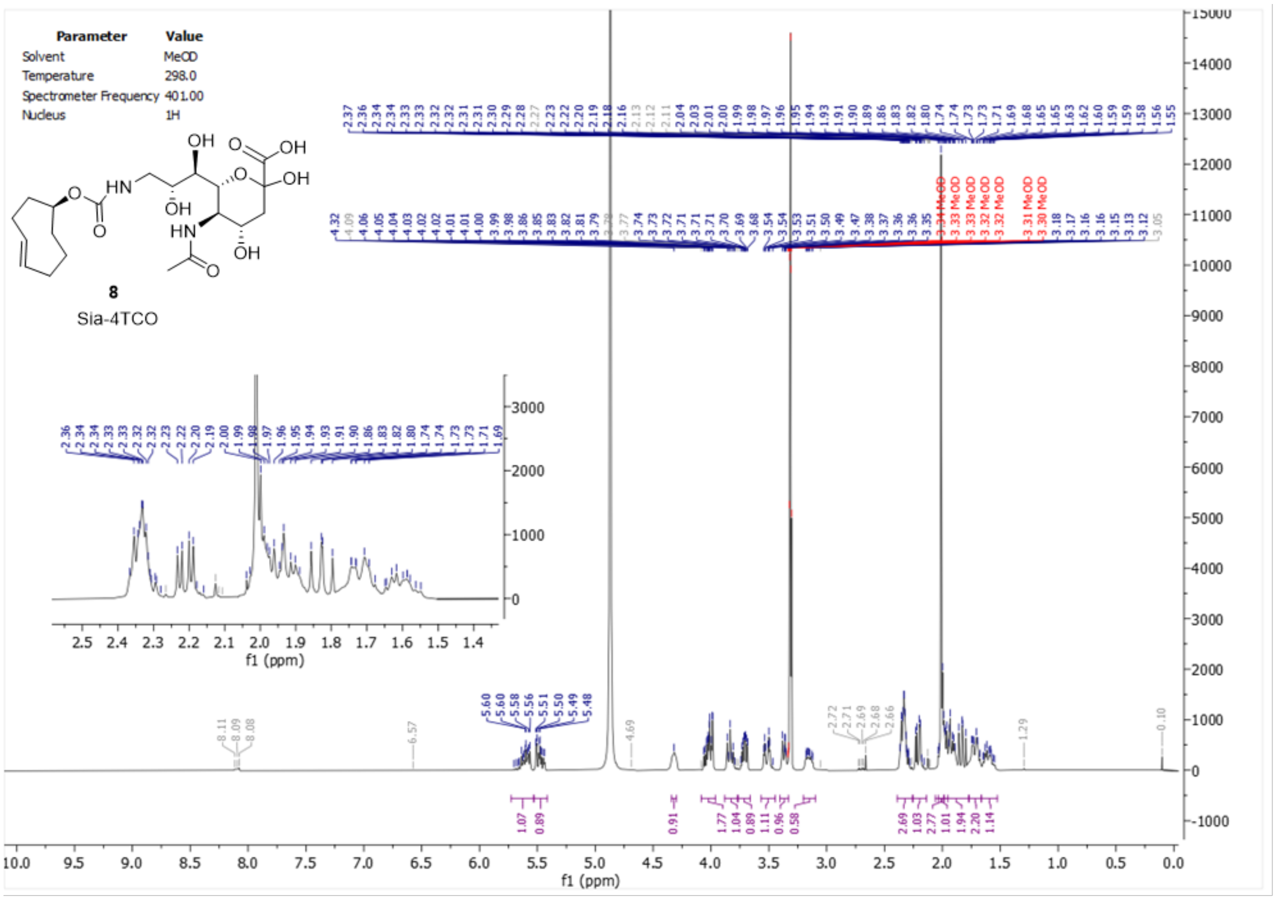

Parameter Value
Solvent MeOD
Temperature 298.0
Spectrometer Frequency 401.00
Nucleus 1H
8
Sia-4TCO
f1 (ppm)


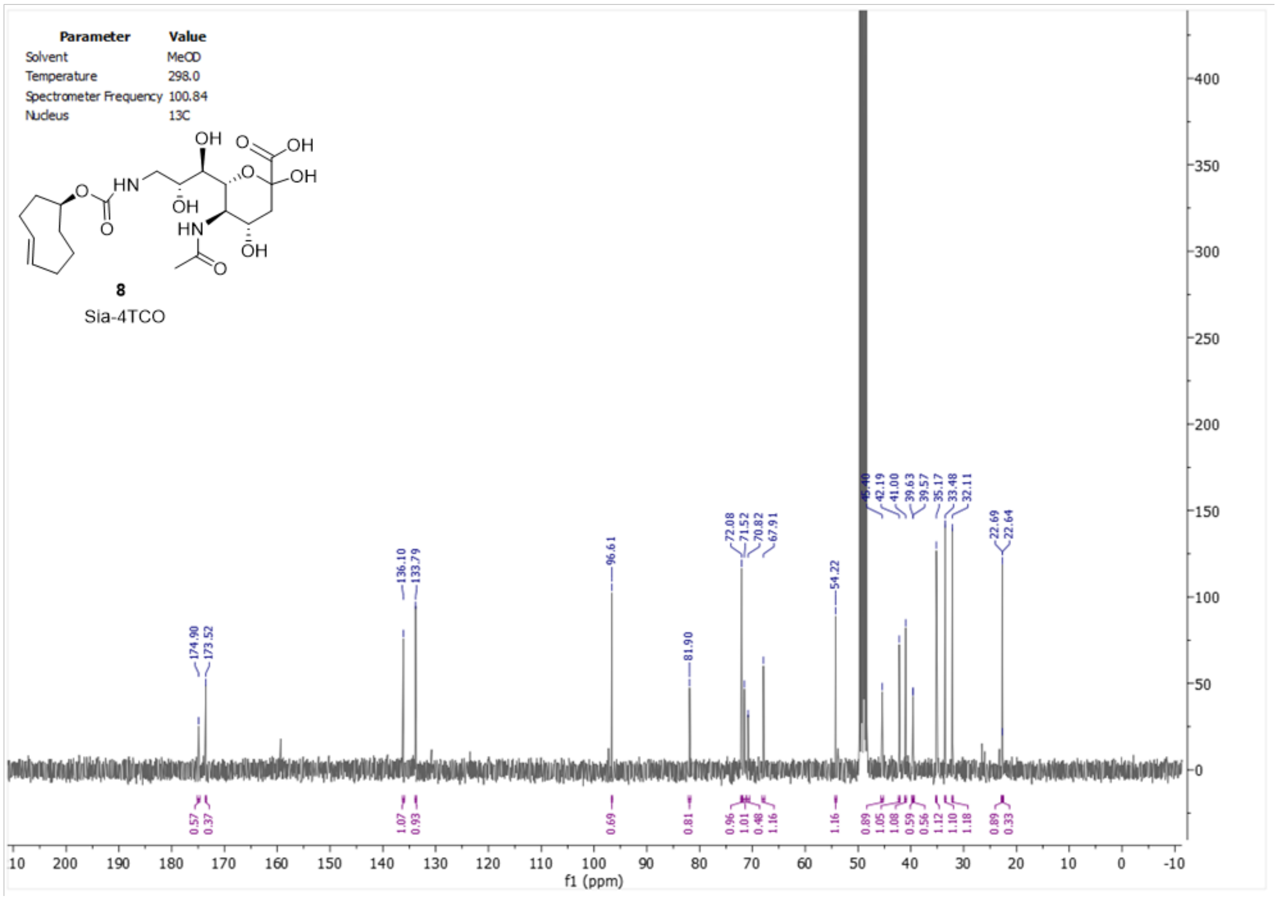

Parameter Value
Solvent MeOD
Temperature 298.0
Spectrometer Frequency 100.84
Nucleus 13C
8
Sia-4TCO
174.90
173.52
136.10
133.79
96.61
81.90
72.08
71.52
70.82
67.91
54.22
42.19
41.00
39.63
39.57
35.17
33.48
32.11
22.69
22.64
f1 (ppm)

| Parameter | Value |
|---|---|
| Solvent | MeOD |
| Temperature | 298.2 |
| Spectrometer Frequency | 499.98 |
| Nucleus | 1H |

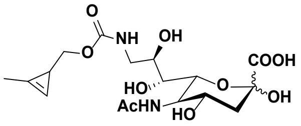

**Sia-Cp**
**10**

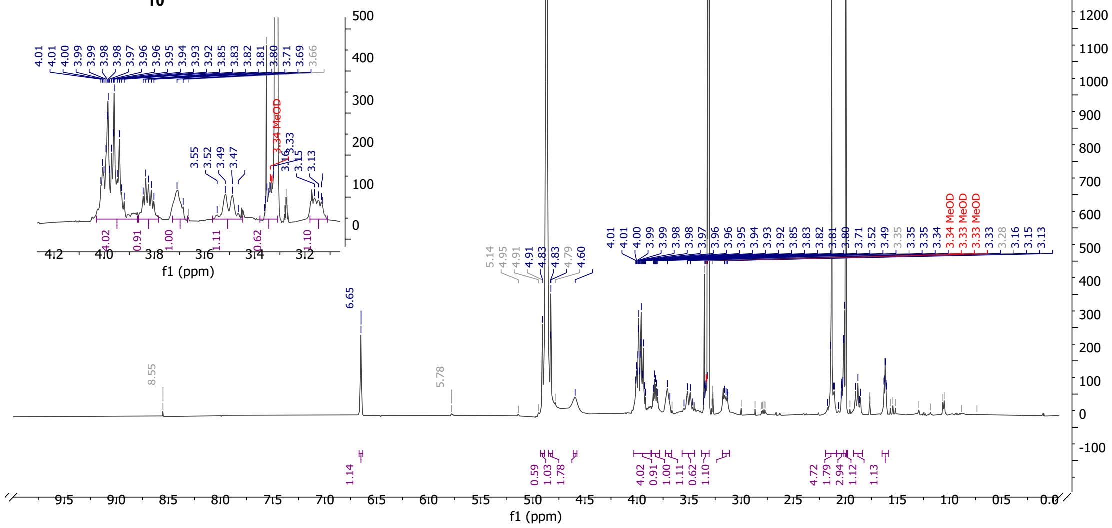


| Parameter | Value |
|---|---|
| Solvent | MeOD |
| Temperature | 298.2 |
| Spectrometer Frequency | 125.73 |
| Nucleus | 13C |

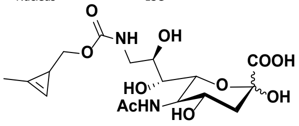

**Sia-Cp**
**10**

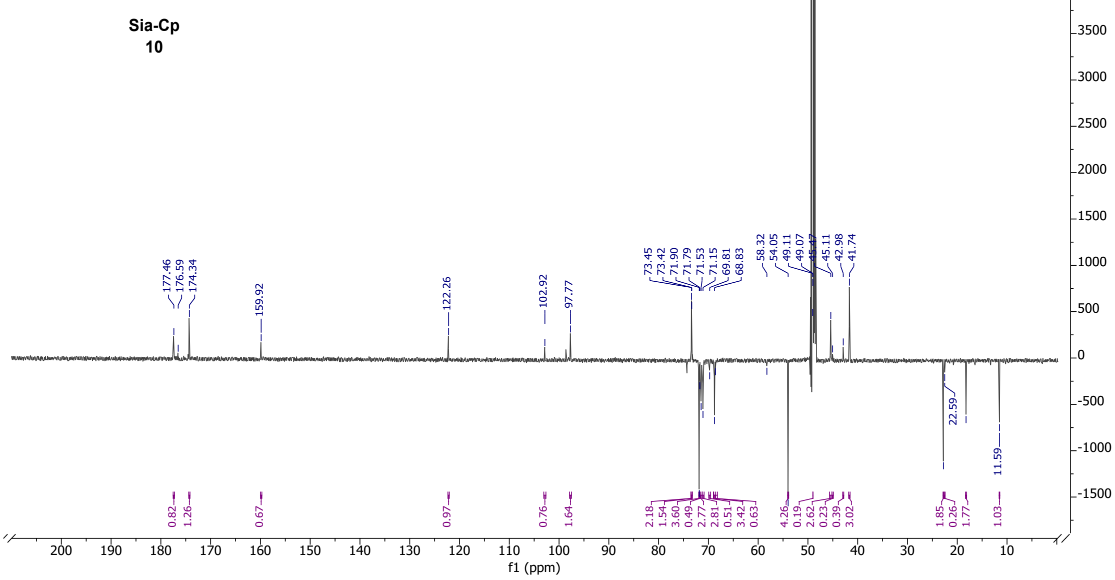

| Parameter | Value |
| --- | --- |
| Solvent | MeOD |
| Temperature | 298.0 |
| Spectrometer Frequency | 400.13 |
| Nucleus | 1H |

Sia-Norb
**12**

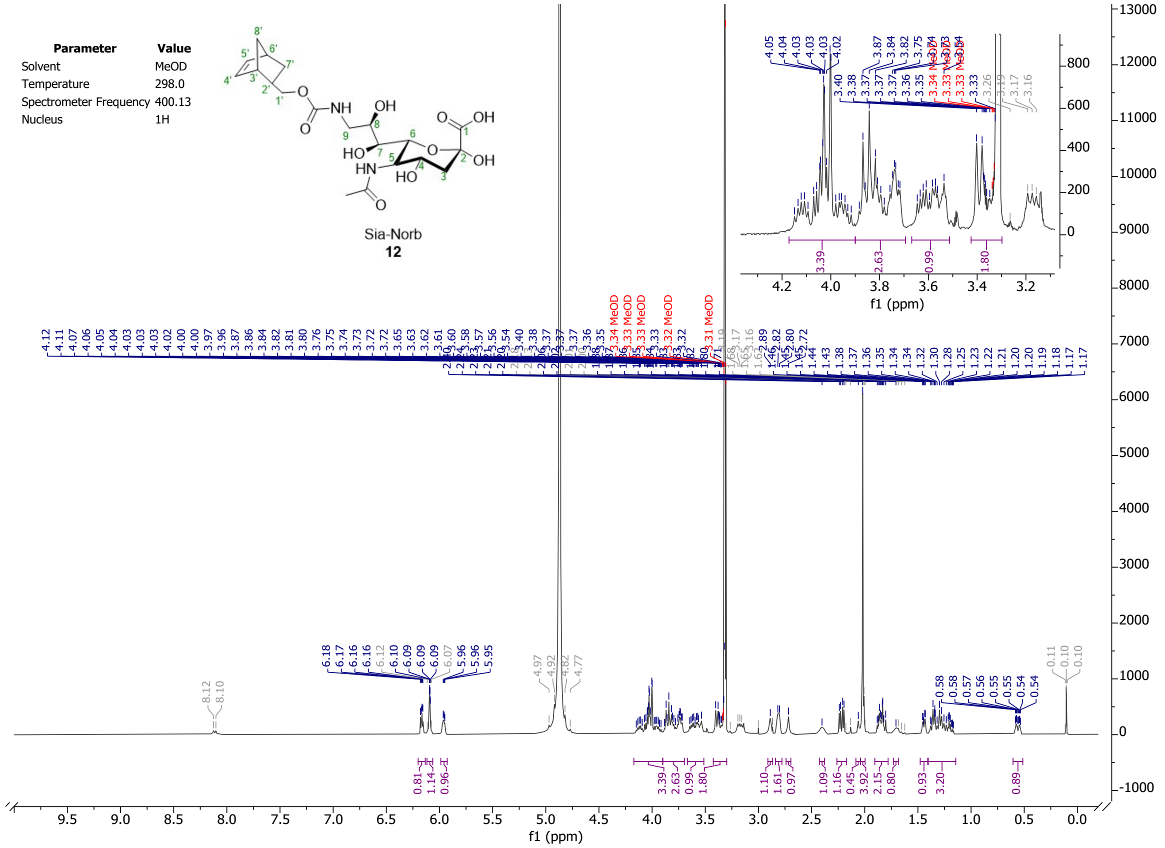


| Parameter | Value |
| --- | --- |
| Solvent | MeOD |
| Temperature | 298.0 |
| Spectrometer Frequency | 100.62 |
| Nucleus | 13C |

Sia-Norb
**12**

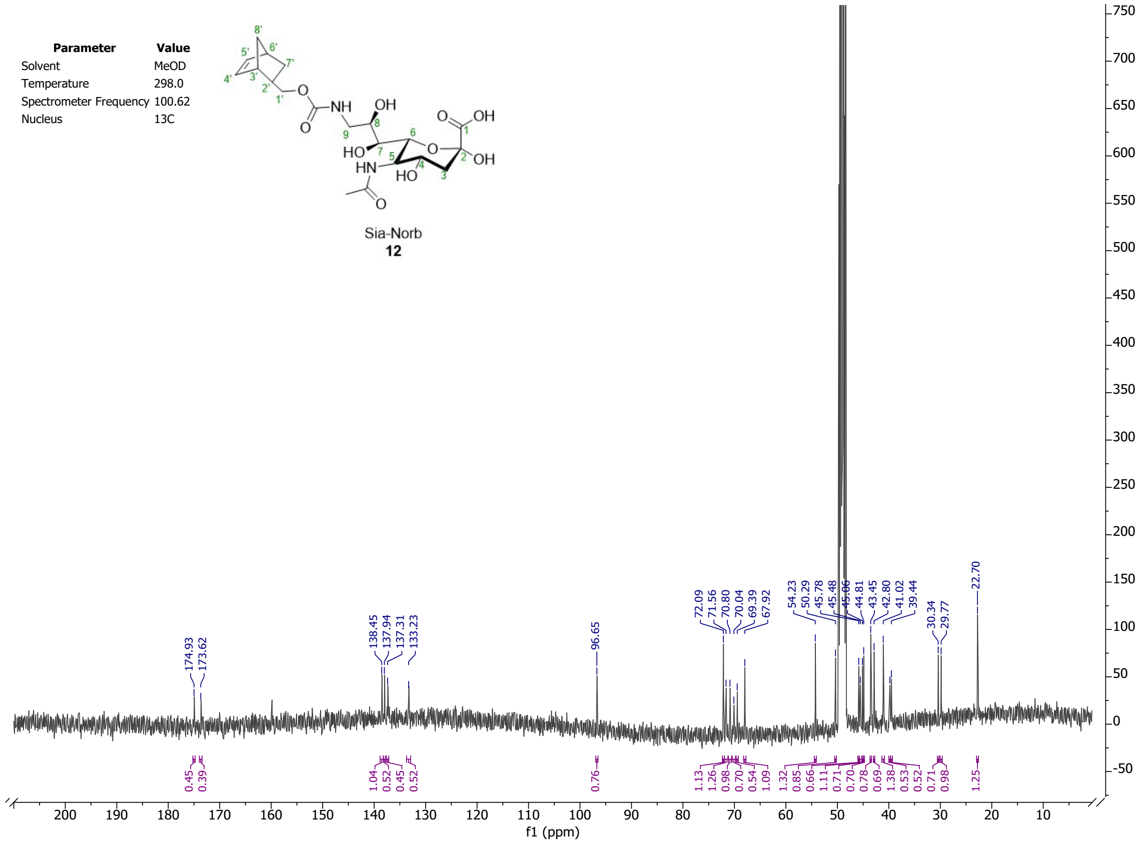

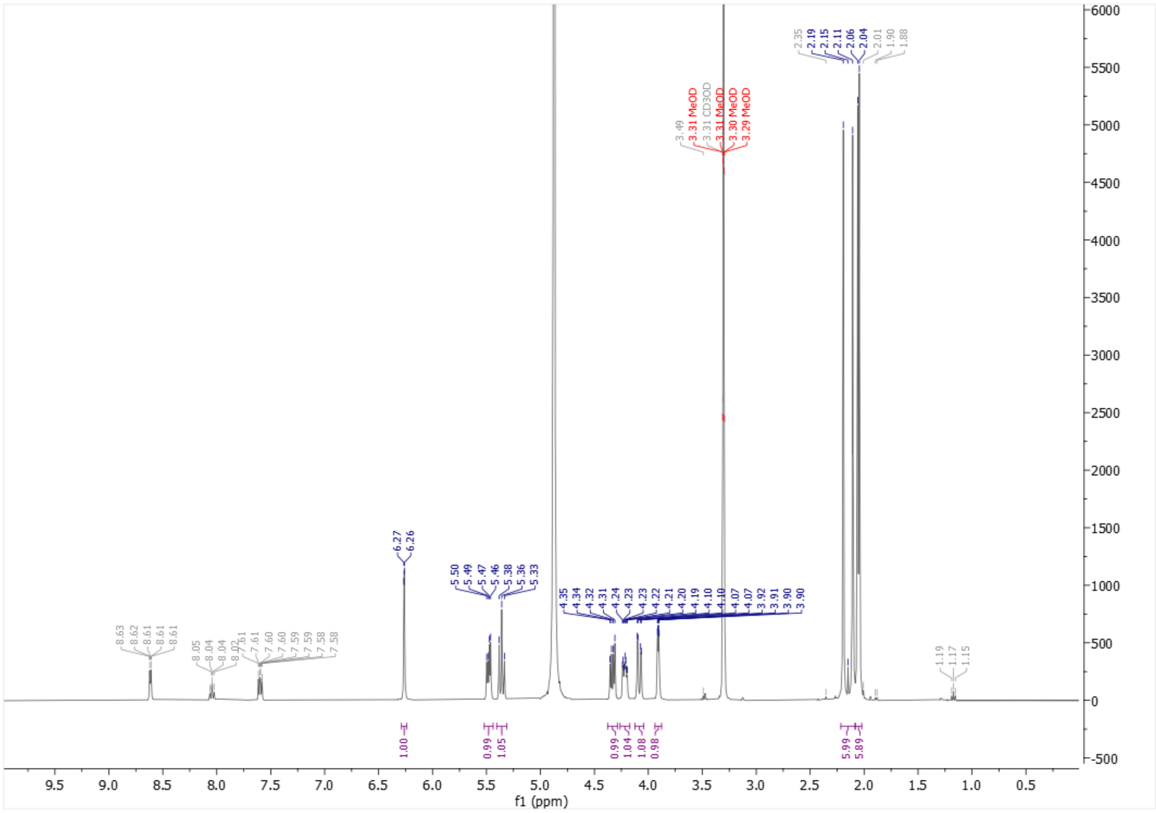

f1 (ppm)
3.31 MeOD
3.31 CD3OD
3.30 MeOD
3.29 MeOD


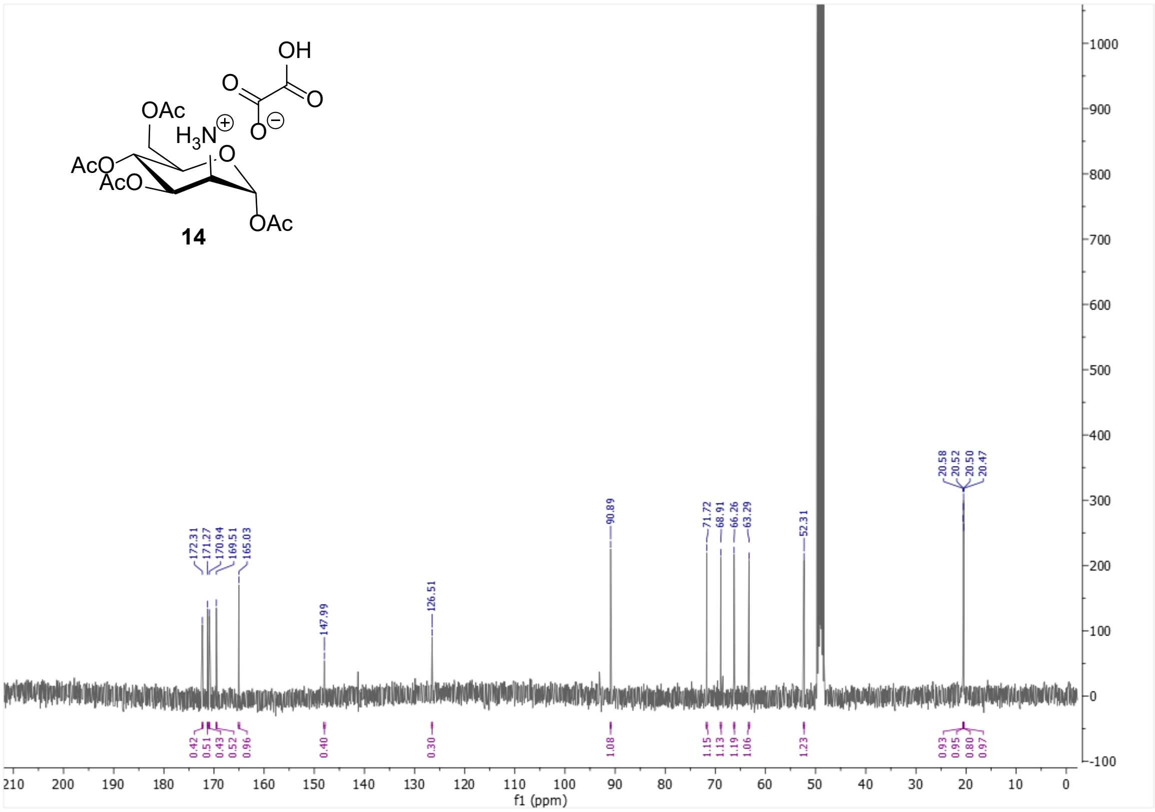

**14**


f1 (ppm)
172.31
171.27
170.94
169.51
165.03
147.99
126.51
90.89
71.72
68.91
66.26
63.29
52.31
20.58
20.52
20.50
20.47

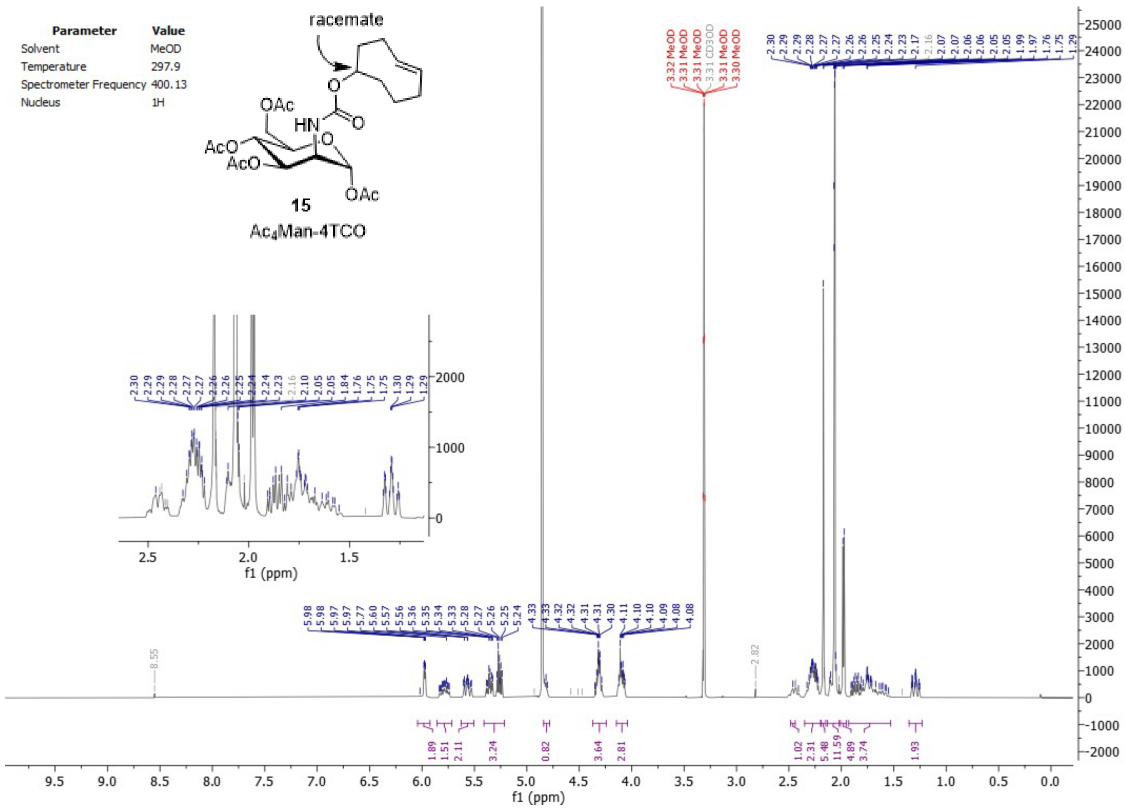
Parameter Value
Solvent MeOD
Temperature 297.9
Spectrometer Frequency 400.13
Nucleus 1H
racemate
OAc
HN
O
AcO
AcO
OAc
15
Ac4Man-4TCO
f1 (ppm)

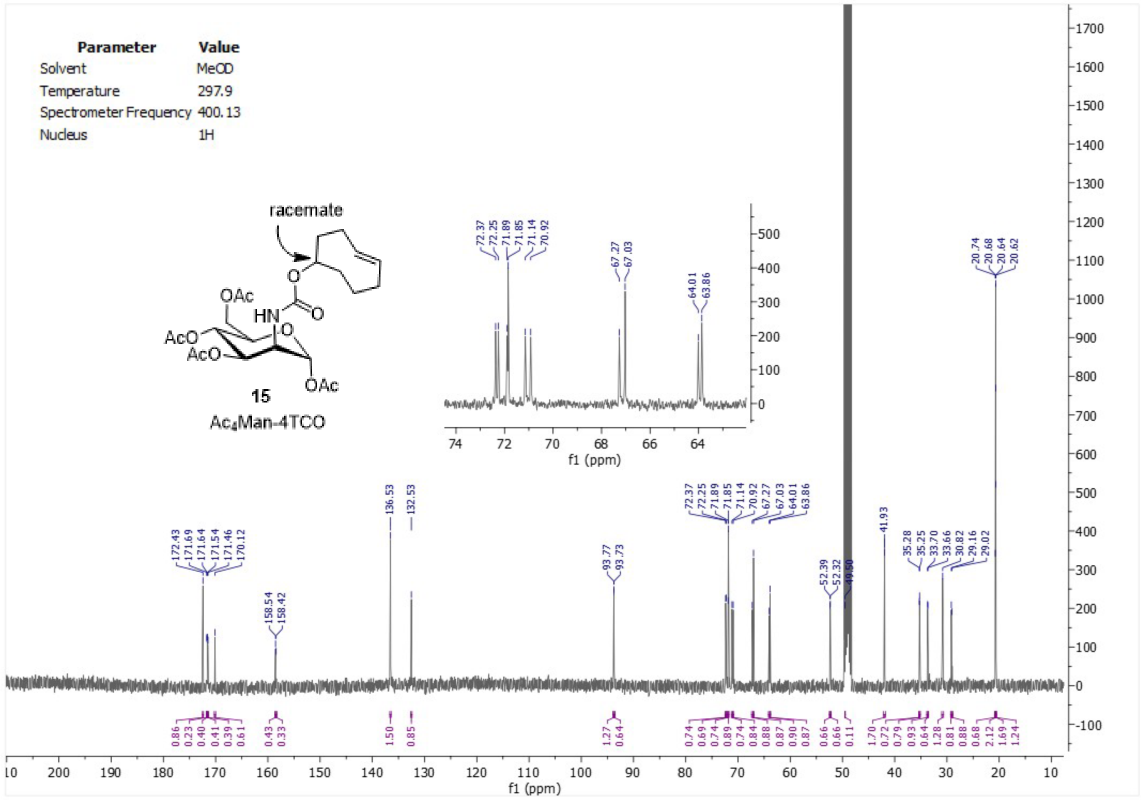
Parameter Value
Solvent MeOD
Temperature 297.9
Spectrometer Frequency 400.13
Nucleus 1H
racemate
OAc
HN
O
AcO
AcO
OAc
15
Ac4Man-4TCO
f1 (ppm)

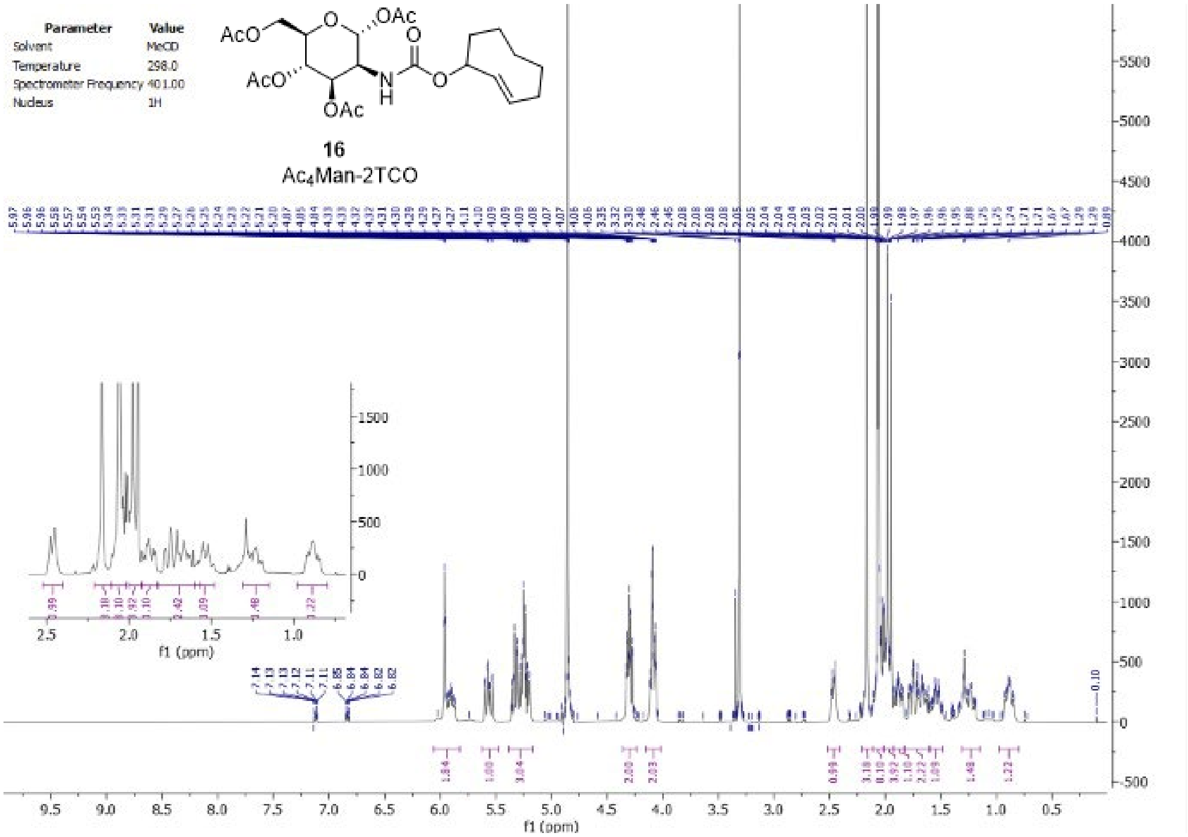
Parameter Value
Solvent MeOD
Temperature 298.0
Spectrometer Frequency 401.00
Nucleus 1H
16
Ac4Man-2TCO
f1 (ppm)

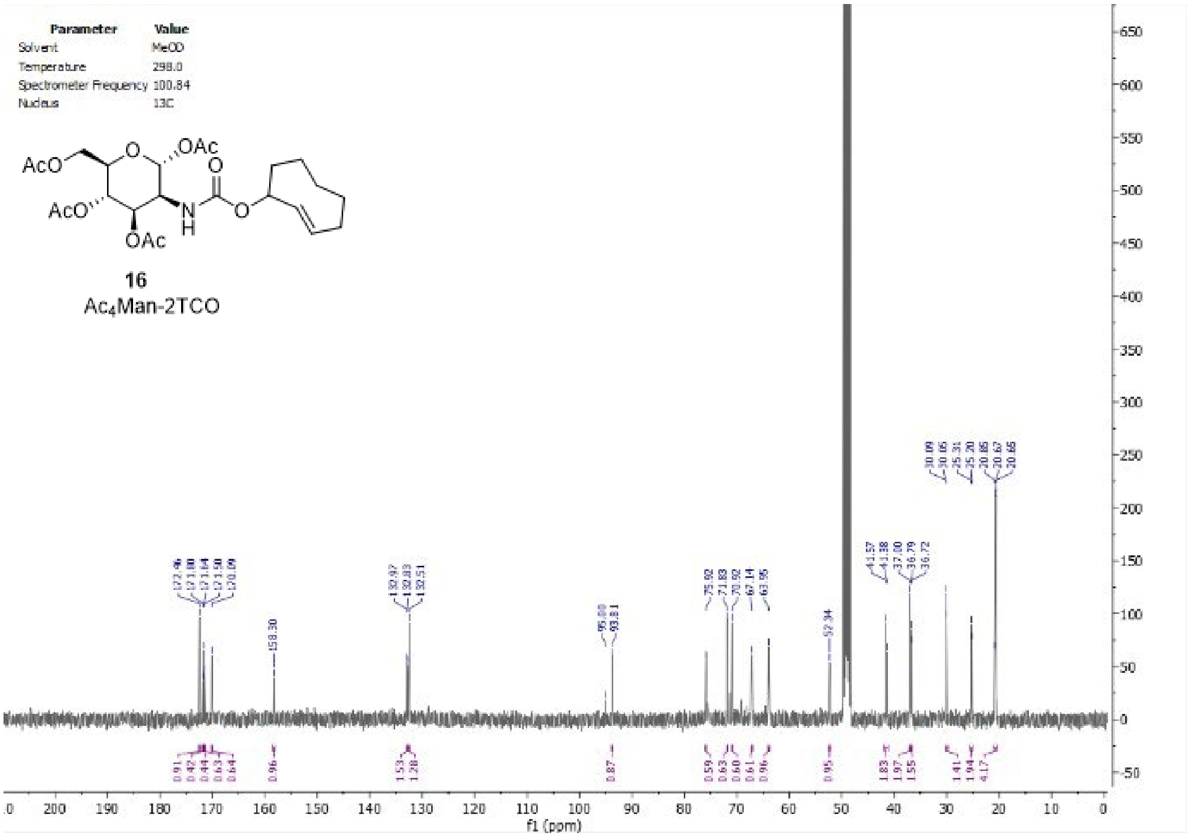
Parameter Value
Solvent MeOD
Temperature 298.0
Spectrometer Frequency 100.84
Nucleus 13C
16
Ac4Man-2TCO
f1 (ppm)

#### SulfoCy3-PEG3-Tz $^1$H NMR (401 MHz, DMSO-$d_6$)

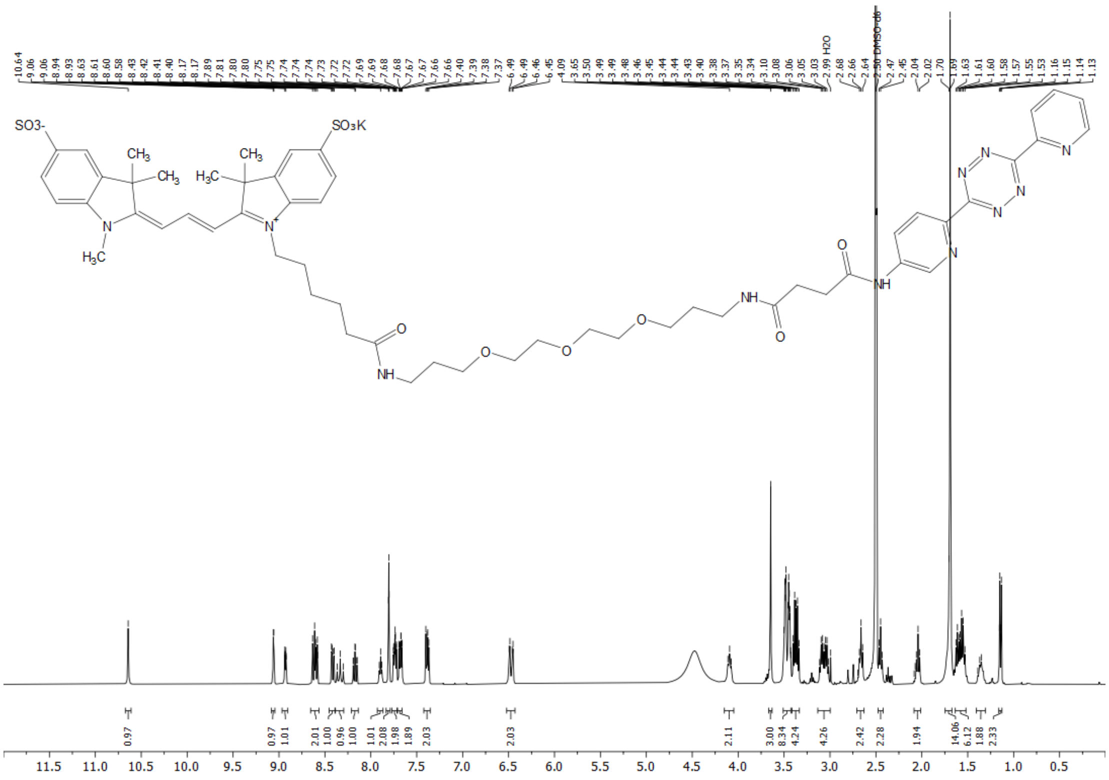


#### SulfoCy3-PEG3-Tz $^{13}$C NMR (101 MHz, DMSO-$d_6$)

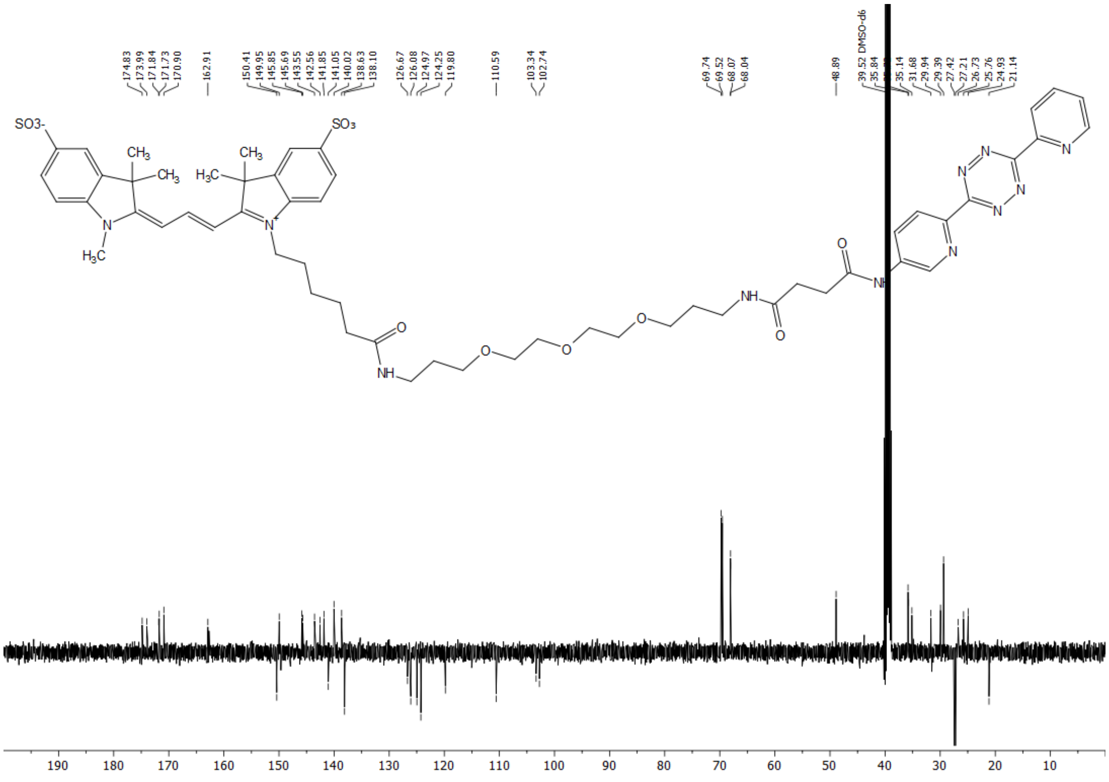

Possible alternative route via Chinese patent:

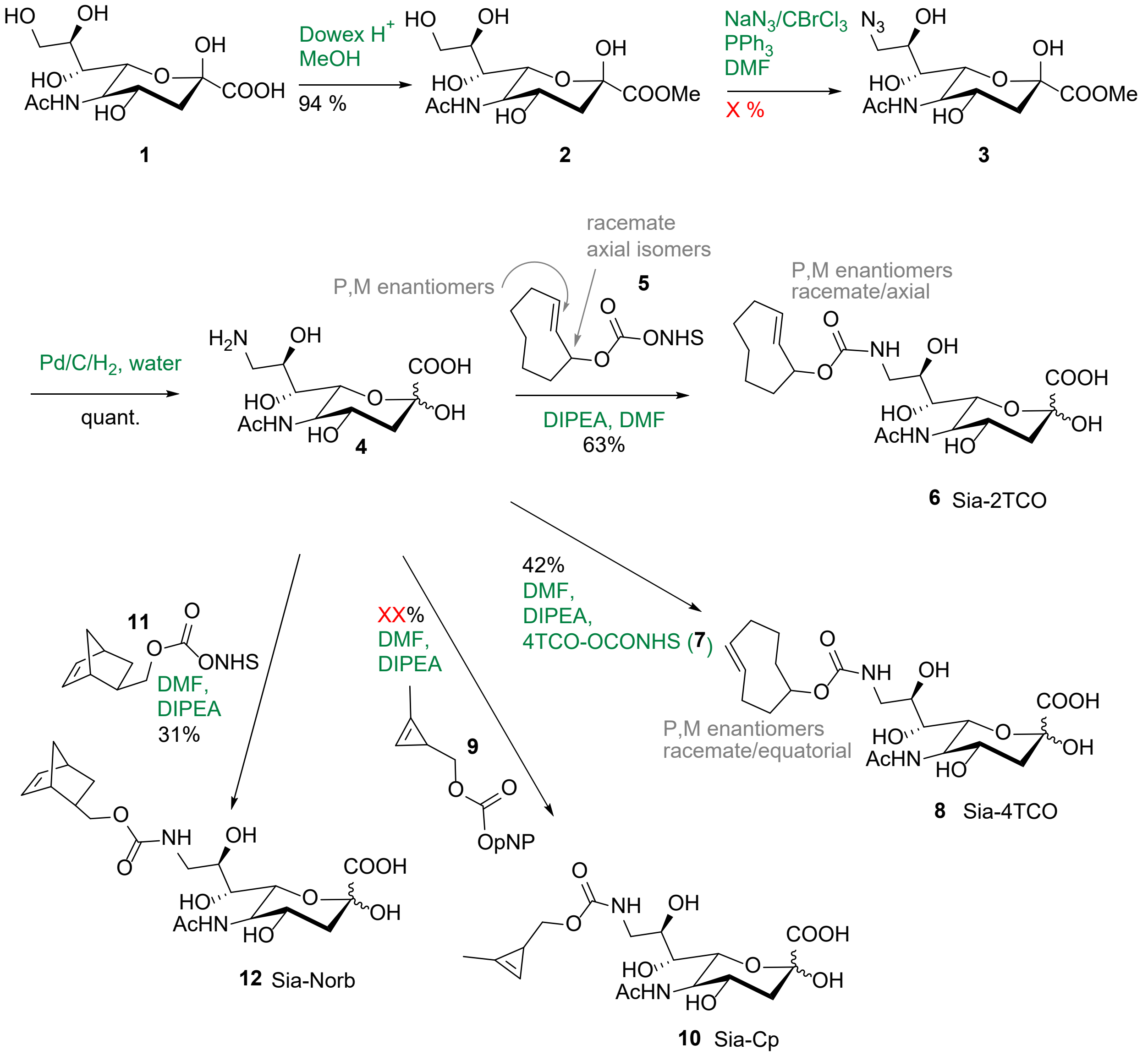


#### Methyl 5-Acetamido-9-azido-3,5,9-trideoxy-D-glycero-α-D-galacto-2-nonulopyranosylonate (X)

#### Methyl (E)-5-Acetamido-3,5-dideoxy-D-glycero-β-D-galacto-non-2-ulopyranosate (X)

Compound **X** was prepared as in lit.[10] (tenhle patent je opravdu podivny, musime se domluvit, jak ho ocitovat).

1.Freshly prepared Dowex50WX8 in $H^+$ cycle (4 g) was added to sialic acid (**1**; 15 g; 48.5 mmol) dissolved in dry MeOH (300 mL), and the mixture was stirred under an argon atmosphere for 48 hours. The resulting clear solution of the product was concentrated under vacuum to yield the methyl ester of sialic acid as an off-white solid. Crystallization (MeOH - EtOAc; 2 : 3) yielded the methyl ester of sialic acid, 14.7 g (94 %). Analytical data correspond to lit.[1]. $^1$H NMR (400 MHz, MeOD) δ = 1.90 (dd, *J*=12.9, 11.3, 1H, H-3a), 2.03 (s, 3H, NHAc), 2.23 (dd, *J*=12.9, 4.9, 1H, H-3b), 3.49 (dd, *J*=9.1, 1.5, 1H, H-7), 3.63 (dd, *J*=11.2, 5.7, 1H, H-9a), 3.71 (ddd, *J*=8.8, 5.7, 2.8, 1H, H-8), 3.79 (s, 3H, $OCH_3$), 3.77 – 3.88 (m, 2H, 10, H-5), 3.97 – 4.10 (m, 2H, H-6, H-4). $^{13}$C NMR (101 MHz, MeOD) 22.64 (NHCO*C*$H_3$), 40.70 (C-3), 53.14 (C-5), 54.33 ($OCH_3$), 64.83 (C-9), 67.85 (C-4), 70.19 (C-6), 71.64 (C-8), 72.09 (C-7), 96.68 (C-2), 171.77 (*C*OO$CH_3$), 175.12 (NH*C*O$CH_3$).

2. To the solution of sialic acid methyl ester (400 mg; 1.2 mmol), triphenylphosphine (714 mg; 2.7 mmol) and $NaN_3$ (804 mg; 12.4 mmol) in dry DMF (5.7 mL) at 0°C bromotrichloromethane (266 µL; 2.7 mmol) was added slowly. The mixture was stirred with exclusion of light 30 min at 0°C, then 16 hours at room temperature, and concentrated. Product **X** was purified by flash chromatography on column of silica, using $CH_3CN$, followed by C18 flash column chromatography (using water as the eluent). The **X** was isolated as a white solid ( mg; %). Analytical data in accordance with lit.[11]. $^1$H NMR (400 MHz, MeOD) δ = 1.85 – 1.96 (m, 1H, H-3a), 2.03 (s, 2H, NHAc), 2.22 (dd, *J*=12.9, 4.9, 1H, H-3b), 3.33 – 3.44 (m, 1H, H-9a), 3.40 – 3.48 (m, 1H, H-7), 3.48 – 3.62 (m, 1H, H-9b), 3.79 (s, 3H, $OCH_3$), 3.72 – 3.90 (m, 3H, H-8, H-5), 3.94 – 4.10 (m, 2H, H-6, H-4). $^{13}$C NMR (101 MHz, MeOD) δ = 22.62 (HNCO*C*$H_3$), 40.71 (C-3), 53.18 ($OCH_3$), 54.34 (C-5), 55.56 (C-9), 67.81 (C-4), 70.84 (C-8), 70.93 (C-7), 71.87 (C-6), 96.67 (C-2), 171.71 (*C*O$CH_3$), 175.17 (NH*C*O$CH_3$).
